# A multimodal perturbation atlas defines the phenotypic resolution of cellular morphology

**DOI:** 10.64898/2026.06.01.728087

**Authors:** Chad Liu, Alexander Hillsley, Madhurya Sekhar, Cassidy A. Jones, Gabriel Sturm, Taihei Fujimori, Diane M. Wiener, Karen W. Cheng, Talon Chandler, Alexander Lin, Duo Peng, Max Frank, Leah C. Dorman, Ilakkiyan Jeyakumar, Ivan E. Ivanov, Lin Luan, Greg Courville, Chris B. Charlton, Eduardo Hirata-Miyasaki, Sydney Ripsky, Ziwen Liu, Ritvik Vasan, Kyle I. Harrington, Kyle Awayan, Trang Le, Yiyun Rao, Giovanni Palla, Vincent Turon-Lagot, Miguel Cid-Rosas, Carolina Arias, Brian DeFelice, Josh Elias, Norma F. Neff, Alan R. Lowe, Shalin B. Mehta, Loïc A. Royer, Rafael Gómez-Sjöberg, Manuel D. Leonetti

## Abstract

Modeling cellular behavior requires measurements that capture how cells evolve across time, environments, and interventions. Microscopy is uniquely suited to this goal: it is non-destructive and can be applied to living cells in their native context. Yet its phenotypic resolving power remains incompletely characterized relative to molecular assays. Here, we present a multimodal perturbation atlas of 1,000 pooled CRISPR knockouts in A549 cells, profiled by fluorescence microscopy (42 live, 13 fixed markers), label-free quantitative phase imaging of the same live cells (at single timepoints), and single-cell RNA sequencing (scRNA-seq). We develop deep learning frameworks to interpret the rich cell-biological signatures in these ∼65M single-cell profiles. At matched reagent cost, phase imaging exceeds the phenotypic resolution of both fluorescence imaging and scRNA-seq, and more reliably recovers higher-order pathway organization. These results establish intrinsic morphology as a high-precision readout of cellular state, and lay a foundation for live-cell profiling of phenotypic trajectories.

## Introduction

Cell biology aims to understand the molecular circuits that govern cellular function. Genetic screens are central to this effort: by explicitly connecting defined perturbations to biological responses, they provide a causal window into cellular function that purely observational measurements cannot^1^. From this foundation, the ambition of modern AI-driven biology is to use screening data to construct predictive *in silico* models of cellular state—virtual cells and biological world models^2,3^. Building these models, however, places two demands on cellular phenotyping. First, model training will require data that prioritize the most informative measurements for a given task. Hence, the biological information content of different modalities must be quantitatively compared. Second, because cells are complex dynamical systems that constantly integrate signals about their internal state and environment^4^, models of cellular state will ultimately require perturbation data that capture not only fixed endpoints, but how cells evolve across time, context, and interventions^1–3,5^. Identifying scalable, live-cell-compatible modalities for the dynamic interrogation of cellular systems is therefore central to the next generation of functional genomics.

Single-cell RNA sequencing (scRNA-seq) has emerged as a defining measurement modality of the genomic era. A cell’s transcriptome captures tens of thousands of expression dimensions per cell, simultaneously encoding cell type, lineage, differentiation state, and signaling activity^6^. Pooled screening approaches such as Perturb-seq and CROP-seq combine CRISPR perturbations with single-cell transcriptomics at scale^7–9^. By directly linking perturbation to phenotypes, these pooled screens provide rich interventional data for mapping causal biological relationships, enabling the dissection of regulatory networks, genetic interactions, and heterogeneous cell states^10,11^. But scRNA-seq remains costly at scale, and is destructive: it observes only fixed endpoints. Computational trajectories inferred from splicing kinetics or optimal transport offer indirect readouts of dynamics^12–14^, but cannot substitute for direct, longitudinal observation of the same cell.

Cell morphology, by contrast, is one of biology’s oldest high-content phenotypes. Before molecular profiling became available, cellular and tissue architecture provided the basis for inferring biological states. In the 19^th^ century, Virchow placed morphology at the center of pathological reasoning and diagnosis^15^, on the principle that the functional properties of cells—intrinsically driven by chemical and physical processes—are largely reflected in their structure. Modern image-based screening methods have since transformed the exploration of cellular morphology into a scalable and quantitative strategy^16,17^. From multidimensional cytological fingerprints of the effect of drugs^18^ to the standardized Cell Painting assay^19,20^, high-content imaging has established that hundreds of cellular features (captured across nuclei, organelles, cytoskeleton, or signaling markers) can discriminate small-molecule mechanisms, infer gene function, and reveal variant-specific biology^21–26^. More recently, Optical Pooled Screening (OPS) has bridged the scalability of pooled CRISPR experiments with the power of cellular imaging, linking each perturbation to single-cell images^27,28^. Deployed across tens of millions of cells, OPS has shown that cellular morphology resolves co-functional cellular modules and enables the unbiased discovery of gene function^29–31^. In parallel, advances in representation learning have established image embeddings as a high-dimensional phenotypic space that tracks the molecular state of the cell ^32–37^.

A unique advantage of imaging is its non-destructive nature: it allows the same cells to be followed across time and observed in their environmental context, positioning imaging data as a key asset for training biological world models. Realizing this opportunity, however, will require a quantitative understanding of imaging’s phenotypic resolution, especially relative to molecular readouts. Two intertwined questions remain to be addressed. What is the phenotypic resolving power of live-cell imaging, particularly relative to scRNA-seq (the current reference for high-content cellular readout)? And how should imaging modalities be prioritized to maximize the biological information they extract? Little is known about how different imaging modalities and markers contribute to that information, and to what extent they are redundant with one another^22,38^. Even less explored is the information content of label-free imaging. Quantitative phase and bright-field imaging capture cell morphology without the need for exogenous stains or fluorescent reporters. Virtual staining can identify organelles from label-free data^39–41^, and phase signal alone can support classification of cell type, cell cycle, and disease states^41–43^ — indicating that the intrinsic optical properties of cells encode biologically meaningful variation. If label-free imaging can resolve cellular state on par with fluorescence stains or scRNA-seq, screening campaigns could be performed on native live cells, across time, at dramatically reduced cost and complexity.

Here we address these questions directly, and quantitatively define the phenotypic resolution of cellular morphology. We constructed a multimodal CRISPR perturbation atlas in which 1,000 gene knockouts were profiled in A549 lung adenocarcinoma cells, and developed a deep learning framework to explore the rich biology it uncovered. For each perturbation, we collected three orthogonal readouts of cellular phenotypes: *(i)* high-content fluorescence imaging across 42 live- and 13 fixed-cell markers, spanning organelles, signaling, and cytoskeleton, totaling ∼65M individual cells; *(ii)* label-free quantitative phase imaging of the same live cells (at single timepoints); and *(iii)* 0.6M single-cell transcriptomes via CROP-seq. This design allows us to directly compare phenotypic resolution (that is, the ability of each assay to distinguish individual perturbations) across these three core modalities and across the different imaging markers. Strikingly, provided enough cell coverage and at matched reagent cost, label-free phase imaging exceeds the phenotypic resolution of both fluorescence imaging and scRNA-seq across a broad range of perturbations. Our results establish intrinsic cell morphology as a high-precision measure of cellular state. They lay the foundation for live-cell perturbation atlases in which cellular trajectories can be measured dynamically, repeatedly, and in native biological context.

## Results

### A multimodal perturbation atlas

To quantitatively compare the biological information captured by different phenotypic readouts, we designed a multimodal screening strategy in which a single CRISPR perturbation library was profiled across multiple orthogonal assays (Figure 1A). We assembled a pooled knockout (KO) library targeting 1,000 genes, designed to span all major cellular pathways (Suppl. Fig. 1A, Suppl. Table 1). Essential processes such as gene expression, translation, and protein transport were deliberately over-represented because they yielded rich phenotypic signatures in previous screens^11,29,30^. Each gene was targeted by four sgRNAs, alongside 211 non-targeting sgRNAs as negative controls (Non-Targeting Controls, NTCs). As a cellular system, we chose A549 human lung adenocarcinoma cells, a line widely used in image-based phenotypic profiling^30,31,44,45^. Cas9 expression was placed under doxycycline-inducible control, and a 90-h induction window was chosen to balance the penetrance of perturbed phenotypes against the loss of cells carrying essential-gene knockouts^29^. sgRNAs were delivered by lentiviral transduction in a modified CROP-seq^8,46^ expression construct, compatible with both imaging and scRNA-seq readouts.

**Figure 1:**
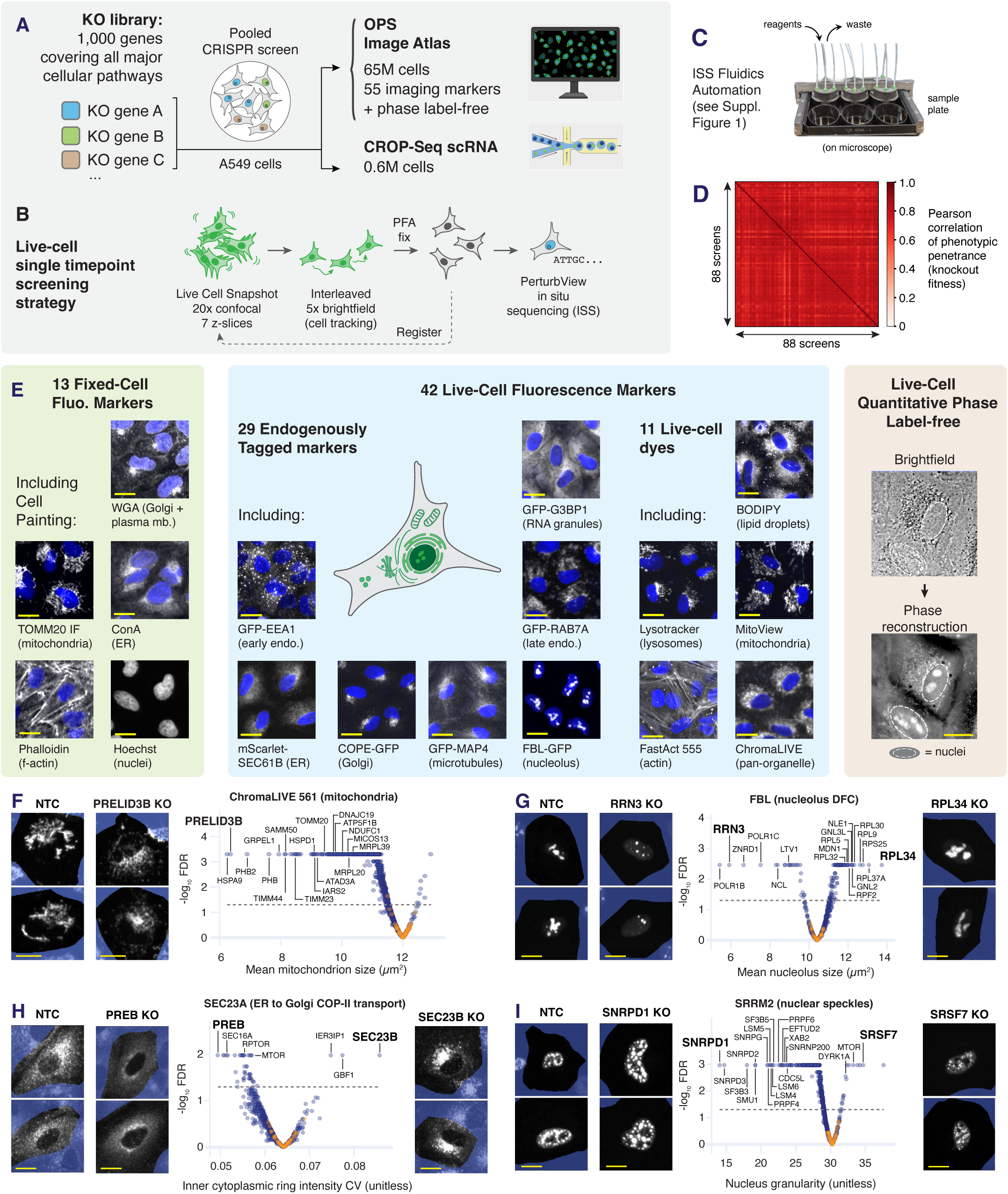
A multimodal pooled CRISPR screen of 1,000 gene knockouts. **(A)** Illustration of our multimodal pooled CRISPR screen strategy combining Optical Pooled Screens (OPS) and CROP-seq single-cell RNA (scRNA) sequencing. **(B)** Live-cell screening workflow. Live cells were imaged by 20× confocal microscopy across seven z-planes, with interleaved 5× brightfield imaging for cell tracking, before fixation and PerturbView *in situ* sequencing (ISS). **(C)** ISS chemistry was fully automated using a custom fluidics system integrated within an epifluorescence microscope. See also Suppl. Fig. 1B. **(D)** Correlation heatmap of phenotypic penetrance across 88 screens, using gene knockout fitness phenotypes. Fitness was quantified as the number of cells per sgRNA, normalized to cells with non-targeting controls. See also Suppl. Fig. 1D. **(E)** Our image atlas included 55 intracellular markers (42 imaged in live cells, 13 in fixed cells), capturing all major organelles and some key stress responses (see also Suppl. Figs. 2 and 3). Label-free quantitative phase imaging was performed in parallel on all live cell assays. For fluorescence images, marker signal is shown in grayscale, while nuclei are shown in blue. **(F) to (I)** Volcano plots validate expected biology in our screens. Each symbol: median measurement for a KO, calculated from up to 5,000 random cells sampled from different biological replicate experiments. Orange symbols: non-targeting controls, blue symbols: KO perturbations. (F) Mean area per mitochondrion decreases upon knockout of mitochondrial-associated genes. (G) Mean area per nucleolus decreases upon knockout of POLR1 (RNA Pol I) subunits, and increases upon knockout of ribosomal subunits. (H) Density of ER-to-Golgi transport vesicles is significantly altered when vesicular trafficking genes are perturbed. (I) Nuclear speckle granularity is significantly altered upon knockout of splicing factors. False discovery rate (FDR) was calculated against non-targeting control sgRNAs. Definition of the features being quantified and comprehensive plot values can be found in Suppl. Tables 12 and 13, respectively. All scale bars: 20 µm.

Comparing live and fixed phenotypes was central to our design; we therefore developed a live-cell-compatible OPS workflow (Figure 1B). High-content fluorescence phenotypes were acquired by confocal microscopy (20×, 0.55 NA, 7 z-slices spanning the whole cell). Live cells were imaged at single timepoints to balance the instrument time needed for acquisition, and individual screens were processed as batches of three wells in a 6-well plate (∼30 cm^2^), requiring ∼7 h to image the full surface area. Once live-cell acquisition was complete, the plate was fixed in 4% PFA, and sgRNA identity in each cell was decoded by PerturbView T7-based *in situ* sequencing (ISS)^46^. Because cells could move during the ∼7 h live-cell acquisition window, we regularly imaged the entire plate at low (5×) magnification with brightfield illumination to dynamically track nuclear positions and cell movement (Figure 1B). This tracking data was used to register individual cells between live-cell phenotyping and ISS images. To deploy this workflow at scale, we built an automated custom fluidics system integrated within an epifluorescence microscope to handle 4-color ISS chemistry steps^27^ (Figure 1C, Suppl. Fig. 1B). In its current iteration, this ISS instrument fully automates 10 sequencing cycles in <23 hours, without human intervention.

Together, this strategy enabled 88 separate screens spanning three orthogonal modalities: (*i*) fluorescence microscopy across 55 subcellular markers, imaged independently (including some repeats), (*ii*) label-free quantitative phase imaging of the same live cells (at single timepoints), and (*iii*) scRNA-seq via CROP-seq. The resulting atlas comprises image-based phenotypes from 64,918,233 perturbed cells, with an average coverage of 403 cells per sgRNA (1,610 cells per gene knockout) for fluorescence markers and 15,032 cells per sgRNA (60,126 cells per gene knockout) for quantitative phase imaging (Suppl. Fig. 1C). scRNA-seq phenotypes were measured in 606,075 cells. As a quantitative readout of perturbation penetrance in each screen, we measured the relative cell count of each knockout as a proxy for fitness phenotypes (Suppl. Fig. 1D). Tracking this metric across all imaging and scRNA-seq screens confirmed consistent penetrance and a high degree of reproducibility between experiments (Figure 1D, Suppl. Fig. 1D).

Image-based screens have historically focused on a small set of landmark organelle markers — most prominently those of the Cell Painting assay, selected to maximize compartment coverage with a compact panel of spectrally orthogonal dyes^19^. To provide a more comprehensive comparison of phenotypic resolution across markers, we designed our atlas around a deliberately diverse panel (Figure 1E). Thirteen markers were imaged in fixed cells, including a Cell Painting set as a benchmark of classical practice (Suppl. Fig. 2). To test the performance of live-cell screens, we further imaged 42 markers in live cells (as single snapshots), comprising 11 dyes, 29 endogenously tagged cellular landmarks, and two biosensors (Suppl. Fig. 3). Collectively, these markers cover all major organelles (from nuclear compartments to mitochondria to the plasma membrane) together with a few functional sensors for key stress responses including the unfolded protein response, caspase activity, and oxidative stress. In parallel, every live-cell screen was paired with label-free imaging. 2D quantitative phase was reconstructed from 3D brightfield data using the recently introduced WaveOrder algorithm^47^ (Suppl. Fig. 4A-C). This machine learning framework models the physics of microscopy optics to optimize phase fidelity across the large fields of view required at our screening scale.

Classical image-feature analysis confirmed that our screens recover expected biology, validating our protocol and highlighting the phenotypic richness captured in our multimodal perturbation atlas (Figures 1F-I). In a screen measuring the area of mitochondria, the highest-scoring phenotypes were strongly enriched for knockouts of mitochondrial proteins (Figure 1F). In a screen measuring the size of nucleoli, the site of ribosome biogenesis, top hits were enriched for ribosomal proteins and subunits of RNA polymerase I, which transcribes ribosomal RNA (rRNA; Figure 1G). The same coherence extended to less canonical sub-structures: a screen tracking COP-II ER-to-Golgi transport vesicles recovered components of the vesicular trafficking machinery (Figure 1H), while profiling the morphology of nuclear speckles (reservoirs of factors involved in transcription and pre-mRNA processing^48^) revealed hits enriched in splicing factors (Figure 1I). Further examples are presented in Suppl. Figure 5, underscoring the breadth of cell biology resolved by our dataset.

### Image embeddings reveal coherent programs of co-functional genes

To exploit the full phenotypic resolution captured by our image atlas, we built a computational workflow that transforms single-cell images into a high-dimensional embedding of the corresponding genetic perturbation (Figure 2A; Suppl. Fig. 6A). Image features were first aggregated at the level of individual sgRNAs, and then at the level of individual gene knockouts, yielding one vector per knockout that represents its phenotype across all imaged cells (Suppl. Fig. 6B). To quantitatively compare different embedding strategies, we adopted the mean average precision (mAP) framework^49^, a rank-based, data-driven information retrieval metric well suited to high-dimensional profiling data because it makes no assumption about feature distribution, linearity, or sample size. Concretely, mAP quantifies how reliably the embeddings of biologically related perturbations retrieve one another against a background of unrelated perturbations. mAP = 1 indicates that related perturbations are each other’s immediate nearest neighbors in embedding space, while mAP close to zero indicates that they are ranked no better than expected by chance. We applied mAP at two complementary scales of biological relatedness (Figures 2B, 2C). A “gene-knockout mAP” asks how similar embeddings from sgRNAs targeting the same gene are, and how well they are disambiguated from sgRNAs targeting other genes. Comparably, a “protein-complex mAP” asks how similar embeddings from different knocked-out subunits of the same complex are (complexes curated from the EBI Complex Portal^50^), and how well they are disambiguated from unrelated protein complexes. Both tasks rest on a strong biological prior: sgRNAs targeting the same gene, and knockouts targeting subunits of the same complex, should yield closely related phenotypes.

**Figure 2:**
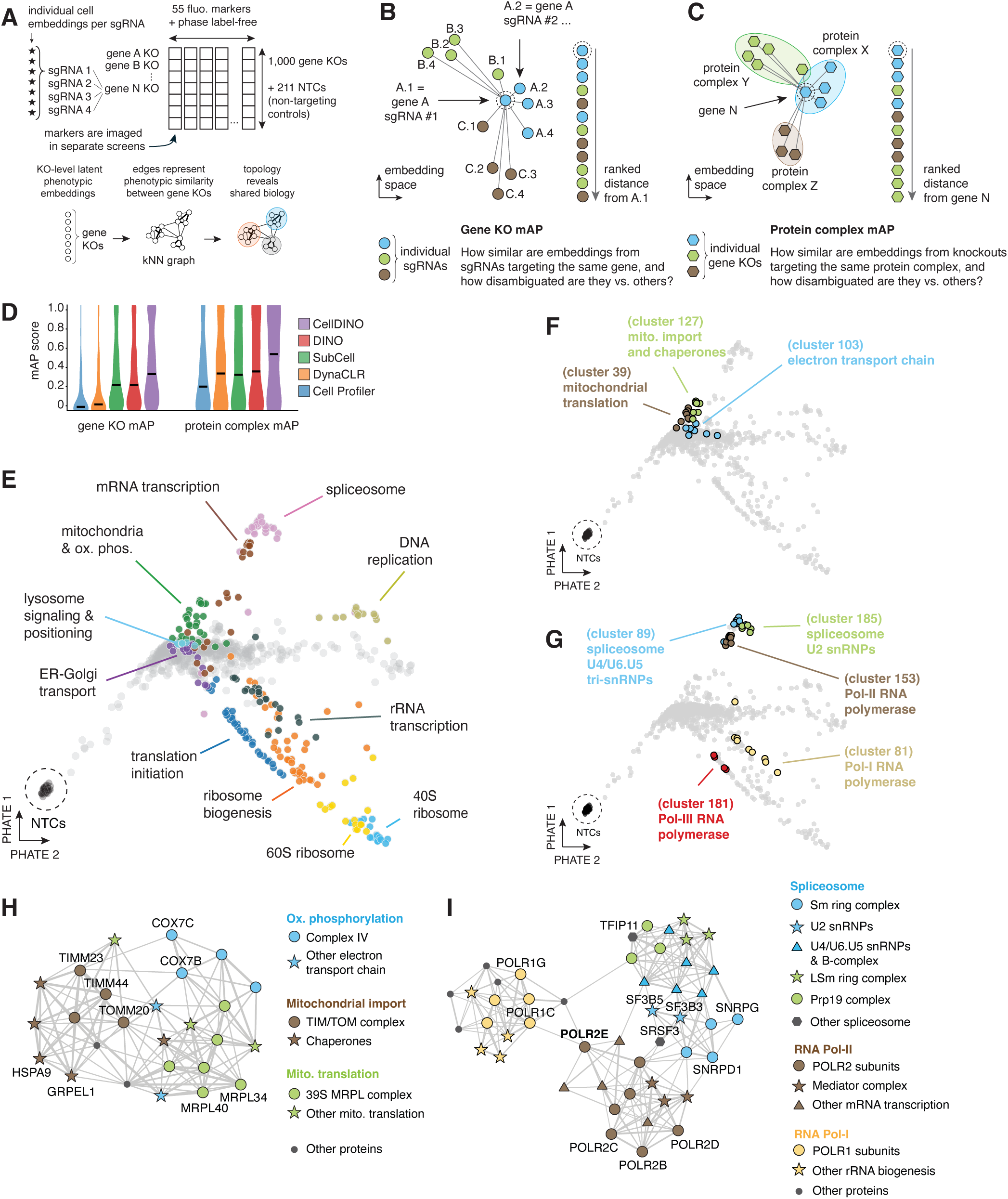
Cell-DINO image embeddings organize gene knockouts by shared biological function. **(A)** Image embedding workflow. For each marker, single-cell embeddings were aggregated at the sgRNA and gene-knockout levels. Embeddings obtained from different imaging markers were then concatenated to generate a multimodal representation for each perturbation. A k-nearest neighbor (k-NN) graph, constructed from gene knockout-level embeddings, captures phenotypic similarity between perturbations. **(B)** Illustration of gene-level mean average precision (mAP) metric. Gene-level mAP measures how reliably sgRNAs targeting the same gene can be disambiguated from unrelated perturbations. mAP values were calculated in the full-dimensional embedding space. **(C)** Same as (B). Protein-complex mAP measures how reliably knockouts targeting subunits of the same protein complex can be disambiguated from unrelated perturbations. **(D)** Benchmark of image representations across mAP retrieval metrics. Violin plots show distributions of gene-level and protein-complex mAP for CellProfiler, DynaCLR, SubCell, DINOv3, and Cell-DINO image embeddings. Horizontal lines indicate median mAP. **(E)** Two-dimensional PHATE visualization of Cell-DINO embeddings for 1,000 gene knockouts + 52 NTC “pseudo-genes” in which NTCs were pooled in groups of 4 to mimic a gene KO. Image embeddings were assembled using all 55 fluorescence markers and label-free phase imaging. Points are colored according to low-resolution Leiden clusters (resolution = 4), and clusters with clear biological enrichment are highlighted. See also data in Suppl. Table 10. **(F) and (G)** Same layout as (E), colored by high-resolution Leiden clusters (resolution = 29). Select clusters were manually annotated. See also data in Suppl. Table 10. **(H) and (I)** Topology of local k-NN graphs (n_neighbor = 10) from clusters highlighted in (F) and (G). Nodes connected with a single edge were pruned for clarity, and Pol III subunits are not shown. Fully annotated graphs are found in Suppl. Data S1.

We benchmarked five representations of our image data against this dual mAP criterion (Figure 2D). CellProfiler^51,52^, traditionally used in Cell Painting assays, describes each cell by a hand-engineered catalogue of shape, intensity, and texture features. The other four are deep learning-based approaches which, rather than predefining the features, learn them directly from the data: DynaCLR^53^, a contrastive-learning model which we re-trained on our dataset; SubCell, a microscopy foundation model trained on fluorescence images from the Human Protein Atlas^54^; DINOv3, a self-supervised vision transformer pre-trained on >1.7 billion web images^55^; and Cell-DINO, a DINO-based encoder pre-trained specifically on cellular images^56^. Rankings between the two mAP tasks were highly consistent, with all deep-learning representations outperforming classical featurization, and Cell-DINO scoring highest on both (Figure 2D). We therefore retained Cell-DINO as the image embedding solution for all subsequent analyses.

Pairwise correlation between gene-knockout embeddings provides a direct readout of phenotypic similarity^29^. We therefore also analyzed the full image dataset as a k-nearest-neighbor (k-NN) graph (Figure 2A), in which each node is a gene knockout and edges connect each gene to its k most-similar neighbors in phenotypic space — the same approach routinely used in scRNA-seq to delineate cell-type and cell-state communities^57,58^. Unsupervised Leiden clustering^59^ of this graph partitioned the perturbed list of genes into well-defined territories that align with major cellular processes, including translation, organellar transport, and oxidative phosphorylation (Figure 2E; visualized by PHATE^60^; data in Suppl. Table 10).

Higher-resolution clustering further resolved finer modules of co-functioning genes (Suppl. Table 10). Within the broad mitochondrial territory, distinct graph sub-clusters separated protein import and folding, the 39S mitochondrial ribosome, and respiratory-chain complexes (Figure 2F). Within the transcription territory, the three eukaryotic RNA polymerases — Pol I, Pol II, and Pol III — resolved into independent, internally coherent clusters that each recovered their expected subunit composition (Figure 2G).

Beyond clustering, the specific topology of the graph recapitulated known biology: functionally related processes occupy adjacent graph neighborhoods. Within the mitochondrial neighborhood (Figure 2H), subunits of the TIM/TOM translocase connected to their associated chaperones, while mitochondrial ribosomal subunits and electron transport chain complexes occupied distinct but adjacent positions. mRNA transcription and splicing likewise mapped to interconnected graph territories (Figure 2I): the LSm ring, the Prp19 complex, and the U2-U6 snRNPs formed separate splicing modules, all adjacent to the mRNA-transcription machinery (Pol II subunits, Mediator complex). Within this region, position tracked the temporal order of splicing itself^61^. Factors acting at the earliest steps (e.g. SRSF3, which nucleates E complex formation, and the U2 snRNP components SF3B3 and SF3B5, which license A complex assembly) sat closest to the Pol II territory, whereas late-acting factors such as Prp19 subunits and TFIP11, which promotes spliceosome disassembly, occupied the most distant positions (Figure 2I). Most strikingly, while the Pol I and Pol II clusters were otherwise well separated — reflecting their distinct roles in ribosomal- versus messenger-RNA transcription — POLR2E, a subunit shared by both RNA polymerase complexes, sat at the boundary between them, bridging the two clusters in precisely the position predicted by its shared biology (Figure 2I). Together, these observations establish that data-driven embeddings of our image atlas can resolve core cell biology and map co-functional genes, protein complexes and functional pathways.

### Deep-learning frameworks capture the cell biology that defines individual knockouts

Our perturbation atlas provides a large compendium of single-cell biological observations: for every gene knockout, our broad marker panel offers a direct link between the function of the perturbed gene and the biology read out by each marker. To facilitate the exploration and interpretation of our data across the many dimensions of markers and perturbations, we developed a deep learning framework to surface the cells whose phenotype most defines a given knockout. We trained a set-transformer^62^ classifier that takes as input a set of single-cell images from a given knockout (“bag”) and predicts which of the 1,001 perturbations (knockout or NTC) the bag represents (Figure 3A). We also trained the same architecture at the level of protein complexes, aggregating cells across shared subunits and asking the model to discriminate one complex from another. Given enough cells per bag, both gene-level and complex-level classifiers reached near-perfect accuracy when evaluated on held-out images, and performed equally well on the combined fluorescence panel and on phase imaging alone (Figure 3B; Suppl. Tables 16-17). This performance was enabled by the size of our atlas: training on an order of magnitude fewer cells produced a corresponding drop in validation accuracy (Figure 3C).

**Figure 3:**
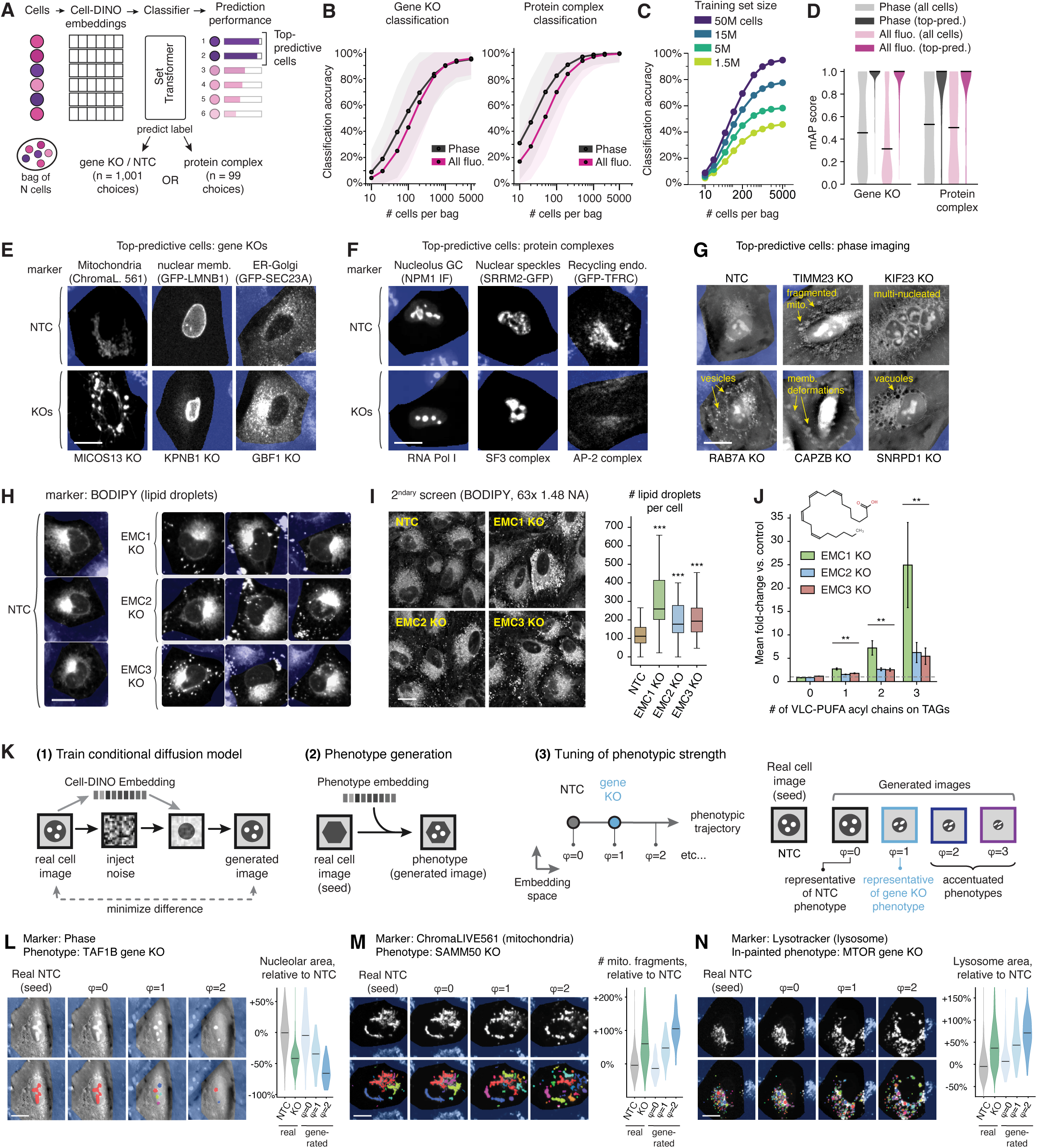
Deep-learning frameworks facilitate cell biological interpretation of phenotypes. **(A)** Classification-based selection of top-predictive cells. A set transformer takes a bag of N single-cell images from a given knockout and classifies either the knockout identity (n = 1,001 classes, including NTCs), or its assigned protein complex (n = 99 classes). Cells within each bag are weighted according to their contribution to the classification performance; the highest-weighted cells are retained as archetype examples of the prediction class. Per-gene accuracies for gene-KO and protein-complex classification at 100-cells bag size are presented in Suppl. Tables 16 and 17, respectively. **(B)** Top-1 classification accuracy as a function of bag size (10–5,000 cells per bag; log10 x-axis). Solid lines: median across classes; shaded bands: interquartile range (Q1–Q3). Left: gene-KO classifier (n = 1,001). Right: protein-complex classifier (n = 99). Two input modalities are compared: label-free 2D phase (black) and all combined fluorescent-marker channels (magenta). **(C)** Top-1 classification accuracy from phase image embeddings, as a function of training set size. **(D)** mAP score distributions calculated from all available cells, or top-predictive cells only. Horizontal bars display median value. **(E)** Top-predictive cells for representative gene knockouts (fluorescence). Each column shows the top-predictive cells for both NTC (top) and gene knockout condition (bottom), showing the marker channel most predictive for that knockout: MICOS13 (marker: mitochondria, ChromaLIVE 561), KPNB1 (marker: nuclear membrane, GFP-LMNB1), GBF1 (marker: ER–Golgi, GFP-SEC23A). (F) Same as (E), but showing top-predictive knockout cells for protein complexes: RNA Pol I complex (marker: nucleolus, NPM1 IF), SF3b complex (marker: nuclear speckles, SRRM2-GFP), AP-2 complex (marker: recycling endosomes, GFP-TFRC). (G) Top-predictive cells in the label-free 2D phase modality. (H) Top-predictive cells reveal upregulation of BODIPY signal (a lipid droplet dye) as a defining feature of EMC subunit knockouts. (I) EMC KO phenotype was validated in an arrayed secondary screen, in which knockouts were processed individually. BODIPY-stained cells were imaged at high magnification (63× confocal microscopy, 1.47 NA objective) to resolve individual lipid droplets. Image quantification (right inset) reveals increased lipid droplet counts in knockout cells (n ≥ 55 cells per condition). Boxes represent 25th, 50th, and 75th percentiles; whiskers represent 1.5 times the interquartile range. *** p < 10^-4^ (Mann-Whitney U test, two-sided). (J) Lipid remodeling in EMC KO cells. Quantification of lipid species by mass-spectrometry lipidomics revealed a significant increase in triacylglycerides (TAGs, the neutral lipid component of lipid droplets) containing VLC-PUFA chains (VLC-PUFA: very long-chain poly-unsaturated fatty acids). Barplots show mean fold-change of mass spectrometry intensities of lipid species between KO and NTC control cells. Error bars show s.e.m. from n = 3 replicates for EMC1 and EMC3, and n = 2 for EMC2. ** p < 10^-3^ (Welch t-test against control). (K) Principle of image generation with a conditional diffusion model (see text for details). (L) **, (M)** and **(N)** Image generation across a range of phenotypic strength. Real images (left panels) are used as seeds, and the diffusion model was used to generate images representing the effect of a KO perturbation on that same cell. Perturbation strength can be tuned by the scaling factor φ, allowing for the generation of accentuated phenotypes which can expose phenotypic signatures. Bottom rows show segmentation masks. Violin plots: quantification of specified image features across n ≥ 100 cells. Horizontal bars display median values. Scale bars = 20 µm.

To extract interpretable phenotypes from these classifiers, we leveraged a removal-based explainability strategy^63^, scoring each cell by how much its inclusion in the bag raises the classifier’s probability of predicting the correct class (see Materials and Methods). The highest-scoring cells for a given knockout or complex, which we term “top-predictive cells”, are those the model relies on most, and therefore act as phenotypic representatives that provide a natural entry point for biological interpretation. Validating this approach, gene-knockout or protein-complex mAP scores were significantly higher when computed on top-predictive cells than on all cells in the dataset (Figure 3D).

At the level of individual knockouts, the top-predictive cells consistently surfaced the markers most diagnostic for the perturbed gene’s biology (Figure 3E). For MICOS13, which organizes the mitochondrial inner membrane and its cristae, the ChromaLIVE 561 dye revealed mitochondrial hypertrophy. For KPNB1 (a major nuclear import receptor), the lamina marker GFP-LMNB1 showed a condensation of the nuclear membrane. GBF1 offered a more subtle example: its top-predictive cells came from imaging GFP-SEC23A, a component of the COPII ER-to-Golgi vesicular coat, whose signal was markedly elevated upon knockout. This observation is consistent with the canonical role of GBF1 as the Arf1 guanine-nucleotide exchange factor that maintains Golgi architecture^64^, since loss of Arf1 activation collapses the Golgi and expands the ER exit sites at which COPII components accumulate.

The classifier trained at the level of protein complexes achieved similar outcomes (Figure 3F). Top-predictive cells for knockout of the RNA Polymerase I complex revealed rounded and smaller nucleoli from NPM1 staining, a sign of disrupted rRNA transcription^65^. For the SF3b complex, a core module of the spliceosomal U2 snRNP, top-predictive cells displayed an agglomeration of SRRM2-GFP signal, consistent with the enlargement of nuclear speckles that follows splicing inhibition^66^. And for the AP-2 complex, a key clathrin adaptor for vesicular endocytosis, they showed a decrease in the recycling-endosome pool of GFP-TFRC, as expected when internalization of its transferrin-receptor cargo is blocked.

Top-predictive cells from phase imaging recovered comparably interpretable phenotypes without any fluorescent label (Figure 3G): mitochondrial fragmentation upon knockout of TIMM23, a core component of the mitochondrial import channel; multinucleated cells upon knockout of KIF23, a motor protein essential for cytokinesis; accumulation of intracellular vesicles upon knockout of RAB7A, a regulator of endosomal maturation; cell elongation and membrane deformation upon knockout of the actin-binding protein CAPZB; and prominent cytoplasmic vacuoles upon knockout of the core spliceosomal Sm protein SNRPD1, consistent with the autophagic vacuolization reported upon Sm depletion^67^.

Across markers, complexes, and label-free images, these phenotypes consistently recapitulated established gene function. But top-predictive cells also surfaced phenotypes that point to less-characterized biology. The ER membrane protein complex (EMC) is a multi-subunit insertase and chaperone for membrane proteins^68^. Its top-predictive cells showed an increased vesicular signal in BODIPY, a stain for lipid droplets, which was highly consistent across the three EMC subunits in our perturbation set (EMC1/2/3, Figure 3H). We validated this phenotype in an arrayed secondary screen in which each subunit was knocked out separately and imaged at higher resolution, revealing a significant increase in lipid droplet number (Figure 3I). Because lipid droplets store neutral lipids, primarily triacylglycerides (TAGs)^69^, we asked whether EMC loss also remodels lipid droplet composition. Mass-spectrometry lipidomics of knockout cells indeed revealed a significant up-regulation of TAGs containing very long-chain polyunsaturated fatty acids (VLC-PUFAs, Figure 3J). Intriguingly, hepatocyte-specific *Emc3* ablation was recently reported to decrease lipid droplet content in the mouse liver^70^, pointing to an interplay between EMC function and lipid droplet homeostasis that might vary across tissues. Dissecting this mechanistic link is beyond the scope of the present work, but this example illustrates how top-predictive cells can generate hypotheses for mechanistic follow-up.

The examples above involved strong, readily recognizable phenotypic signatures. In many other cases, the phenotypes of top-predictive cells were subtler and harder to interpret. To address this challenge, we developed a second deep-learning framework based on a diffusion image-generation model^71^. Large amounts of noise were injected into real cell images, and the model was conditioned on the Cell-DINO embedding of the starting cell to reconstruct a de-noised image (Figure 3K, panel 1). Minimizing the loss between real and generated images teaches the model to generate cell images representative of a given embedding, which can then be used to induce phenotypes *in silico*: starting from a real cell image as a seed, the effect of a perturbation can be applied to that same cell (Figure 3K, panel 2). This generative strategy confers two separate advantages. First, it removes the confounding effect of cell-to-cell morphological heterogeneity, which otherwise masks phenotypic differences between perturbations. In our model, the seed image was designed to act as a background that holds the overall cell morphology fixed, making those differences far easier to recognize. Second, this strategy allows phenotypes to be accentuated (Figure 3K, panel 3). In embedding space, the vector connecting NTC and gene-knockout embeddings defines a phenotypic trajectory, along which we define phenotype strength by a factor φ (akin to a reaction coordinate in phenotypic space). Whereas φ = 1 corresponds to the measured knockout phenotype, φ > 1 generates images representing more accentuated signatures along the same trajectory.

Figures 3L–N illustrate how image generation can aid the interpretation of the signatures that define different perturbations. Knockout of TAF1B, which targets RNA Pol I to rDNA, validated the approach: consistent with the known effect of disrupting rRNA transcription^65^, the generated images displayed a decrease in nucleolar area upon perturbation that became more pronounced upon accentuation (Figure 3L). We then used the image generation framework to interpret phenotypic signatures that were harder to identify. Knockout of SAMM50, a core component of the mitochondrial sorting and assembly machinery, led to mitochondrial fragmentation that was mildly detectable at φ = 1, but readily apparent in the accentuated images (Figure 3M). Similarly, accentuated images underscored an increase in lysosomal size upon MTOR knockout, consistent with the central role of MTOR in lysosomal homeostasis^72^ (Figure 3N).

Together, the top-predictive-cell and image-generation frameworks capture perturbation biology in a form that is both interpretable and quantitative, providing a scalable, data-driven entry point for the phenotypic dissection of any perturbation in our dataset. To make both accessible, we built OPSin, the <u>O</u>ptical <u>P</u>ooled <u>S</u>creening <u>in</u>terpretability viewer, available at opsin.apps.czbiohub.org.

### Phase imaging exceeds the phenotypic resolution of fluorescence markers given sufficient cell coverage

We next dissected the phenotypic resolution contributed by each individual imaging marker; that is, the ability of a single marker to identify and disambiguate biological relationships among gene knockouts. Using Cell-DINO embeddings and the mAP framework introduced above, we computed gene-level and protein-complex mAP scores for each channel of our atlas separately. Marker performance spanned a broad range and was uniformly lower than the score obtained when all channels were combined (Figure 4A, Suppl. Tables 7 and 8). Most strikingly, when all available cells were used, label-free quantitative phase imaging outperformed every fluorescence channel on both retrieval tasks.

**Figure 4:**
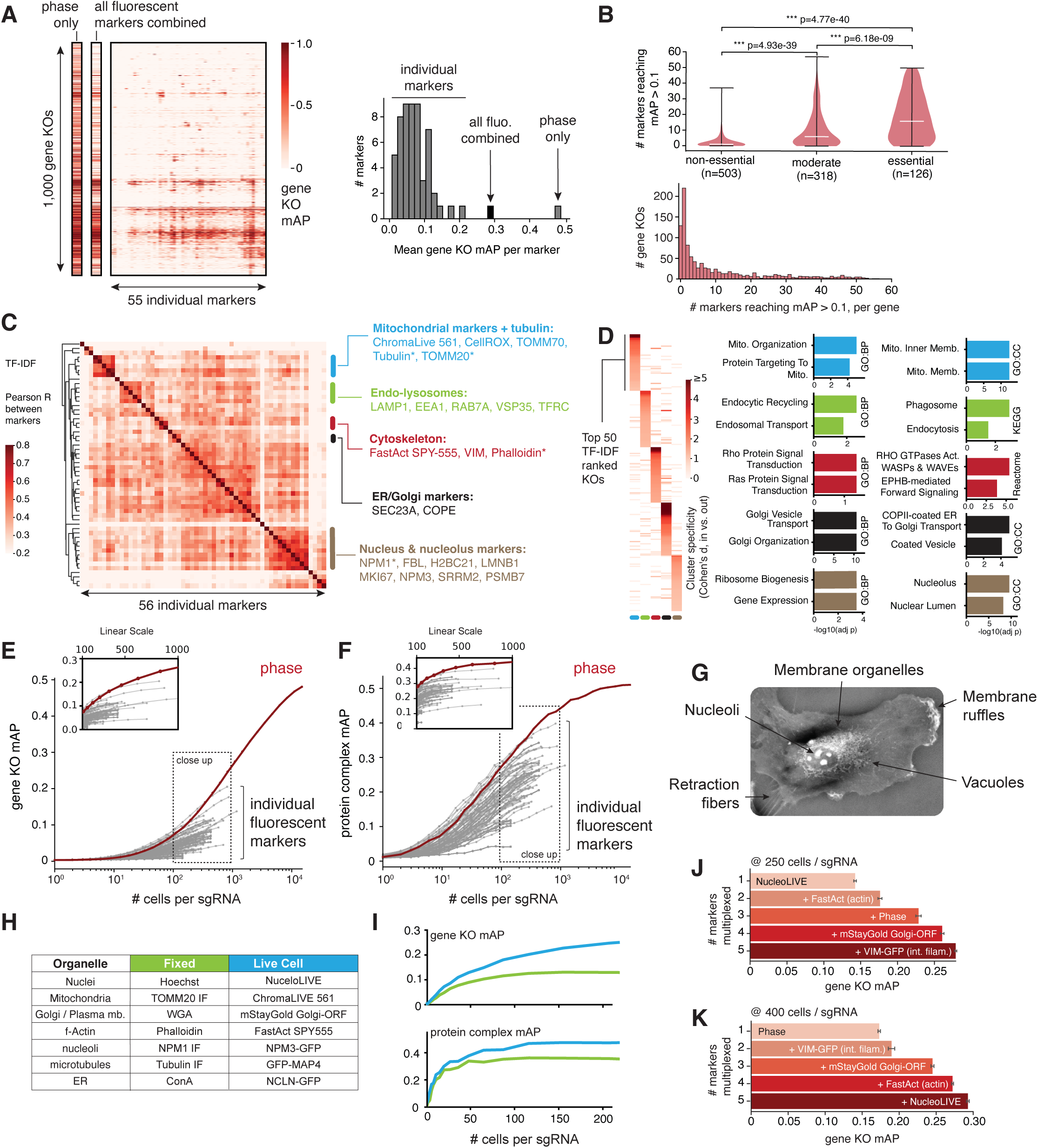
Label-free phase imaging outperforms fluorescent markers at scale. **(A)** Gene-level discrimination performance across imaging markers and modalities, using all available cells per marker. Heatmap shows gene-level mAP for 1,000 gene knockouts (rows) across individual imaging modalities (columns): phase-only, all fluorescent markers combined, and individual-marker representations. See Suppl. Table 7 for individual mAP values for all genes and all markers. Some knockouts have high mAP values across many markers (horizontal lines in the heatmap). These are the knockouts that can be reliably resolved by multiple markers. Right: histogram distribution of mean gene-level mAP across individual markers, with phase-only and all-fluorescence combined values indicated. See Suppl. Table 8 for mean mAP values per channel. **(B)** Number of markers resolving each gene knockout. Top: histogram showing the number of markers with gene-level mAP > 0.1 for each knockout. Bottom: same metric stratified by essentiality class. Violin plots show distributions for non-essential (DepMap scores from -0.5 to 0.5), moderate (DepMap scores -0.5 to - 1.5), and essential genes (DepMap scores -1.5 to -2.5). Pairwise differences were assessed with the two-sided Wilcoxon rank-sum test. Significance is annotated as: * p < 0.05, ** p < 0.01, *** p < 0.001, ns p ≥ 0.05. Essential genes can be resolved by multiple markers independently. **(C)** Correlating the markers according to the genes each marker uniquely resolves reveals similarity in the biology these markers capture. Hierarchically clustered heatmap of pairwise Pearson correlations between the 56 imaging markers (including phase). For each marker, a TF-IDF-weighted gene-level mAP profile was computed across the 1,000 gene knockouts in the screen (see Suppl. Table 9 for per-gene TF-IDF values). The TF-IDF normalization results in a per-knockout adaptive threshold that accounts for the wide dynamic range of mAP values across the 1,000 knockouts. This down-weights knockouts resolved by many markers and amplifies those resolved by few, so the resulting profile emphasizes each marker’s diagnostic hits. The Pearson correlation r between two markers’ TF-IDF profiles measures how similarly their diagnostic signal is distributed across knockouts. The fully annotated heatmap is found in Suppl. Data S1. **(D)** Ontology enrichment analysis from gene knockouts uniquely resolved by the marker clusters highlighted in (C). For each cluster of markers highlighted in (C), the top-50 knockouts with highest TF-IDF-weighted mAP values were identified. How specific these knockouts are to each cluster is measured using Cohen’s d score. Larger d indicates knockouts whose distinctiveness signal is more specifically concentrated in that cluster’s markers rather than spread across the panel. Right: pathway and ontology enrichment for the same per-cluster top-50 driver knockouts, with the 1,000-gene screen library as background. Ontologies: GO-BP, GO Biological Process; GO-CC, GO Cellular Component. **(E) and (F)** Gene-level and protein-complex mAP scores as a function of cell coverage per sgRNA (log10 scale). For each marker and sgRNA combination, random subsets of cell embeddings were selected to produce a down-sampling curve, and Cell-DINO embeddings were generated as in Figure 2. To ensure proper library representation across cell counts, mAP scores are only plotted if more than half of total sgRNAs reach the corresponding x-axis value of cell count. Close-up panels between 100 – 1,000 cells per sgRNA show the transition region (on a linear scale) where 2D phase embeddings begin to achieve the highest mAP scores. **(G)** Example 2D quantitative phase-contrast image of a cell, with visible features annotated. **(H)** Seven different Cell Painting markers were used to image fixed cells, and live-cell markers were chosen to match the covered organelle structures to provide as direct a comparison as possible. **(I)** Cells were down-sampled per sgRNA as in (E) and (F), and embeddings from all 7 live or fixed Cell Painting markers were assembled into a single sgRNA embedding. For both fixed and live cells, different markers were treated as independent channels. **(J)** and **(K)** Gene-level mAP scores upon multiplexing up to 5 imaging markers. In this computational multiplexing, embeddings from individual markers were concatenated and PCA-reduced before mAP score calculation, following the same strategy outlined in Suppl. Fig. 6B. Marker combinations were chosen using a greedy search algorithm. Results are shown at 250 (J) and 400 (K) cells per sgRNA coverage.

The phenotype of a subset of gene knockouts was resolved across many markers at once (Figure 4B). These knockouts were enriched for essential genes, indicating that severe loss-of-function phenotypes are detectable by a wide range of markers. Conversely, the majority of knockouts were resolved by only a small number of markers, suggesting that different markers illuminate complementary aspects of cellular state (Figure 4B, see Suppl. Table 7 for detailed scores per gene). To identify which gene family was preferentially captured by each marker, we re-weighted gene-level mAP scores using the Term Frequency– Inverse Document Frequency (TF-IDF) transformation, a statistical metric used in natural language processing. TF-IDF re-scoring surfaces knockouts that score highly in a given marker but poorly elsewhere, isolating the channel-specific signal from the shared essentiality background (Suppl. Table 9). Hierarchical clustering of channels by the correlation of their TF-IDF–weighted gene profiles partitioned the markers into coherent and biologically interpretable groups (Figure 4C). For example, mitochondrial markers (GFP-TOMM70, TOMM20 immunofluorescence, and ChromaLIVE 561) clustered together across live and fixed conditions, alongside markers reporting indirectly on mitochondrial state: the CellROX oxidative-stress sensor, and a stain for microtubules, which spatially template the mitochondrial network. Gene-ontology enrichment on the top 50 TF-IDF–ranked knockouts per channel recovered sensible biology (Figure 4D). For example, the cluster of mitochondrial markers was enriched for genes involved in mitochondrial organization and protein import, while the actin-marker cluster was enriched for cell-projection regulators and Rho-GTPase signaling — both directly linked to actin dynamics. Collectively, these results show that individual imaging markers share resolving power for the pronounced phenotypes of essential genes but access distinct, complementary aspects of more subtle cellular states.

The analysis above pooled all available cells per channel. In practice, however, the number of cells required to resolve phenotypes is itself a property of each marker. We therefore quantified how gene-level and protein-complex mAP scores scaled with the number of cells imaged per sgRNA (Figures 4E, 4F). Above ∼400 cells per sgRNA, phase embeddings consistently outperformed every fluorescence channel on both tasks. We interpret this performance as a reflection of the information density of phase imaging: a single phase image of a cell simultaneously captures pan-cellular morphology including membrane organelles, plasma-membrane architecture, and sub-nuclear compartments (Figure 4G), whereas a fluorescence image isolates one of these features at a time. Therefore, while phase imaging extracts more dimensions of cellular state per cell, it requires more cells before its higher-dimensional phenotypes can be statistically resolved from one another.

A defining advantage of imaging is that it is acquired directly on live cells. To ask whether the live-cell modality itself, independent of phase, confers a resolution advantage over classical fixed-cell phenotyping, we benchmarked a set of Cell Painting markers in their fixed-cell format against live-cell counterparts of the same markers (Figure 4H). Across both gene-level and protein-complex mAP, and across all cell-coverage regimes, the live-cell panel outperformed its fixed-cell version (Figure 4I). Live-cell imaging can therefore match — and, on these markers, exceed — the phenotypic resolution of established fixed-cell assays.

Finally, we asked which combination of markers maximizes phenotypic resolution at a given cell coverage. We used a greedy selection to build multiplexed panels by adding at each step the marker that, among all others, most improved gene-level mAP. At 250 cells per sgRNA (Figure 4J), the NucleoLIVE DNA dye was the individual channel with the highest average mAP, underscoring the power of chromatin and nuclear morphology as a readout of cellular state^73^. Starting from NucleoLIVE, markers for actin, the Golgi complex^74^, and intermediate filaments were selected in turn, together with phase imaging, raising mAP 1.8-fold overall, with each additional channel contributing progressively less. At 400 cells per sgRNA (Figure 4K), the selected panel was largely unchanged, with the exception that phase imaging was now the highest-resolution single channel. A compact panel of four fluorescent markers and phase imaging therefore captures most of the accessible phenotypic resolution across this range of coverage, with nuclear and cytoskeletal morphology contributing the most information from fluorescence imaging.

### A direct comparison of the phenotypic resolution of imaging and scRNA-seq

A central goal of our atlas was to compare the phenotypic resolution of imaging directly against scRNA-seq, the current benchmark for high-content perturbation phenotyping. We therefore profiled the same 1,000-gene perturbation library by CROP-seq^8^, sequencing a median of 141 cells per sgRNA (564 cells per knockout) at a median depth of 26,103 unique molecular identifiers (UMIs) per cell covering 6,083 expressed transcripts (Suppl. Fig. 7A-D). This matches or exceeds the per-perturbation coverage of published reference datasets (Suppl. Fig. 7A)^11,75^. To convert single-cell transcriptomes into perturbation-level embeddings, we used the deep learning model sVAE+^76^ to decompose gene expression profiles into 50 latent gene programs that represent the transcriptional response to each knockout (Figure 5A).

**Figure 5:**
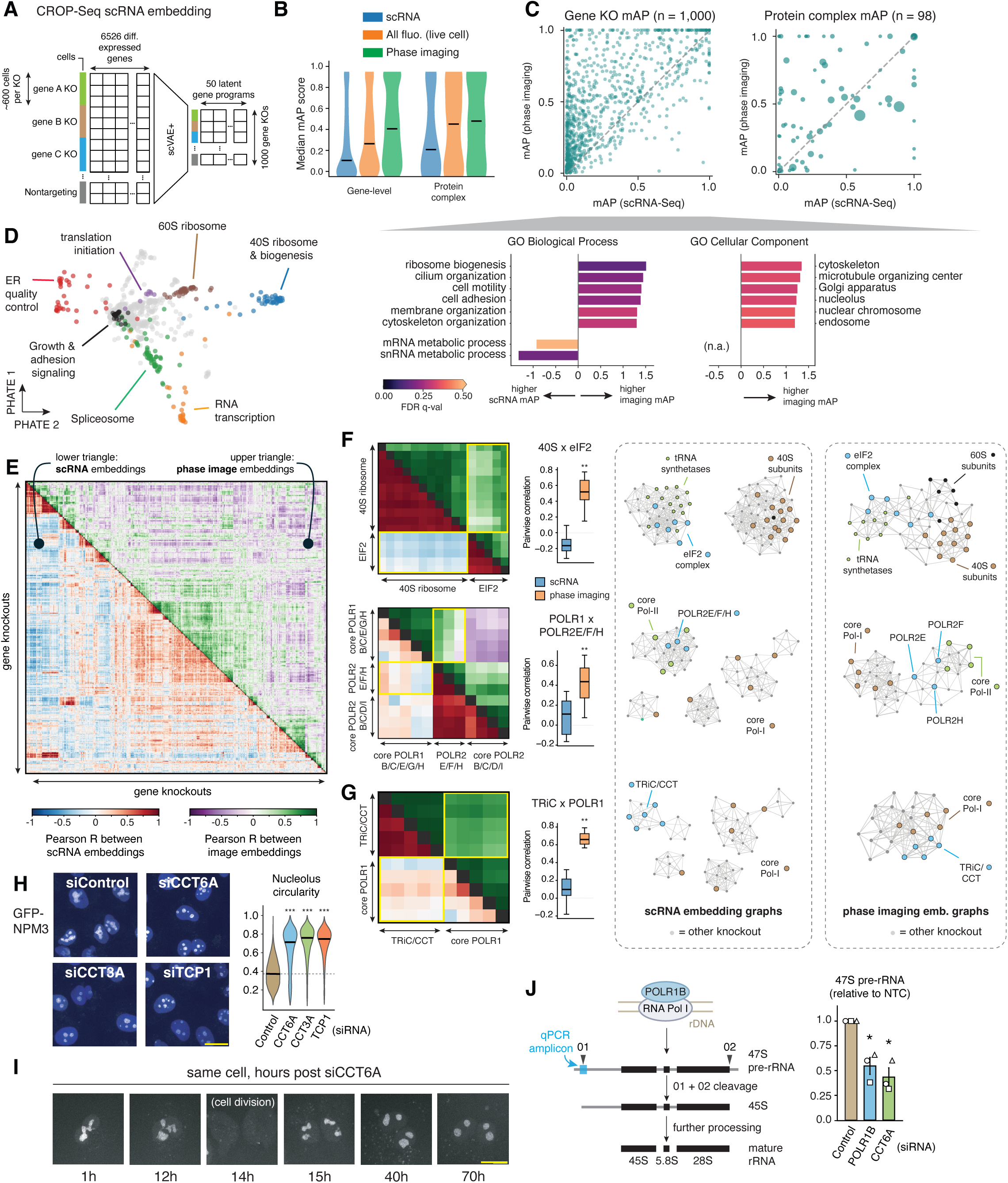
Imaging and scRNA embeddings reveal shared and modality-specific perturbation structure. **(A)** Illustration of our CROP-seq embedding strategy. The sVAE+ model uses a variational autoencoder to learn the sparse transcriptome effect of each knockout perturbation on a set of latent gene programs. We use the learned perturbation effect matrix as our perturbation embedding, compressing the scRNA expression profiles from individual perturbed cells into a sparse 50-dimensional latent vector for each gene knockout. **(B)** Violin plots show distributions of gene-level and protein-complex mAP across image and scRNA embeddings. Horizontal bar: median mAP. **(C)** Scatterplot of individual gene-level (left) and protein-complex (right) mAP scores. The difference between 2D phase imaging and scRNA gene-level mAP scores (distance from diagonal) was used to perform a gene set enrichment test against the GO Slim Biological Process and GO Slim Cellular Component ontology sets (bottom panel). The five most significant terms (excluding near-synonym terms) are shown as bar graphs where the x-axis indicates the normalized enrichment score and the color indicates the negative log10 False discovery rate (q-value) of the enrichment. **(D)** Two-dimensional PHATE visualization of sVAE+ scRNA embeddings for each perturbation. Points are colored according to low-resolution Leiden clusters (resolution = 2), and some clusters with clear biological enrichment are highlighted. See also data in Suppl. Table 11. **(E)** Heatmap of pairwise Pearson correlations of gene knockout embeddings for scRNA (bottom left triangle) and phase imaging (top right triangle). Only knockouts with a correlation coefficient < 0.4 to their nearest 300 neighbors in both modalities are shown (n = 598 gene knockouts), effectively filtering out knockouts that only show promiscuous perturbation effects. See correlation values for imaging and scRNA embeddings in Suppl. Tables 14 and 15, respectively. **(F)** and **(G)** Phenotypic relationships between co-functioning protein complexes in scRNA vs. phase image embeddings. For each set of protein complexes, the pairwise Pearson correlations of knockouts for individual complex subunits are shown as in (E). Top right triangle: image embeddings; bottom left triangle: scRNA embeddings. For each heatmap, the distribution of correlation values in the yellow boxes is shown as boxplots (boxes represent 25th, 50th, and 75th percentiles; whiskers represent 1.5 times the interquartile range; ** p < 0.001 Mann-Whitney U test). On the right panels, the topology of the local k-NN graphs (n_neighbor = 10) around the same protein complexes is shown. For clarity, only nodes connected by >3 edges are shown. **(F)** eIF2 and 40S ribosome protein complexes (top) and RNA Pol I and Pol II protein complexes (bottom). **(G)** TRiC and Pol I protein complexes. Fully annotated panels are found in Suppl. Data S1. **(H)** and **(I)** siRNA knockdown of TRiC subunits results in rounded nucleoli. Images in (H) were taken at 72 h post siRNA. Scale bars = 20 µm. Inset: quantification of nucleolar circularity from image segmentations (n = 3,371 cells per group). *** p < 0.001 (two-sided Mann–Whitney U test). **(J)** qPCR quantification of 47S pre-rRNA levels in knockdown cells. A qPCR amplicon was designed to span the 01 cleavage site (the first step of rRNA processing) in order to only quantify newly synthesized rRNA. Measurements were performed across three independent biological replicate experiments (open symbols). Quantification plots show average ± s.e.m. * p < 0.05 (two-tailed one-sample t-test).

A direct comparison of imaging and scRNA-seq is complicated by the fact that the two modalities cannot be matched simultaneously on cell number and on cost. Under the conditions of our experiments, reagent cost per cell was $0.29 for CROP-seq (including droplet generation, library preparation and sequencing) and $0.002 for OPS (including T7-based *in situ* sequencing), a 145-fold difference. Our imaging and scRNA datasets differ 108-fold in cell number (∼0.6M versus ∼65M cells) and are, as a result, comparable in reagent cost. Using phase imaging as the reference modality, we therefore conducted a comparison of phenotypic resolution at both matched cell coverage (Suppl. Fig. 8A) and matched cost (Suppl. Fig. 8B), which showed opposite trade-offs. To reach the same gene-level phenotypic resolution, phase imaging required 11-fold more cells than scRNA-seq, yet remained 13-fold cheaper. The per-cell advantage of scRNA-seq is intuitive: a transcriptome reports directly on thousands of variables per cell, whereas an image captures a projection of cellular state. In practice, however, cost rather than cell number is the main binding constraint for screening experiments. Within the range of cell number accessible in our experiment, the maximal resolution reached by phase imaging was far higher than that reached by scRNA-seq. Complex-level resolution from the CROP-seq data plateaued (Suppl. Fig. 8A), suggesting that additional cells might not close this gap.

When using all available cells in our datasets (i.e., at matched cost), phase imaging outperformed both scRNA-seq and all fluorescence-based readouts on average (Figure 5B). At the level of individual knockouts, phase imaging achieved a higher score than scRNA-seq for the majority of perturbations (Figure 5C). Gene-ontology enrichment of the knockouts best resolved by each modality recovered a biologically coherent split (Figure 5C, bottom panel): knockouts of genes involved in mRNA expression, splicing, and transport were better captured by scRNA-seq, whereas knockouts of genes involved in cellular structure, migration, and division were better captured by imaging — reflecting the cellular features each modality is intrinsically tuned to read out.

Despite these differences in absolute resolution, the two modalities produced strongly concordant profiles of cell biology. Unsupervised clustering of the scRNA embeddings recovered well-defined clusters of co-functional genes (Figure 5D, Suppl. Table 11), mirroring the organization obtained from imaging (Figure 2). Clusters obtained from scRNA or phase imaging were highly congruent (Suppl. Fig. 9A). To quantify this congruence, we focused on biologically interpretable clusters (i.e., those significantly enriched for one or more gene-ontology terms) and computed the adjusted Rand index^77^ (ARI), a chance-corrected measure of clustering agreement where 1 indicates identical partitions and 0 the level expected at random. The ARI between phase imaging- and scRNA-derived partitions was 0.48, far above the null distribution from permuted labels (Suppl. Fig. 9B). The same concordance was apparent at the gene-pair level: a heatmap of pairwise embedding similarity between knockouts revealed broadly the same gene–gene relationships in imaging and scRNA space (Figure 5E).

A systematic difference between imaging and scRNA emerged on closer inspection. Although both modalities resolved similar fine-grained clusters (which typically corresponded to subunits of a single protein complex), the higher-order organization linking complexes into shared pathways was more reliably recovered by imaging. Two examples illustrate this trend (Figure 5F). First, knockouts of the eIF2 ternary complex and the 40S small ribosomal subunit, which physically assemble into the 43S pre-initiation complex during translation initiation, were strongly correlated in imaging but not in scRNA space (Figure 5F, heatmaps; Suppl. Fig. 8C). Reflecting this correlation, eIF2, 40S, 60S subunits and tRNA synthetases formed inter-connected communities in the image-embedding k-NN graph, while being separated in the scRNA-embedding graph. Second, the shared Pol I/II subunits POLR2E/F/H bridged the Pol I and Pol II clusters in imaging (as described in a previous section), but did not connect them in scRNA (Figure 5F; Suppl. Fig. 8C). To quantify this more systematically, we asked how well each modality grouped genes belonging to the same functional module, using GO Molecular Function and KEGG Pathway annotations. Both ontologies group genes that act together in the cell to carry out a shared, higher-level process. Measuring the fractional connectivity in the k-NN graphs for each ontology term revealed that imaging connected these functional communities more often than scRNA-seq (p = 0.051; Suppl. Fig. 8D). Consistent with the case examples above, this suggests that imaging better captures the integrated cellular consequence of a perturbation across coupled biochemical pathways.

Exploring relationships uniquely surfaced by imaging, we identified a strong phenotypic correlation between knockouts of subunits of the chaperonin TRiC/CCT and knockouts of RNA Polymerase I (Figure 5G, Suppl. Fig. 8C). Inspection of top-predictive cells revealed that TRiC knockouts produced rounded nucleoli, the same phenotype seen for RNA Pol I subunit knockouts and indicative of disrupted ribosomal-RNA transcription^65^. A secondary screen reproduced this result with an orthogonal perturbation: siRNA silencing of individual TRiC subunits induced time-dependent nucleolar rounding (Figures 5H-I). We further established a direct functional link between TRiC activity and rRNA transcription: 47S pre-rRNA levels measured by qPCR were significantly decreased upon CCT6A knockdown, mirroring the effect of knocking down RNA Pol I through its POLR1B subunit (Figure 5J). The precise mechanism by which TRiC acts on rRNA transcription remains to be established, but prior work suggests two connections. First, TRiC is required to fold TCAB1, the chaperone that retains telomerase and scaRNAs at Cajal bodies. TRiC loss depletes TCAB1 and mislocalizes these RNAs to nucleoli^78^, which in turn may affect Pol I activity^79^. Second, a large fraction of ribosome-biogenesis factors (including TCAB1) are WD40-repeat proteins, a substrate class particularly dependent on TRiC for folding^80,81^. The phenotype we observe may therefore reflect collateral disruption of nucleolar ribosome biogenesis by accumulated misfolding of TRiC substrates.

Together, our results establish that high-content live imaging matches and, at least in the conditions of our experiments, can exceed the phenotypic resolution of scRNA-seq given enough cell coverage. Imaging also captures higher-order pathway organization that transcriptional readouts might not resolve. Live-cell imaging — and label-free phase imaging in particular — therefore emerges as a phenotyping modality on par with the highest-information readouts currently in use.

## Discussion

Here, we presented a large multimodal perturbation atlas that serves three complementary purposes. First, it is a compendium of cell-biological observations spanning the function of 1,000 human genes. Our panel of 42 live and 13 fixed-cell markers extends phenotypic coverage well beyond the canonical organelle landmarks of established image-profiling assays. These profiles can be explored interactively at opsin.apps.czbiohub.org. Each marker offers a window onto how the rest of the genome functionally connects to a particular cellular process. Profiling mitochondrial markers, for example, reveals how perturbations across diverse pathways converge on mitochondrial homeostasis. Across our atlas, image embeddings resolved cell biology at the level of individual genes, complexes, and pathways. While the full exploration of these observations and their mechanistic implications is beyond the scope of this first analysis, we introduce complementary deep learning frameworks that surface the most defining single-cell phenotypes of each knockout as a starting point for follow-up. These frameworks serve as a generalizable strategy for the exploration of image-based screening phenotypes.

Second, we provide a rich data resource comprising image profiles from ∼65M single cells (most of them acquired in live cells, including phase images for all cells), and >600k single-cell transcriptomes, matched across 1,000 perturbations. We expect this resource to accelerate discovery and support new approaches in data analysis and AI model development. Mapping phenotypic relationships between modalities (for example, building models that predict transcriptomes from images) is one particularly promising direction. A recent screen combining imaging and Perturb-seq in mouse macrophages showed how OPS image data can be used to predict scRNA signatures in perturbed cells^82^. Our diverse atlas of markers could be used to further refine these predictions, for example by identifying imaging markers whose variations track with specific gene expression modules. Finally, our large corpus could also be used to train new image representation models. Given our results, models trained specifically for phase image representation could be especially relevant. Specialized models might be more efficient at capturing image information than the models we used for the present study, and might further improve the phenotypic resolution that can be achieved from image data. Overall, to maximize access and re-usability, both our raw and processed data are openly available at biohub.ai/ops-explorer.

Third, the atlas allows us to quantitatively benchmark the phenotypic resolving power of different modalities against each other — the central analysis of this study. For matched markers, live-cell imaging outperformed its fixed-cell counterpart. While this may partly reflect differences in marker design and signal quality, it is also consistent with the well-documented effects of chemical fixation and permeabilization on epitope accessibility, protein localization, and subcellular structure^83–85^. Fixed-cell assays remain indispensable for *in situ* molecular detection, multiplexed labeling, and many antibody-based measurements (for example, mapping post-translational modifications). Our results suggest, however, that live-cell measurements preserve phenotypic information that is partially lost or distorted upon fixation, at least in the conditions of our assays.

Beyond comparing live versus fixed conditions, our data establish that label-free quantitative phase imaging exceeded the phenotypic resolution of both fluorescence imaging and scRNA-seq at matched reagent cost. The advantage over individual fluorescence channels is intuitive: each fluorescence channel isolates one molecular or structural landmark at a time, while phase imaging captures cell-wide optical density distribution, organelle organization, membrane architecture, and nuclear morphology simultaneously. Because phase imaging extracts more dimensions of cellular state per cell, more cells might need to be sampled for those dimensions to be statistically resolved. The same coverage trade-off is harder to map out against transcriptomics, since high cell coverage remains prohibitively expensive with scRNA-seq. In practice, the trade-off is therefore also economic: to reach the same phenotypic resolution, phase imaging was 13-fold less expensive at the time of writing.

Fundamentally, imaging and transcriptomics are complementary measurements of cellular states, and combining both captures these states more completely than either alone^45,86,87^. In our data, imaging and scRNA-seq produced highly congruent clustering of co-functional genes and protein complexes, demonstrating that each modality can capture the core modules of cellular organization. Similar results have been very recently described in a multimodal screening study in mouse macrophages^82^. But our results also show that, on average, image embeddings better resolved the higher-order functional connections between these core modules. This suggests that the physical phenotype of a perturbation can integrate consequences across coupled biochemical processes that do not always appear as coherent transcriptional covariation. Mapping this intermediate scale of cell organization (between individual molecular components and whole-cell behavior) is central to understanding the cell as an integrated system^4,5^, and especially relevant for emerging efforts to build causal foundation models and AI virtual cells^1–3^.

Several comparative dimensions remain to be explored for a comprehensive evaluation of the phenotypic resolution of different assays. Our results derive from a single cell line (A549) and a defined perturbation system. Broader generalization will require profiling across diverse cellular contexts, including different cell types and 3D cell models such as organoids. Furthermore, our experimental design focused on single channels of fluorescence imaging at a time. How phase compares to (or could be combined with) highly multiplexed fluorescence remains an open question. Multiplexing in live cells is limited by spectral overlap and phototoxicity, but hyperspectral unmixing^88^, Raman imaging^89^ and compressed-sensing strategies^90,91^ offer solutions to expand live-cell channel capacity. More fundamentally, the best imaging strategy to resolve the greatest range of cellular states remains to be defined. Even with our large coverage of channels, about 18% of the genes tested produced no discernible phenotype in our assays (mAP < 0.1). This might partly reflect incomplete perturbation penetrance, compensation, or functional redundancy, but many perturbations may also require bespoke imaging markers to surface their phenotypes (e.g., dedicated biosensors). In both cases, multiplexed live-cell measurements (including with phase imaging) would offer a natural route forward.

Taken together, our results position live-cell imaging as a powerful and scalable modality for functional genomics. Methods like Cell Painting^19,20^, OPS^27–31^, pooled scRNA-seq screens^8,9,11,92^ and emerging multimodal technologies^87,93^ have established that perturbation atlases are powerful tools to chart the cellular phenotypic landscape. However, the trajectories by which cells navigate this landscape have remained largely inaccessible at scale because most studies are conducted in fixed or disrupted cells. Our results show that live-cell imaging, and in particular label-free morphology, can reach comparable phenotypic resolution while preserving access to time, movement, and native cellular context. The next frontier will involve perturbation campaigns in which the cells are repeatedly observed as they transition through state space. By enabling scalable live-cell OPS, and demonstrating the resolving power of intrinsic morphology, this work lays a foundation for such time-resolved atlases of cellular function.

## Acknowledgements

We sincerely thank P. Blainey, I. Cheeseman, M. Di Bernardo, and their teams for sharing protocols and advice for setting up OPS. We thank T. Kudo for advice with *in situ* sequencing; J. Bragantini for advice with cell tracking; C. Januel and R. Baltazar-Nunez for help with tissue culture; R. Arjyal and A. Seng for high-throughput sequencing; I. Gray and J. Truong for mass spectrometry; S. Varra for support with computational microscopy; A. Kalinin for introducing us to the mAP framework; M. Logan, G. Yun, J. Gadiane, J. Panganiban and J. Mann for operational support; and S. Schmid and J. Swedlow for critical feedback. M.D.L. thanks C.L. Tan for continuous discussions. Some figure elements were created using BioRender.com. We thank the Biohub and its donors, Priscilla Chan and Mark Zuckerberg, for funding this work.

**Materials and Methods** are available as a Supplementary File.

## Data and Code availability

All raw and processed data are publicly available at biohub.ai/ops-explorer, where they can be interactively explored and downloaded. The OPSin viewer is publicly available at opsin.apps.czbiohub.org.

The analyzed data used in figures for this manuscript are archived on Zenodo (zenodo.org/records/21970895). All code associated with this paper is available at github.com/czbiohub-sf/cyclops-monorepo. Within this repository, specific sub-repositories were created for image-preprocessing and ISS base-calling (github.com/czbiohub-sf/cyclops-process), feature extraction, embedding, and interpretability (github.com/czbiohub-sf/cyclops-model), organelle segmentation (github.com/czbiohub-sf/organelle-profiler), and shared utilities (github.com/czbiohub-sf/cyclops-utils).

## Declaration of generative AI and AI-assisted technologies in the writing process

During the preparation of this work, the authors used Claude Opus 5.0 to improve the readability and language of the manuscript. After using this tool, the authors reviewed and edited the content as needed and take full responsibility for the content of the published article.

## List of Supplementary Tables

**Table 1 - CRISPR KO library**. Related to Figure 1A and Supplementary Figure 1A.

**Table 2 - Gene effect (DepMap fitness**). Related to Figure 1D and Figure 4B.

**Table 3 - Live cell dyes**. Related to Figure 1E and Supplementary Figure 3.

**Table 4 - Endogenous tagging constructs**. Related to Figure 1E and Supplementary Figure 3.

**Table 5 - List of EBI protein complexes**. Related to Figure 2C.

**Table 6 – Automation hardware bill of materials**. Related to Figure 1C and Supplementary Figure 1B.

**Table 7 – Gene-level mAP per marker**. Related to Figure 4A.

**Table 8 – Average mAP per marker at maximal cell coverage**. Related to Figure 4A.

**Table 9 - TF-IDF values for each gene KO across all imaging markers.** Related to Figure 4C.

**Table 10 – Image embeddings (using all markers + phase).** PHATE, UMAP and Leiden definition. Related to Figure 2E.

**Table 11 – scRNA embeddings (from CROP-Seq): PHATE, UMAP and Leiden definition.** Related to Figure 5D.

**Table 12 - Volcano plot metadata, feature sources and explanations.** Related to Figure 1F-I and Supplementary Figure 5.

**Table 13 - Volcano plot raw data.** Related to Figure 1F-I and Supplementary Figure 5.

**Table 14 - Gene/gene Pearson correlations for phase imaging Cell-DINO embeddings.** Related to Figure 5E.

**Table 15 - Gene/gene Pearson correlations for scRNA sVAE+ embeddings.** Related to Figure 5E.

**Table 16 - Set classifier gene-KO accuracy per marker, at a bag size of 100 cells.** Related to Figure 3B.

**Table 17 - Set classifier protein-complex accuracy per marker, at a bag size of 100 cells.** Related to Figure 3B.

## List of Supplementary Files

**Data S1: fully annotated panels from Figures 2, 3 and 5**.

## Materials and Methods

From Liu et al. *A multimodal perturbation atlas defines the phenotypic resolution of cellular morphology*.

## 1. Cell Engineering and Quality Control

### 1.1 Cell Culture

All cells unless otherwise indicated were cultured in standard conditions with DMEM GlutaMAX (Gibco, 10566024) supplemented with 10% FBS (Omega Scientific, cat. no. FB-11) at 37°C, 5% CO_2_ and detached using 0.25% trypsin-EDTA (Gibco, cat. no. 25200056).

### 1.2 Generation of clonal Cas9 cell line

HP138-neo PiggyBac construct (Addgene #134247), encoding Cas9 and rtTA under a Tet-inducible promoter, was co-transfected with the Super PiggyBac transposase (System Biosciences, PBDNA) at a 1:1 mass ratio (1 µg HP138: 1 µg transposase) using FuGENE HD-1000 at a 3:1 reagent:DNA ratio. Stable integrants were selected beginning 3 days post-transfection with 500 µg/mL G418, with media refreshed every 3–4 days. Selection was completed after 2 weeks when no viable cells remained in untransfected control wells.

Pooled stable cells were single-cell sorted into 96-well plates in DMEM GlutaMAX supplemented with 20% FBS. Clones were expanded and validated using four criteria: inducible Cas9 expression, Cas9 activity, normal growth, and intact p53/p21 induction. Inducible Cas9 expression was assessed after 24 h of treatment with 1 µg/mL doxycycline by staining with anti-Cas9 antibody (Cell Signaling Technology, #14697; 1:500; overnight at 4°C), followed by goat anti-rabbit IgG (H+L) Alexa Fluor 647 secondary antibody. Cas9 activity was evaluated by transduction with pXPR_011, a self-targeting GFP reporter construct that expresses both GFP and an sgRNA targeting GFP. Clones with robust Cas9 induction, high GFP knockout efficiency, normal growth relative to parental A549 cells, and an intact p53 response were selected for downstream engineering. To confirm intact p53 signaling, clones were treated with 40 µM etoposide for 24 h and stained overnight at 4°C for p53 (Cell Signaling Technology, #2524S) and p21 (Cell Signaling Technology, #2947S). Samples were then stained for 1 h with goat anti-mouse IgG (H+L) Alexa Fluor 488 (Invitrogen, #A-11001) and goat anti-rabbit IgG (H+L) Alexa Fluor 647 (Invitrogen, #A-21245), each at 1:1000 dilution, before imaging with a Leica 20×/0.55 air objective.

### 1.3 Guide RNA and HDR donor design for CRISPR knock-in

All cell lines expressing endogenously tagged fluorescent proteins were generated by CRISPR knock-in using the monoclonal dox-inducible-Cas9 A549 cell line described above. Guide RNAs and homology-directed repair (HDR) donors for fluorescently labeling each protein of interest were designed using the protoSpaceJAM platform^1^ or sourced from previously validated tagging libraries^2^. For protoSpaceJAM, the following settings were used: SpCas9 was used as the nuclease, and three candidate gRNAs were generated per locus. Guides targeting 5′ untranslated regions or regions proximal to splice junctions were penalized in the guide selection process. HDR donors were designed with 500 bp homology arms, with trimming constrained to a minimum of 300 bp. Recoding was applied at full intensity, with prioritization of PAM-disrupting mutations, and a recut potential threshold of 0.03. The optimal guide RNA and HDR donor pair was chosen based on the highest ranking produced by the protoSpaceJAM algorithm.

Fluorescent reporter payloads were incorporated between the homology arms: 3xHA-mEGFP-XTEN80 was used for N-terminus tagging, and XTEN80-mEGFP-3xHA was used for C-terminus tagging as in Hein et al.^2^, which were specified as “terminus- and/or position-specific payloads” within the ProtoSpaceJAM design platform. For tagging SEC61B and H2BC21, mScarlet-I was used as the fluorescent tag. Single-strand RNAs encoding guide RNAs were purchased from Integrated DNA Technologies (IDT DNA). HDR donor sequences with PCR primer binding sites were synthesized and cloned into the pTwist Amp high plasmid backbone by Twist Bioscience. The double-strand DNA HDR donors were generated via PCR amplification from the synthesized plasmids using KAPA HiFi HotStart ReadyMix (Roche, 07958919001), supplemented with 2 µM biotinylated PCR primers and 1 M Betaine. Final PCR products were purified using 0.7x SPRI beads and eluted in 20 µL of nuclease-free water.

### 1.4 RNP complex electroporation and cell enrichment by FACS

To prepare Ribonucleoprotein (RNP) complexes and HDR donors, 11.52 µg of *Streptococcus pyogenes* Cas9 (SpCas9), 4 µg of guide RNA, and 0.8 µg of HDR donor were combined on ice. For the RNP complex formation targeting RAB7A, MAP4, VAPA, and TFRC, 90 pmol of target-specific crRNA and 90 pmol of tracrRNA were combined and incubated at 70°C for 5 min, followed by 37°C incubation with SpCas9 for 10 min. For electroporation, 220,000 cells were spun down at 500*g* for 5 min at room temperature. The cells were washed twice with 1 mL of DPBS (Gibco, 14190250) and then resuspended in 16.4 µL of SF Cell Line Nucleofector® Solution supplemented with 3.6 µL of Supplement 1 solution (Lonza, V4SC-2096). The resuspended cells were combined with the RNP complex and HDR donor, transferred to a 96-well Nucleocuvette Plate (Lonza, V4SC-2096), and electroporated using the FF-130 program on an Amaxa Nucleofector 96-well Shuttle Device (Lonza, AAM-1001S). Following electroporation, cells were transferred to a 12-well culture plate containing 1 mL of culture media supplemented with 1 µM M3814 for recovery. One week post-electroporation, cells were sorted to enrich the top 0.1% of fluorescent-positive cells (Sony Cell Sorter SH800 or Invitrogen BigFoot Cell Sorter). Following culture expansion, a second round of FACS was performed to remove fluorescent-negative populations.

### 1.5 UPRE reporter cell line generation and validation

The 5xUPRE-mNG reporter was built into a backbone for lentiviral expression that has been previously characterized (plasmid pBA407 - Addgene #85970). This parental vector was digested with SalI and BstBI to swap mCherry for mNeonGreen2. The UPRE promoter region contains 5 UPR elements (UPREs, 5′-TGACGTGG-3′) upstream of the c-fos minimal promoter as previously described^3^. Cells were seeded in a T75, and the following day, cells were transduced with 2 mL of lentiviruses expressing the 5xUPRE-mNG reporter and incubated at 37°C for 72 hours. The cells were expanded in two T175 and incubated for 24 hours. To enrich for cells expressing the reporter, cells from one T175 were treated with 100 nM Thapsigargin (Tgn) for 24h before FACS sorting. To assess reporter efficiency, single cell clones were seeded in a 96-well plate at 1e4 cells/well for 24 hours, then treated with 100 nM Tgn or a DMSO control for 24 hours. The reporter activation in individual clones was analyzed by flow cytometry on a Cytoflex (Beckman), and the clonal cell line with the best signal-to-noise ratio and homogeneity of reporter activation was selected.

### 1.6 cis-Golgi reporter cell line generation

To establish the cis-Golgi reporter cell line, a landing pad was first integrated into the AAVS1 safe harbor locus of the monoclonal dox-inducible-Cas9 A549 line via CRISPR knock-in. The landing pad construct comprised a Pa01 recombinase-P2A-mTagBFP2 cassette^4^ driven by a CAG promoter with chicken HS4 and UCOE insulator elements^5^, with a Pa01 attB site positioned between the promoter and the recombinase.

BFP-positive cells were enriched by FACS, single-cell cloned, and validated using a co-culture assay. The cis-Golgi reporter payload—consisting of a Pa01 attP site, an mStayGold-tagged CENPR altORF peptide^6^, and an hGH polyA sequence flanked by WPRE and cHS4 insulators—was subsequently electroporated into the validated landing pad clone. Reporter-integrated cells were isolated by two rounds of FACS enrichment for GFP signal.

### 1.7 Competitive Fitness Co-culture Assay

To ensure tagged cell lines maintained stable fluorescent expression and a growth rate within the range of the parental line, co-culture competition assays were conducted over 10 days for each endogenously tagged cell line. Tagged cells were seeded individually at a density of 20,000 cells/well in a 24-well culture plate, along with the untagged parental control. Each cell line was also seeded at an equal ratio with the parental line (co-culture), by plating 10,000 cells each for a total of 20,000 cells per well. Fluorescent intensity distributions of these cultures were measured using the CytoFLEX flow cytometer at 3-4 day intervals over the 10-day assay period. The relative growth rate of the tagged line compared to the parental line at day 4 was determined by fitting the time-course data of the fluorescent-positive cell fraction in the co-culture. The calculation is described below:

Assuming exponential growth kinetics, the relative growth rate of tagged cells (*y*(*t*)) and parental cells (*x*(*t*)) at time *t* with the same initial conditions (*x_0_* = *y_0_*) was defined by the following equation:

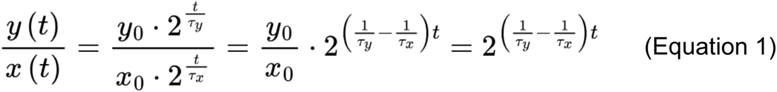

Here, the difference of the reciprocals of their doubling time is an unknown parameter. To estimate this unknown parameter, the time-course data for the fluorescent-positive fraction in the co-culture was fitted to the co-culture growth competition model as below:

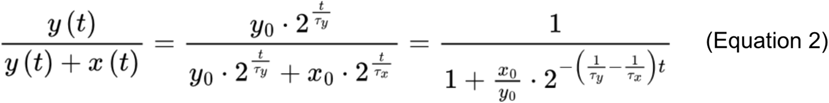

The estimated parameter was then substituted into Equation 1, setting time *t* to 4, to calculate the relative growth rate at day 4.

Tagged cell lines exhibiting a relative growth rate less than 72% of the parental line, signal loss over the 10-day period, or inconsistent localization patterns were excluded from subsequent screens.

### 1.8 Live-cell dye growth QC

To determine the optimal concentration of live-cell dyes that does not impair cell growth, dox-inducible-Cas9 A549 cells were plated at 20,000 cells/well in a 24-well plate. The following day, media was replaced with dye-containing culture media at concentrations ranging from 0.1X to 2X. After 24 hours, the total cell count was quantified by flow cytometry. The concentration resulting in a cell count greater than 90% of the no-dye control was selected for screening.

Dye concentrations and treatment conditions can be found in Supplementary Table 3.

## 2. CRISPR library design

The 1,000 genes were selected to maximize the cell state space covered by perturbations, while ensuring balanced representations of pathways annotated in the KEGG, GO, and REACT ontology databases. We used previous genome-scale screens^7–9^ as a reference to identify perturbations that diversify phenotypic cell state.

Briefly, genes were selected from each cluster in the Funk et al.^7^ OPS dataset based on their proximity to the cluster centroid and proportionally to the size of the cluster, with a 1:1 ratio of genes from ‘non-outlier’ and ‘outlier’ clusters which separate from NTC-targeted cells. Genes were also selected from the Replogle et al.^8^ Perturb-seq dataset based on their distance in embedding space from each other and NTCs, with genes that showed higher correlation between effects in K562 and RPE1 cell lines prioritized. UPR-specific genes^3^, mitochondrial-specific hits^9^, and other genes needed to provide complete REACT pathway coverage were manually added to this gene panel. Guides were selected using algorithms from Feldman et al.^10^, with prefix_length of 7 and edit_distance of 1 (with final sequencing length of 10 bp to assist error correction). In addition to the existing Brunello and TKOv3 libraries, GeCKO and CRISPick libraries were also included^11–14^. Due to Brunello being prioritized during selection, 82.6% of sgRNAs selected were from Brunello. Library information can be found in Supplementary Table 1.

## 3. Library transduction

### 3.1 Library construction

To construct a PerturbView-compatible CROP-seq vector, inverse PCR was performed using the QuikChange II XL Site-Directed Mutagenesis Kit (Agilent, #20052) and CROPseq-puro-v2 plasmid (Addgene, #127458). The modification was introduced in two steps: first, deletion using primers:

5’-ataatttcttggggtagtttgcagtttgcactatcatatgcttaccg-3’ and

5’-cggtaagcatatgatagtcccaaaactgcaaactacccaagaaattat-3’, then insertion using primers:

5’-ataatttcttggggtagtttgcagtttttaatacgactcactatagggactatcatatgcttaccg-3’ and

5’-cggtaagcatatgatagtccctatagtagtgtattaaaactgcaaactacccaagaaattat-3’.

This generated a backbone containing a 13-bp replacement in the U6 promoter. The 1,000 gene library was then synthesized and cloned by Twist Bioscience. NGS validation showed a 95th/5th percentile skew ratio of 2.19, and 96.7% of sequences correct.

### 3.2 Lentivirus generation

HEK293Ts were plated in a 6-well plate at a density of 800,000 cells/well. The following day, after confirming that the cells were ∼70% confluent, cells were transfected with gag/pol, REV, TAT, and VSVG packaging plasmids (1:1:1:2 ratio by mass) and library plasmid pool using Mirus-LT1 (Mirus Bio, MIR 2300). The next day, the media was replaced by DMEM supplemented with ViralBoost (Alstem, VB100). The following day, the top 1 mL of the supernatant in each well was harvested and stored in -80°C.

### 3.3 Titer determination

Titer was determined by following Feldman et al.^10^. Cells were seeded at a density of 300,000 cells/well in 4 wells of a 12-well plate. The next day, 0 µL, 1 µL, 5µL, and 25 µL of collected lentivirus was added to separate wells. 24 hours later, cells were trypsinized and 1/4 and 1/40 of cells from each condition were moved to separate wells of a new 12-well plate with growth media. Puromycin (InvivoGen, cat. no. ant-pr-1) was added to the higher density sample from each condition for a final concentration of 1 µg/mL. Two days later, the untransduced cells were checked for complete killing. Viable cells were then trypsinized and counted with a flow cytometer after washing off the dead cells.

For each condition, the following was estimated:

● MOI = 0.1 x (cell count from 1/4 seeding density well with antibiotic)/(cell count from 1/40 seeding density well without antibiotic)
● Viral titer = (total number of cells x MOI)/virus volume

The total number of cells was the count assumed at the time of transduction. When seeding 300,000 cells, the number inputted into the equation would be 600,000 to account for the approximate doubling of cells overnight.

### 3.4 Transduction and Cas9 induction

Cells were transduced at an MOI of 0.1 while maintaining at least 500× library coverage. Lentivirus was added directly to the culture medium at 37°C for 24 h, after which the medium was replaced with regular growth medium containing 1–2 µg/mL puromycin while maintaining at least 1,000× library coverage. After 9–11 days of selection and routine passaging in puromycin, cells were seeded in 6-well glass-bottom plates (Cellvis, P061.5HN) at 80,000 cells per well with 1 µg/mL doxycycline to induce Cas9 expression. Twelve hours before imaging, the medium was exchanged for phenol red-free DMEM (Gibco, #31053028) supplemented with 10% FBS and, where applicable, live-cell dyes. Cells were phenotyped 86–94 h after doxycycline addition.

## 4. Live-cell phenotyping

Live-cell imaging was performed on a Leica DMi8 microscope with an Andor Dragonfly spinning disk confocal module (model number CR-DFLY-502 with 40 µm pinhole size). Cells were maintained at 37°C and 5% CO2 using an OkoLab environmental control chamber. Data for image-based phenotyping were collected at 20× magnification (Leica 20 0.55 NA, model number 506519) in up to three channels acquired simultaneously. All experiments acquired a brightfield channel using 448/20 nm 0.4 NA Kohler illumination, and optionally fluorescence channels using 488 nm, 561 nm, or 637 nm excitation and detection through 525/50 nm, 610/75 nm, or 700/75 nm emission filters. Transmitted light was directed towards a custom imaging port at the back of the microscope using a 475 nm long-pass dichroic beamsplitter positioned at the microscope filter turret and imaged on an Andor Zyla camera. Fluorescence channels were imaged on Teledyne Photometrics Prime BSI (shorter wavelengths) and Prime BSI Express (longer wavelengths) cameras using a Cairn TwinCam image splitter with exchangeable 560 nm and 660 nm long-pass dichroic beamsplitters.

In each well of the plate we acquired 2,345 fields of view distributed in a circle at the center of the well with 8% overlap between adjacent fields of view. At each field of view we acquired 7 z-slices spaced 2 µm apart. In order to track these motile cells over the duration of the experiment we further interleaved 5× imaging (Leica 5x 0.15 NA, model number 556058) between phenotyping runs of each well. We acquired 148 fields of view with 8% overlap in each well, collecting transmission images with 0.07 NA illumination from 9 z-slices spaced 25 µm apart.

Prior to each live imaging experiment, we imaged 1 µm FluoSpheres carboxylate, yellow-green (505/515) (Invitrogen F8823) suspended in 1% agarose solution (Bio-rad #1613101) diluted in distilled water. The beads were imaged in 28 z-slices spaced 1 µm apart. The beads were simultaneously captured in the brightfield channel and the fluorescence channel. For fluorescence, the beads were excited with the 488 laser and were detected through the 525/50 nm and 610/75 nm emission filters.

The microscope was controlled through µManager^15,16^ and data were collected with a custom Python script using Pycro-Manager^17^. Live-cell imaging experiments lasted ∼7 hours.

## 5. Fixed cell phenotyping

### 5.1 Iterative organellular staining

The live-cell label-free acquisition protocol was performed as usual (see Live imaging). Cells were immediately fixed with 4% paraformaldehyde for 20 minutes, permeabilized with 0.1% Triton X-100 (Sigma, T9284) in PBS for 10 minutes and blocked with 1% Bovine Serum Albumin (BSA) (VWR, 0332) in PBS for 30 minutes. Samples were washed three times with PBS between each step. A master mix with Hoechst 33342 (Invitrogen, H3570) (diluted 1:5000), Wheat Germ Agglutinin Alexa 555 (Invitrogen, W32464) (diluted 1:666), TOMM20 primary antibody (abcam, ab186735) (diluted 1:500) and phalloidin Alexa 650 (Invitrogen, A30107) (diluted 1:1000) was made and used to stain samples at room temperature for 45 minutes. Samples were washed three times with PBS and incubated with a goat anti-rabbit IgG (H+L) Alexa Fluor 488 antibody (Invitrogen, A11008) (diluted 1:500) for 15 minutes. Samples were washed three times and imaged in PBS. For fluorophore bleaching, a solution of 1 mg/mL lithium borohydride (Strem, 9303971G) was freshly prepared and rested for 10 minutes before incubating samples with the solution for 1 hour at room temperature. Samples were then washed three times with PBS and re-blocked with 1% BSA. For the second round of staining, a master mix with alpha-tubulin primary antibody conjugated with Alexa 555 (Cell Signaling, 5059) (diluted 1:200), NPM1 primary antibody conjugated with Alexa 488 (Invitrogen, MA3-25200-A488) (diluted 1:3000) and concanavalin A - Alexa 647 (Invitrogen, C21421) (diluted 1:2000) was made and used to stain samples at room temperature for 45 minutes. Samples were washed three times with PBS and imaged. Once imaging was complete, samples were once again bleached with lithium borohydride and then processed for in situ sequencing following the protocol in Kudo et al.^18^. The same imaging conditions used for live-cell phenotyping were applied: a 20×/0.55 Leica air objective on a Leica DMi8 microscope equipped with an Andor Dragonfly spinning-disk confocal module, with 7 z-slices acquired at 2 µm increments.

### 5.2 Iterative immunofluorescence of signaling proteins

Multiplexed immunofluorescence was performed by iterative indirect immunofluorescence imaging (4i) as described^19^, with the following modifications. Cells were seeded in 6-well plates and fixed in ∼4% paraformaldehyde for 20 minutes at room temperature, then permeabilized in 0.5% Triton X-100 in PBS for 10 min. In each cycle, one mouse and one rabbit primary antibody were co-applied at 1:500 in conventional blocking buffer (1% w/v BSA in PBS) for 1 h at room temperature, followed by a 15-min incubation with goat anti-mouse IgG (H+L) Alexa Fluor 488 (Invitrogen A-11001) and goat anti-rabbit IgG (H+L) Alexa Fluor 647 (Invitrogen A-21245), each at 1:500. Nuclei were counterstained with Hoechst 33342 (1:10,000 in PBS) for 5 min prior to imaging in 4i imaging buffer (700 mM N-acetyl-L-cysteine, 100 mM HEPES, pH 7.4).

The following primary antibodies were used, each at 1:500: anti-p53 (clone 1C12, mouse monoclonal, Cell Signaling Technology Cat# 2524); anti-β-catenin (clone L54E2, mouse monoclonal, Cell Signaling Technology Cat# 2677); anti-c-Myc (clone Y69, rabbit recombinant monoclonal, Abcam Cat# ab32072); anti-phospho-Rb (Ser807/811) (clone D20B12, rabbit monoclonal, Cell Signaling Technology Cat# 8516); anti-p21 Waf1/Cip1 (clone 12D1, rabbit monoclonal, Cell Signaling Technology Cat# 2947); and anti-phospho-S6 ribosomal protein (Ser235/236) (rabbit polyclonal, Cell Signaling Technology Cat# 2211).

Images were acquired on a Leica DMi8 with a Hamamatsu Orca Fusion BT with a HC PL APO 20×/0.80 dry objective, controlled by µManager. Samples were excited with a Leica mercury arc lamp and emission was collected through one of the Semrock filter sets (dichroic centerlines 552, 652, GFP and DAPI). A single z-plane was collected per field of view in three channels (Hoechst, Alexa Fluor 488, Alexa Fluor 647).

## 6. T7-based barcode amplification

Our protocol for barcode amplification and detection closely follows Feldman et al.^10^ and Kudo et al.^18^, with some modifications.

### 6.1 Fixation

Cells were fixed with a final working concentration of 4% paraformaldehyde (Electron Microscopy Sciences, SKU: 15710) as soon as image acquisition was complete, followed by 3 washes with PBS (Gibco, cat. no. 14190094). Endogenously tagged cell lines were chemically bleached with a solution of 3% hydrogen peroxide (Millipore Sigma, cat. no. H1009) and 20 mM HCl (Sigma, cat. no. H9892) in DI water for 10 minutes^20^, followed by 3 washes with PBS. 70% ethanol (VWR, E505) was used to permeabilize the cells at room temperature for 30 minutes, followed by 6 dilutive washes with 1X DPBS + 0.05% Tween (PBS-T) (Teknova, cat. no. T0710), and 2 more standard PBS-T washes. Heat decrosslinking was performed by adding 0.3 M sodium chloride (Invitrogen, ref. No. AM9759) in 0.1 M sodium bicarbonate (Millipore Sigma, cat. no. 36486) and incubated at 65°C for 4 hours.

### 6.2 In vitro transcription

After decrosslinking, samples were washed 3 times with PBS-T. IVT solution was assembled from the HiScribe T7 Quick High Yield RNA Synthesis Kit (New England Biolabs, cat. no. E2050L), using 0.53× T7 polymerase, 10 mM NTP Buffer Mix (New England Biolabs, cat. no. E2050L), 5 mM DTT (New England Biolabs, cat. no. E2050L), and 0.4 U/µL RiboLock (Thermo Fisher Scientific, cat. no. EO0384) diluted in DI water. Plates were sealed to prevent evaporation and incubated at 37°C overnight.

### 6.2 Reverse transcription

After incubation, the IVT solution was removed, and one PBS-T wash was performed. 1 µM 5’ Locked Nucleic Acid (5’ LNA) reverse transcription primer with 5’ primary amine (custom order from IDT: /5AmMC12/A+CT+CG+GT+GC+CA+CT+TTTTCAA)^21^ in PBS-T with RiboLock was added to the wells, and incubated at room temperature for 30 minutes. The cells were fixed after one PBS-T wash in a solution of 3.2% paraformaldehyde and 0.1% glutaraldehyde (Electron Microscopy Sciences, cat. no. 16120) in PBS-T at room temperature for 30 minutes. Fixation was quenched with Tris-HCl (Teknova, cat. no. T1080) at a 1:1 ratio of fixative, and then wells were washed 3 times with PBS-T. A freshly made solution of 4.8 U/µL H-minus Reverse Transcriptase enzyme and 1× buffer (Thermo Fisher Scientific, cat. no. EP0452), 250 µM dNTPs (New England Biolabs, cat. no. N0447L), 1 µM 5’ LNA reverse transcription primer, 0.2 U/µL recombinant albumin (New England Biolabs, cat. no. B9200S), and 0.4 U/µL RiboLock diluted in sterile DI water was added to each well to begin reverse transcription of the IVT product. Plates were incubated at 37°C for 3 hours, then 42°C for 30 minutes.

### 6.3 Rolling circle amplification

Reverse Transcription solution was removed and plates were washed 6 times with PBS-T. Samples were fixed in 3.2% paraformaldehyde and 0.1% glutaraldehyde in PBS-T at room temperature for 30 minutes, and then washed 3 times with PBS-T. Gap-fill solution containing 0.1 µM of the padlock probe: /5Phos/GTTTCAGAGCTATGCTCTCCTGTTCGCCAAATTCTACCCACCACCCACTCTCCAaaggacgaaaCACCg,

50 nM dNTPs, 0.4 U/µL RNAse H (Qiagen, cat. no. Y9220L), 0.02 U/µL TaqIT enzyme (Qiagen, cat. no. P7620L), 0.5 U/µL Ampligase and 1X buffer (Lucigen, cat. no. A3210K) with 0.2 mg/µL recombinant albumin diluted in DI water was added to each well and incubated at 37°C for 5 minutes, then 90 minutes at 45°C. After 3 PBS-T washes, the RCA solution was made with 5% glycerol (Sigma-Aldrich, cat. no. G5516), 250 µM dNTPs, 1 U/µL Phi-29 enzyme and 1X buffer (Thermo Fisher Scientific, cat. no. EP0094) in DI water and added to each well and incubated overnight at 30°C. After 17-20 hours, plates were washed 3× the next morning with PBS-T. To prepare for sequencing-by-synthesis, SBS primer:

5’-GCCAAATTCTACCCACCACCCACTCTCCAaaggacgaaaCACCg

1 µM in 2x SSC buffer (Invitrogen, cat. no. AM9770) was added to each well and incubated at 37°C for 30 minutes, followed by 3 PBS-T washes. The samples were then stained with 1:10,000 Hoechst 33342 (Invitrogen, H3570) in 2x SSC buffer. Plates were stored at 4°C in 2mL PBS-T per well.

## 7. In situ sequencing

### 7.1 Reagents

Incorporation and cleavage mixes (Illumina MiSeq Nano v2, MS-102-2003; reagent 1 and 4) were kept at 4°C, light protected. PR2 was brought to room temperature and diluted up to 50% with PBS-T before beginning Automated ISS.

### 7.2 Hardware instrument overview

The automated *in situ* sequencing system was built by combining a custom fluidic automation system with a commercial epifluorescence microscope (Leica DMi8) to enable sequencing in three wells of a six-well plate. The microscope is equipped with an Andor Zyla 4.2 sCMOS camera, an Okolab Cage Incubation system (H203-T-ENCLOSURE), a mercury arc excitation lamp (Excelitas X-cite Exact), and five Semrock filters optimized for the MiSeq dyes (Semrock, dichroic centerlines: 552 nm, 596 nm, 652 nm, 695 nm, and DAPI - see the Bill of Materials (BOM) included in the Supplementary Table 6). The wells were imaged with a 5x objective (Leica HC PL FL 5×/0.15).

The incubation chamber was set to the maximum temperature setting (55°C), and custom insulating sleeves were added to the inlet and exit ducts to limit thermal losses, to achieve a chamber temperature of ∼50°C. Automated fluidic handling was accomplished with a 4-channel peristaltic pump (Masterflex Ismatec Reglo MFLX78001-49), to dispense three reagents (PR2 buffer, Incorporation, and Cleavage) and for waste removal, plus two custom manifolds of pinch valves (BioChem Two-Way Normally Closed 24V). One manifold (with 9 valves) selects which well to send each reagent to, and the other (with 3 valves) which well to remove waste from. The valves are actuated by a custom 12-channel controller connected to the PC via USB (see custom part 2-0358). A flow sensor (Sensirion SLF3C-1300F) was connected to each fluidic channel, between the peristaltic pump and the pinch valves, to monitor the dispensed and removed volumes. A custom data acquisition board interfaces with the flow sensors (see custom part 2-0731) and connects to the PC via USB.

Custom, 3D printed caps covered each well on the plate and held the reagent dispensing and waste removal tubes into the well at the proper location and depth. The caps were held in place using a 3D-printed bar that attached magnetically to a set of 3D-printed parts that held down the well plate on the microscope stage (see custom assembly 2-0269). The fluidics lines were managed such that no strain or drift was introduced as the microscope stage moved during acquisition. Critical to the design of the fluidics system, the reagent ports are located at the front of the microscope and the waste toward the back. A permanent static tilt was introduced at the base of the microscope with the Thorlabs Pedestal Base (Thorlabs PB4) to facilitate full fluid aspiration from the wells. Tefzel (ETFE) material was used for the fluidics lines in the fluidics caps as it was empirically determined to have the lowest effect to the image quality (especially important is its low fluorescence). A fluidics schematic is included in Suppl. Fig. 1B.

Incorporation and Cleavage mixes were stored in 50 mL conical tubes, maintained at 4°C throughout sequencing via a chiller (Fisher Scientific Mini Dry Bath) and protected from light with a custom 3D-printed cover (see custom parts 3-1828 and 3-1829). PR2 buffer was stored at room temperature in a glass media bottle, and waste was collected in a glass media bottle.

A Bill of Materials (BOM), included in Supplementary Table 6, contains the custom and commercially-available parts to build the automated ISS system. All CAD assemblies and parts were designed in Onshape. The Onshape CAD document is available at the following link: https://cad.onshape.com/documents/1ca6b324eff8e4599cbfe823/w/3a490d4448cb5f087d5522f6/e/6b8c4ed1eb8b1dbfc0bd0568?renderMode=0&uiState=6a18c2e2a82e85f952c651c7.

All 3D printed parts were printed in-house on a Prusa XL and Formlabs Form3 SLA 3D printers, as specified in the BOM. All in-house laser cutting was done with a Universal Laser Systems PLS6.15D laser cutter.

The custom 12-channel valve controller and 4-channel flow sensor interface designs, drivers, and firmware are available on GitHub at the following link: https://github.com/czbiohub-sf/uxibxx-io-board.

### 7.3 Automated ISS Isothermal Protocol

With the cage incubator maintaining ∼50°C, the temperature inside each well during operation was ∼47°C throughout the protocol. As such, adjustments to the standard *in situ* sequencing protocol were made to accommodate isothermal operation. Each round of sequencing begins with removing the full well volume, followed by dispensing 750 µL of incorporation mix. The incorporation mix is incubated for 6 minutes. After the incubation, 1.25 mL of PR2 buffer is added and immediately after the addition 1.25 mL of the well volume is removed. This is repeated 7 times, followed by a full removal of the well volume. Five washes with 1.25 mL of PR2 buffer and a 5-minute incubation each follow. Finally, the full well volume is removed and 2 mL of PR2 buffer is added prior to imaging. After imaging the full well volume is removed, and 825 µL of cleavage mix is added and incubated for 60 minutes. After the incubation, 1.25 mL of PR2 buffer is added and immediately after the full well volume is removed, repeated twice. Three washes with 1.25 mL of PR2 buffer and a one-minute incubation each follow. The well volume is exchanged, and the well is ready for the next round of base addition.

### 7.4 Software and Operation

All standard microscope hardware (camera, microscope stage, filters, light source) is controlled via µManager^15,16^ via the Pycro-Manager Python library^17^. Auxiliary hardware (pumps, valves, and flow sensors) is controlled via serial commands sent through the Python control software. The specific protocol parameters (e.g. number of wells, sequencing depth, volumes, incubation times, etc.) are stored in a configuration file, allowing for independent control of the protocol for each well. The core of the control software is the scheduling logic and task executor. Using the Google OR-tools suite^22^, we model the protocol and resources (camera, pump channel, wells) as a classic constraint optimization scheduling problem with an objective of minimizing total runtime. The task executor runs the optimized schedule and manages resource contention and availability. A standard (3 well, 10 base sequence depth) protocol runs in <23 hours with an optimized schedule, as compared to ∼65.5 hours with sequentially scheduled tasks.

Operation of the instrument takes under 10 minutes of total hands-on time. The incubation chamber and chiller are turned on 1.5 hours prior to the sequencing run to allow for the full system to come to thermal equilibrium. The incorporation and cleavage reagents are extracted into 50 mL conical tubes and inserted into the dry chiller. The PR2 buffer is diluted with PBS-T to a final volume of ∼950 mL and added to the buffer media bottle. The 6 well sequencing plate is inserted into the stage holder and the 3 fluidics caps (see custom part 3-1573 in the BOM) are placed in each of the 3 wells for sequencing. A magnetic bar latches and secures the caps and plate in the stage insert (see custom parts 3-1574 and 3-1587). The user runs the software via command line, which interactively prompts for the image storage location, the objective position for the DAPI image focal plane, and two-point plate tilt calibration (center bottom of well A1, and center top of well A3). While all parameters for the experiment are configurable, standard experiment conditions are provided in the example protocol below. In addition, the standard exposure times are 50 ms for DAPI (which is only acquired in the first round of sequencing), 100 ms for MiSeqG, 100 ms for MiSeqT, 100 ms for MiSeqA, and 500 ms for MiSeqC. The typical sequencing depth is 10 bases, though up to 20 bases have been sequenced. Finally, sequencing of a series of plates back-to-back further reduces the total experiment time, reducing the amount of hands-off time required for thermal equilibrium of the system.

All software, written in Python, controlling the ISS workflow is available on GitHub at the following link: https://github.com/czbiohub-sf/OPSAutomationV1.

## 8. CROP-seq

### 8.1 CROP-seq screens

A549 cells stably expressing doxycycline-inducible Cas9 were transduced with the “1000-gene” library (4,211 sgRNAs). Library cells were thawed and expanded under puromycin selection at >1,000× library coverage, then seeded into 6-well plates at >250× starting cell coverage per sgRNA. Puromycin was withdrawn and doxycycline was added to induce Cas9 expression on the day of cell seeding. After the indicated duration of induction (see below), cells were harvested by trypsinization, pooled across wells within each condition, washed in PBS, and resuspended in PBS + 10% FBS at 1.5 × 10⁶ cells/mL. Single-cell libraries were prepared on the 10X Genomics Chromium GEM-X 3′ platform (#1000691) with 40,000 cells loaded per lane.

Cells were seeded at 90,000 cells per well at 13 days post-transduction and induced with doxycycline for 90 h. The 1,000-gene screen was performed in four independent batches, yielding approximately 150× cell coverage per sgRNA.

Gene-expression libraries were prepared following the standard 10x Genomics protocol. sgRNA cassettes were enriched from full-length cDNA by three sequential PCRs, each monitored by qPCR using the KAPA HiFi HotStart ReadyMix PCR kit (#50-196-5217) with an EvaGreen (#31000) spike-in to avoid overamplification, as previously described^23^. PCR1 was performed from 25 ng of cDNA with cycling at 95°C for 3 min, followed by cycles of 98°C for 20 s, 65°C for 15 s, and 72°C for 40 s, and a final extension at 72°C for 1 min. PCR2 used a 1:20 dilution of PCR1 product with cycling at 95°C for 3 min, followed by cycles of 98°C for 20 s, 65°C for 15 s, and 72°C for 20 s, and a final extension at 72°C for 1 min. PCR3 used a 1:25 dilution of PCR2 product and added Nextera-compatible sample indices unique to each 10x lane, with cycling at 95°C for 3 min, followed by cycles of 98°C for 20 s, 72°C for 15 s, and 72°C for 20 s, and a final extension at 72°C for 1 min. Final libraries were SPRI-purified, quantified by TapeStation, and pooled in equimolar ratios. Gene-expression libraries were sequenced on a NovaSeq X at an average depth of >60,000 reads per cell. sgRNA libraries were sequenced for 20 cycles on a MiSeq i100 using a custom sequencing primer (5′-TAAGAGACAGCTTGTGGAAAGGACGAAACACCG-3′) initiating at the first base of the sgRNA cassette.

### 8.2 CROP-seq alignment and processing

10X data were aligned using the standard count function with Cellranger version 9.0.1 and the human reference genome (primary assembly) GRCh38 version 47. Alignment was run with the --no-secondary tag. QC and analysis were conducted in Python 3.12 with scanpy version 1.12. Analysis functions and notebooks can be found at https://github.com/czbiohub-sf/cropseq1000/. Briefly, GEMs containing over 50 genes and 500 UMIs were read from Cellranger output matrices (‘raw_feature_bc_matrix’) from all batches into a single AnnData object. Barcode assignment from sgRNA sequencing was conducted by reading in cell, guide, and UMI information from matched FASTQ files and discarding reads without a matching cell barcode in the anndata object. After UMI deduplication, each cell was required to have two or more reads with matching guide barcodes; any cell with multiple different barcodes identified was labeled a multiplet unless one barcode had > 10× more reads (UMIs) than the next most common barcode for that cell.

Following barcode assignment, cells were filtered based on their cDNA UMI counts, unique gene counts, and percent mitochondrial content. Only cells with greater than 3,000 unique genes, 8,000 UMIs, and less than 10% mitochondrial content were kept for downstream processing. The gene and count thresholds were chosen based on the shape of the violin and scatter plots shown in Suppl. Fig. 7C to exclude any lower outliers while preserving all cells within the normal distribution of UMI and gene counts for all lanes. Ambient RNA removal was not run, as a trial run of DecontX removed ∼40% of all UMIs compared to the <5% ambient RNA detected by Cellranger (Fraction Reads in Cells metric ∼97-99% across all lanes).

Doublets were identified using the scrublet function within scanpy (scanpy.pp.scrublet) on each 10X lane. ¾ of the identified doublets overlapped with barcode-doublets (cells with more than one barcode identified). Additionally, cells in the top 0.5% of counts and genes per batch were identified as ‘likely multiplets’, over half of which contained multiple barcodes. All doublets identified through high UMI counts, Scrublet, and multiple barcode assignment were removed before downstream processing.

Following filtering, standard log-normalization was conducted with scanpy.pp.normalize_total(target_sum=10,000) and scanpy.pp.log1p. Cell cycle phase was calculated using the cell cycle genes provided in Kowalczyk et al.^24^ and the scanpy tl.score_genes_cell_cycle function.

Pseudo-bulking was conducted by summing raw counts across all cells carrying the same guide using the scanpy get.aggregate function, followed by log normalization, calculation of highly variable genes (‘seurat’ flavor; min_mean = 0.001, min_disp=0.1; 6,526 highly variable genes selected), scaling, PCA, neighbor calculation on the top 40 PCs, UMAP dimensionality reduction, and Leiden cluster calculation (resolution=1).

## 9. Other experimental protocols

### 9.1 siRNA and live-cell time course imaging

A549 cells tagged endogenously with NPM3 were seeded at a density of 5,000 cells/well in a 96 well glass-bottom plate. The next day, the media was swapped out to media with a transfection mix of DharmaFECT 1 transfection reagent (T-2001-01) with Non-targeting, CCT6A, TCP1, and CCT3 siRNA (Horizon Discovery, D2-001810-10, L2-016559-01, L2-012749-01 and L2-018339-01 respectively) for a final concentration of 25 nM in each well. The plate was incubated for 72 hours before the samples were fixed with a final concentration of 4% paraformaldehyde and washed thrice with PBS. The samples were then imaged with a Leica 63× 1.47 NA oil objective, 7 z-slices with step size of 1 µm. For the live-cell time course, cells with the CCT6A knocked down were live-imaged with the same 63× objective at 15-minute intervals, 3 z-slices with a step size of 2 µm starting from 1 hour post transfection.

### 9.2 RT-qPCR for pre-rRNA processing analysis

Knockdown was performed by seeding 7000 A549 cells per well in a 96-well plate in medium containing a transfection mix of DharmaFECT 1 transfection reagent (Horizon Discovery, T-2001-01) and non-targeting control (NTC), POLR1B, or CCT6A siRNA (Horizon Discovery; D2-001810-10, L2-013391-01, and L2-016559-01, respectively) at a final concentration of 25 nM. At 72 h post-transfection, RNA was reverse transcribed and amplified directly from cell lysates using the Power SYBR Green Cells-to-CT Kit (Thermo Fisher Scientific, A25600). Before reverse transcription, cells were washed once with cold PBS and lysed in Lysis Solution containing DNase I (5 min, room temperature), and lysis was stopped with Stop Solution (2 min). Each 20 µL reaction contained 1 µL cell lysate and 200 nM of each primer. Cycling conditions were 95°C for 10 min, followed by 40 cycles of 95°C for 15 s and 60°C for 1 min, and a melt-curve step to confirm a single product.

Pre-rRNA was quantified with a primer pair targeting the 5′ external transcribed spacer (5′-GTGCGTGTCAGGCGTTCT-3′ and 5′-GGGAGAGGAGCAGACGAG-3′), which spans the 01/A′ primary processing site (≈ +414) and is therefore specific to the unprocessed 47S primary transcript. GAPDH (5′-AGCCACATCGCTCAGACAC-3′ and 5′-GCCCAATACGACCAAATCCGTT-3′) served as the reference gene.

Relative pre-rRNA levels were calculated using the 2^−ΔΔCt^ method and expressed as fold-change relative to the paired NTC, set to 1 within each biological replicate. Data are from three independent experiments (separate cell seedings and siRNA transfections); within each experiment, the four wells per condition were averaged, and each sample was measured in two technical qPCR replicates. Data are presented as mean ± s.e.m. of n = 3 biological replicates. Significance was assessed by a two-tailed one-sample t test; p < 0.05 was considered significant. Analyses were performed in Python.

### 9.3. BODIPY staining of neutral lipids and confocal imaging

A549 cells harboring doxycycline-inducible Cas9 and constitutively expressed sgRNAs targeting EMC1, EMC2, or EMC3, or a non-targeting control (NTC) (GCCCTCCATCACCTCTCCAG, ATTGCCATTCGAAAAGCCCA, CACCAATAAGAATCATAGGG, and CGCCTAATTTCCGGATCAAT, respectively), were seeded separately at 15,000 cells per well in 24-well plates. Knockout was induced with 1 µg/mL doxycycline for 72 h, followed by a 12 h incubation in imaging medium containing BODIPY 493/503 (1:10,000). Cells were imaged on a Leica DMi8 microscope with an Andor Dragonfly spinning disk confocal module using a Leica 63×/1.47 NA oil-immersion objective. For each field of view, a single-channel z-stack of 11 planes was acquired at a 0.4 µm axial step.

### 9.4. Lipid-droplet quantification

Images were analyzed in Python using Cellpose v3.1.1.1 (cyto3 model^25^) and scikit-image. Z-stacks were maximum-intensity projected, and cells were segmented from the BODIPY signal with a fixed expected diameter of 22.7 µm (automatic estimation failed on these large, spread cells). Border-touching cells were excluded, and masks were extended into the peripheral cytoplasm (marker-seeded watershed, ≤3 µm) to capture edge droplets. Within each cell, diffuse cytoplasmic/ER signal was removed by white top-hat filtering; lipid droplets were then thresholded, separated by an intensity-maxima–seeded watershed, and retained if 0.03–4.0 µm² with solidity ≥ 0.8 and eccentricity ≤ 0.92. The number of lipid droplets per cell were quantified in 5 fields of view per genotype (NTC, n = 167 cells; EMC1 KO, n = 55; EMC2 KO, n = 82; EMC3 KO, n = 87). Each EMC knockout was compared with the NTC control by two-sided Mann– Whitney U test, with each cell treated as an independent observation; ***p < 0.001.

### 9.5 Lipidomics

The A549 EMC1, EMC2, EMC3 knockout and NTC lines described above were seeded at 80,000 cells per well in 6-well plates (three biological replicates per genotype). Knockout was induced with 1 µg/mL doxycycline for 72 h, followed by 12 h incubation in imaging medium before harvest. Culture supernatant was then aspirated, and plates were immediately placed on ice. Cells were washed twice with 2 mL ice-cold PBS, scraped into 1 mL ice-cold PBS, and transferred to microcentrifuge tubes. Cells were pelleted at 300 g for 5 min at 4°C, and the PBS was aspirated. Lipids were extracted by resuspending each cell pellet in 225 µL of LC-MS-grade methanol containing SPLASH Lipidomix internal standard (Avanti Polar Lipids, Cat# A83707). Samples were vortexed for 5 s and stored at -80°C until further processing.

Lipid extracts were generated as described previously^26^ and subsequently separated by reverse phase liquid chromatography (Thermo Scientific Dionex UltiMate 3000) and coupled to either a Thermo Fusion Lumos (positive ionization mode) or a Thermo qExactive HF (negative ionization mode). Data dependent tandem mass spectra (MS2) were acquired and fragmentation data were used in lipid annotation. Full scan (MS1) spectral resolution was 60k and MS2 resolution was 15k in negative ionization mode and 30k in positive ionization mode. Quality control was performed by comparison of SPLASH Lipidomix peak intensity and shape across all samples (an EMC2 KO replicate was removed due to failing QC in positive ion mode). Peak picking, annotation and alignment across samples were performed by MS-DIAL v4.60^27^. MS1 noise threshold was set to 50k peak height. Lipid features were required to be found in 100% of at least one set of biological replicates to be retained in the final result. All lipid features were required to have a maximum sample intensity 5× greater than the mean signal in all blank samples.

## 10. Reference databases

### 10.1 Protein complexes definition

Curated human protein complexes were retrieved from the EBI Complex Portal^28^ via the IntAct REST API (accessed [2026-05-14]). Predicted entries were discarded, and each remaining complex was annotated with its ECO confidence score (1–5 stars) from the detail endpoint; only complexes with a score ≥ 3 (i.e., at least “inferred by curator”, ECO:0005547) were retained. Protein interactors were parsed to base UniProt accessions (feature-chain and isoform suffixes stripped), and complexes with fewer than two protein members were removed. The retained complexes were intersected with the OPS perturbation gene panel. Each OPS gene was then assigned to its smallest containing complex (ties broken alphabetically by Complex Portal accession), and complexes retaining fewer than two uniquely assigned OPS genes were dropped. The complete set of retained protein complexes is provided in Supplementary Table 5.

### 10.2 Gene essentiality scoring

A549 CERES gene effect score was downloaded from the DepMap Portal on March 27, 2025.

### 10.3 Pathway and molecular-function gene set definition

Human KEGG pathway maps and their gene memberships were retrieved from the KEGG REST API^29^ (accessed [2026-08-05]); disease maps (hsa05*), global/overview maps (hsa01*) and maps of more than 200 genes were discarded, and KEGG gene identifiers were resolved to panel symbols through the KEGG human gene list. GO molecular-function annotations were taken from the UniProt-GOA human GAF^30^ (accessed [2026-06-17]); only aspect-F annotations carrying experimental evidence (EXP/IDA/IPI/IMP/IGI/IEP) and no NOT qualifier were used, propagated to all ancestor terms, and terms covering ≥ 2% of experimentally annotated human genes were dropped as too broad. Both sources were intersected with the 1,000-gene panel, retaining candidates with 2–60 panel genes. Within each source, a candidate with Jaccard ≥ 0.7 to a more specific retained candidate was dropped as a near-duplicate or nested set; each gene was then assigned to its single most specific containing set (GO by information content, KEGG by map size; ties broken by database identifier), and sets retaining fewer than three uniquely assigned genes were dropped and their genes reassigned until stable, giving 56 KEGG and 51 GO sets. The two partitions were pooled without further filtering, so a process annotated in both databases contributes one set from each.

## 11. Data analysis – image processing

Three co-acquired imaging modalities are processed for each experiment: 20× volumetric brightfield and fluorescence for phenotyping (7 z-slices), 5× brightfield for live-cell tracking (9 z-slices), and 5× fluorescence multi-cycle in situ sequencing (ISS). Each modality is independently preprocessed, then registered so that cells can be tracked and single-cell perturbations assigned in the final assembled dataset (Suppl. Fig. 6A). All raw and intermediate data are stored as OME-Zarr^31^ in an HCS plate layout. All code related to image preprocessing and assembly of the final dataset is publicly available at github.com/czbiohub-sf/cyclops-process.

### 11.1 Image ingestion, organization, and corrections

All ISS rounds are converted from OME-TIFF into a single OME-Zarr store with rounds along the time axis, and 5 channels (4 ISS channels, G,T,A,C + DAPI). Tracking and phenotyping data are reorganized into the HCS layout with per-channel geometric transforms applied as needed. Voxel scales are (z, y, x) = (1.0, 1.3, 1.3) µm for ISS, (25.0, 1.3, 1.3) µm for live-cell tracking and (2.0, 0.325, 0.325) µm for phenotyping; individual tiles are 2048 × 2048 pixels.

Lens pin-cushion distortion in the 5× brightfield is corrected per z-slice by applying a pre-calibrated radial polynomial *r → fact(r) = Σ cᵢ·rⁱ* to image coordinates. For fluorescence phenotyping channels, a multiplicative illumination flatfield is estimated as the Gaussian-smoothed (σ = 75 px) mean of 500 sampled positions, max-normalized and clamped to [0.2, 1.0]; a 100-count camera offset is subtracted before division and re-added afterward. For ISS, inter-cycle XY drift is estimated by phase-cross-correlation image stabilization on summed-channel stacks, averaged over 5 tiles sampled from different regions of the well, and applied with subpixel shifts to all tiles.

### 11.2 Tile Stitching

Each modality is stitched into a per-well composite using an established efficient multidimensional big-data stitching algorithm^32^. For 5× tracking and 20× phenotyping, tile positions are estimated from the 2D phase-reconstruction channel, while for ISS they are estimated from the DAPI channel. Briefly, pairwise tile offsets are estimated by phase-cross-correlation for all overlapping tile pairs (8% overlap between neighboring tiles). A linear system is then solved to find the optimal position of all tiles in the well, minimizing the deviation from all pairwise estimated shifts. Euclidean-distance-transform (EDT) blending is used in the overlap regions, weighting each pixel by its distance to the nearest tile edge for smooth blending between tiles. For tracking and ISS timepoints, a single shift map is reused across all timepoints. Final image sizes are roughly: ≈100k × 100k pixels for 20× phenotyping, and ≈13k × 13k pixels for ISS (per cycle) and 5× tracking (per timepoint).

### 11.3 ISS spot detection and base calling

Spots are detected on the stitched and cycle-registered ISS volume with the Spotiflow^33^ general pretrained deep-learning model. Detection is performed independently on each of the four round-0 sequencing channels (G, T, A, C). The per-channel spot lists are concatenated and de-duplicated, assuming that spots within a 5-pixel cutoff are duplicates, to yield a single point cloud per well.

The ISS basecalling algorithm is based on the analysis from Feldman et al.^10^, code available at (https://github.com/blaineylab/opticalpooledscreens). Briefly, for each spot, a (cycle × channel) intensity tensor is sampled from the registered volume and equalized by dividing each spot’s per-channel value by the channel-wise median across all spots in the tile. A within-tile spectral-crosstalk correction is then estimated for each channel, and applied to every spot’s intensity tensor to deconvolve channel bleed-through. The base call per cycle is the argmax over the four corrected channels, yielding an A/C/G/T barcode over the ten rounds.

### 11.4 Phase reconstruction

The brightfield z-stack was reconstructed to a 3D and a 2D phase estimate using WaveOrder^34^, a collection of forward and inverse filters for estimating physical parameters from raw microscopy data. The forward transfer function was computed for both acquisition geometries, and inverse filters were computed using a Tikhonov regularized inverse with a regularization strength of 1e-3. For 3D reconstructions, the raw data was padded with 5 slices of zeros to avoid axial wraparound artifacts.

For 2D phase reconstructions, the in-focus z-plane is a critical parameter that can change across a single field-of-view (Suppl. Fig. 4A, i). To estimate this parameter, we split each field of view into a grid of overlapping “subtiles” (4x4 grid for 5× tracking data, 16x16 grid for 20× phenotyping data, 25-pixel overlap), found the 3D reconstruction’s z-plane with the largest mid-band spatial frequency, and used this plane as the in-focus slice to complete the 2D reconstruction. After reconstructing each subtile, we blended and stitched the subtiles back into a single 2D reconstruction.

We found these nominal reconstructions to have high contrast near the well center, but near edges the contrast degraded (Suppl. Fig. 4B, iii), likely due to the air-media meniscus tilting and vignetting the illumination light (Suppl. Fig. 4A, i).

### 11.5 Optimized tilt and defocus reconstruction

We extended the nominal reconstruction with three scalar parameters per subtile: zenith tilt angle, azimuth tilt angle, and defocus. We split each FOV into an 8x8 grid of 50-pixel-overlap subtiles, and we estimated these three parameters for each individual subtile by maximizing the mid-band power of the 2D phase reconstruction (Suppl. Fig. 4A, ii). Optimization used 10 iterations of PyTorch NAdam with autograd and parameter-specific learning rates (0.005 zenith, 0.01 azimuth, 0.05 defocus). The optimized parameters were then used to create the final reconstruction (Suppl. Fig. 4B, iv).

The optimization is sensitive to initial conditions, particularly at the well edge where a non-tilted initialization is outside the basin of attraction of the optimizer. We mitigated this with a sparse chained optimization, performing an optimization at the center of the well, then using the result to initialize the next optimization along each N, S, E, W direction. We ran 29 and 13 optimization steps for 20× and 5×, respectively, then used the results to initialize with a four-parameter fit. Our initialization fit was of the form a*exp(b*r) + c for the zenith initialization (three parameters) and a constant for the defocus (one parameter). We initialized the azimuth angle with a parameter-free arctan(y/x).

### 11.6 Virtual Staining

Nuclear and membrane signal were predicted from 3D phase reconstructed images using Viscy^35^, a deep learning-based virtual staining model. Separate models were trained for the 20× phenotyping 3D phase reconstruction, and the 5x tracking 2D phase reconstruction.

### 11.7 Nuclear and Cell Segmentation

Nuclei were segmented with Cellpose-SAM^36^ from the DAPI channel for ISS and from the virtually stained nuclei in 20× phenotyping and 5× tracking images (Cellpose parameters: diameter = 30, flow_threshold = 0.7). Nuclei images were first preprocessed with a CLAHE filter (clip_limit = 0.01) to normalize contrast across the image. For ISS only, the nuclear segmentation is then dilated (8 pixels), in order to capture ISS spots that are located at the edge of the nucleus.

To increase processing speed and efficiency, segmentation is performed on the original microscope acquisition tiles (2048 x 2048). Segmentations are then stitched together using the stitching parameters estimated from the phase images (described in 11.2). Rather than blend the overlap region, segmentations are matched using an IoU ≥ 0.1, and relabeled.

Cells were segmented only for 20× phenotyping from the virtually stained nuclei and membrane channels, also using Cellpose-SAM (diameter = 100).

### 11.8 Channel Registration

Per-channel affine transforms are estimated automatically per experiment. Each fluorescence channel is first seeded with a median affine (scale and rotation only), derived from prior experiments. Starting from this seed, the residual affine — correcting for per-experiment translation/offset drift — is estimated using Mattes mutual-information registration on Gaussian low-pass filtered images, which is robust to the inverse intensity correlation between fluorescence and phase signal. Each tile’s candidate transform, together with the unmodified seed as a baseline, is cross-validated by scoring mean normalized mutual information (NMI) against phase across all tiles; the candidate with the highest cross-tile NMI is selected.

### 11.9 Cross-Microscope Registration

ISS, tracking, and phenotyping images, are registered so that base-calls can be tracked to phenotyping images. These registrations are computed automatically, per well from the nuclei segmentations. The 20x phenotyping segmentation is first down-sampled 4x so that it matches the resolution of the 5x live-cell tracking and 5x ISS segmentations. A coarse 2D phase-correlation translation is then estimated, after which source and target cell centroids are matched by Hu-moment shape similarity within a local k-neighbor graph. A 2D similarity transform (rotation + isotropic scale + translation) is then fit iteratively to the matching pairs using a RANSAC algorithm. The transform is then applied and the matching, and transform fitting steps are repeated to refine the final registration. The precision of this registration has a significant effect on the final tracking accuracy.

### 11.10 Cell Tracking

Cells were tracked across live phenotyping imaging, low-magnification tracking with brightfield, and in situ sequencing, using ultrack^37^. Briefly, nuclei segmentations from each registered timepoint are assembled into nodes of a graph. Edges are added connecting the nearest cells in neighboring timepoints (up to 5 neighbors within a distance limit of 25 pixels). This graph is then used along with biological constraints (i.e., cells only divide into 2 daughters, cells do not merge, etc.) to formulate an ILP, which is then solved using Gurobi to get the optimized cell lineages. The final lineage graph is saved in the GEFF Zarr format.

## 12. Data analysis – quantification and statistical analysis

### 12.1 Deep learning inference

Single-cell crops centered on the cell’s bounding box were embedded using the following deep learning image extractors (code available at github.com/czbiohub-sf/cyclops-model):

- DynaCLR^38^ is a self-supervised contrastive learning framework for learning single-cell representation from microscopy, drawing positive pairs from the same cell. Here we trained an OPS DynaCLR model in-house, encoding multiple markers via a bag-of-channels approach with the ConvNext-Tiny backbone and a 256-d projection head. The model was trained using the NT-Xent loss at a temperature of 0.1, a per-GPU batch size of 512 across 4 GPUs (effective batch size 2048, distributed data-parallel), the AdamW optimizer at a learning rate 2×10⁻⁴, with a 10-epoch warmup followed by cosine decay. The augmentations used were random rotation, scaling, horizontal and vertical flips, contrast and intensity jitter, Gaussian smoothing, and Gaussian noise. For training and inference, we used 128×128 cell crops, z-score-normalized, producing 768-d embeddings.
- SubCell^39^ a masked-auto-encoder (MAE) ViT foundation model for fluorescence microscopy, trained on the Human Protein Atlas. We used the released DNA+Protein variant (‘DNA-Protein_MAE-CellS-ProtS-Pool’), a ViT-B/16 encoder followed by a gated-attention pooling head. Subcell takes two-channel inputs, so for each OPS, two-channel inputs (channel 0 = virtual staining nuclei, channel 1 = protein-of-interest) were bilinearly resized to 448 × 448 and per-channel min– max normalized to the range [0, 1]. We used the gated-attention pooler’s output (1536-d) over the final-layer patch tokens for cell embeddings.
- DINOv3^40^: the publicly released DINOv3 self-supervised ViT pretrained on the LVD-1689M curated dataset (1.7B images from the web), was used to embed individual OPS channels. Each OPS cell crop was embedded using ViT-L/16 (pre-trained with 3-channel input). Each single OPS channel was replicated across the three input channels, normalized, and scaled by ImageNet channel standard deviations, masked by the cell-segmentation, and resized to 256 × 256. We obtain the per-cell embedding as the normalized [CLS] token with 1024-d from the final transformer block.
- Cell-DINO^41^: a DINO-v2 ViT-L/16 pretrained on cellular imaging data. Single-channel OPS crops (160x160 pixels, centered around the nucleus) were z-score normalized. For Cell-DINO embeddings, the cells were not masked by their segmentation. The cell crops were then embedded with the channel-adaptive (‘channel_adaptive_dino_vitl16_pretrain_cells’) model. The per-cell and channel embedding is the normalized [CLS]-token (1024-d) extraction protocol described above for DINO.

All embeddings were then post-processed to a compact final embedding following the procedure described below, and representation quality quantified by mAP scoring (below).

### 12.2 Image Embedding

Single-cell features were represented by a Cell-DINO embedding vector. Within each experiment, each feature vector was centered to zero mean and scaled to unit variance, removing plate- and batch-level offsets prior to cross-experiment pooling. Cells from all experiments sharing the same biological reporter (e.g. EEA1 for early endosomes) were pooled, and a PCA was fit (on a maximum of 5 × 10⁶ subsampled cells, then all cells projected through this basis). The top components capturing 80% of cumulative variance were retained. For each marker the PCA features were then averaged across all cells sharing the same sgRNA. For each sgRNA, the embeddings for each marker were then concatenated into a single vector. The mean of non-targeting-control (NTC) guides was then subtracted from every feature and each feature divided by the standard deviation of the NTCs, so the origin represents the unperturbed state, and each axis encodes deviation from it. A second PCA was fit on the concatenated, NTC-normalized guide matrix, this time retaining the top components capturing 40% of cumulative variance to produce a compact final embedding. Gene-level embeddings were obtained by averaging across guides targeting the same gene (Suppl. Fig. 6B).

### 12.3 Organelle Segmentation

Intracellular organelles are segmented from the phenotyping composite by OrganelleProfiler (code available at github.com/czbiohub-sf/organelle-profiler), an adaptation of Nellie^42^, with the algorithm chosen per organelle class according to morphology. Tubular structures (mitochondria, endoplasmic reticulum) are segmented by multi-scale Frangi vesselness filters^43^. Vesicular structures (lysosomes, endosomes, lipid droplets) are detected by Laplacian-of-Gaussian blob detection. Nucleoli are segmented using Laplacian-of-Gaussian blob detection inside the nuclear mask. All organelle masks are written as named OME-NGFF label groups in the v3 composite store alongside the nuclear and whole-cell labels.

### 12.4 Cell and Organelle feature extraction, and volcano plot analysis

For each cell, CellProfiler (using cp_measure^44^) and OrganelleProfiler compute a per-object feature vector covering morphology, intensity, texture, network topology, and subcellular localization. For tubular organelles, the segmentation is skeletonized and analyzed as a graph. Subcellular localization features are computed relative to the cell and nuclear segmentation boundaries. Per-organelle features are aggregated to the cell level using seven summary functions (sum, mean, median, std, min, max, count) and then pooled to per-guide and per-gene levels; outputs are stored as AnnData objects.

Statistical significance for the volcano plots was computed using the guide-level bootstrap procedure described previously^7^. For each feature, a guide-level score was defined as the median value across all cells assigned to that guide RNA, and a gene-level score as the median across the four guide-level scores for that gene. Null distributions were constructed from non-targeting control (NTC) cells. For each distinct cell count n (capped at 5,000) observed among guides, a guide-level null distribution was generated from 10,000 replicates; in each replicate, one NTC guide was drawn at random from the pooled NTC guides, n cells were sampled from that guide with replacement, and their median was recorded. The gene-level null was assembled the same way as the gene-level score: in each of 10,000 replicates, one value was drawn independently from each of the gene’s four guide-level nulls (matched by cell count), and the median was taken. Two-tailed p-values were computed as twice the smaller of the fractions of gene-level null values exceeding or falling below the measured gene-level score. p-values were corrected across all genes using the Benjamini–Hochberg procedure to obtain FDR q-values.

### 12.5 Gene-KO and protein-complex mean Average Precision (mAP)

Phenotypic signal was quantified using copairs mean Average Precision (mAP)^45^. For each test, positive pairs and negative pairs were defined as follows, and AP was computed per query embedding using cosine distance:

Gene-KO mAP — positives: same perturbation, different sgRNAs; negatives: sgRNAs from different perturbations. This metric measures how embeddings from sgRNAs targeting the same gene rank with respect to each other vs. unrelated sgRNAs. It represents the degree to which sgRNAs targeting the same gene can be resolved from unrelated perturbations.

Protein-complex mAP — for each protein complex (EBI Complex Portal, with ≥3 active members), positives: same complex membership, different perturbation; negatives: not in complex. This metric measures how embeddings from knockouts of genes for subunits of the same protein complex rank with respect to each other vs. unrelated knockouts. It represents the degree to which knockouts targeting the same complex can be resolved from unrelated perturbations.

For each test, a null distribution was estimated by sampling 1 × 10⁶ random pair configurations (null_size=1e6, seed=0) and a p-value computed against threshold=0.05, then Bonferroni-corrected across queries. A perturbation/complex is called “distinctive” or “consistent” when its corrected p-value < 0.05.

### 12.6 Clustering of image markers

For each of the 56 imaging markers (including phase), we computed mAP against each of the 1,000 gene knockouts in the screen, yielding a marker × geneKO distinctiveness matrix M. To emphasize each marker’s diagnostic knockouts and downweight knockouts resolved by many markers, M was TF-IDF reweighted:

TF: for each marker, term frequency (TF) was defined as its row-normalized mAP, each value divided by the marker’s row sum. Every marker’s distribution sums to 1, removing differences in overall mAP magnitude.

IDF: for each knockout, document frequency (DF) was defined as the number of markers whose mAP for that knockout exceeded the column mean across markers. This per-knockout adaptive threshold was chosen because mAP values span a wide dynamic range across knockouts. IDF = log((N + 1) / (DF + 1)), with Laplace smoothing.

The TF-IDF-weighted matrix was then used to compute pairwise Pearson correlations between markers. The resulting 56 × 56 correlation matrix was hierarchically clustered with average linkage on 1 − r and rendered as a symmetric heatmap.

### 12.7 Marker Cluster Driver-perturbation Analysis (Figure 4C)

Five marker clusters were identified from the hierarchical clustering of the marker × marker Pearson correlation matrix in Figure 4C: (i) Mitochondrial + tubulin markers (n = 5): ChromaLIVE 561, CellROX, TOMM70, Tubulin (cp), TOMM20 (cp); (ii) Endo-lysosomes markers (n = 5): EEA1, VPS35, RAB7A, TFRC, LAMP1; (iii) Cytoskeleton (n = 3): FastAct-SPY555, VIM, Phalloidin (cp); (iv) ER/Golgi markers (n = 2): SEC23A, COPE; (v) Nucleus & nucleolus markers (n = 8): NPM1 (cp), FBL, H2BC21, LMNB1, MKI67, NPM3, SRRM2, PSMB7.

To determine which knockouts most specifically drove the phenotypic signal of each marker cluster, we calculated a specificity metric analogous to Cohen’s d. For every cluster-knockout combination, we compared the knockout’s average TF-IDF-weighted mAP across markers belonging to the cluster with its average across all remaining markers and normalized this difference by the pooled standard deviation of the two marker sets. A larger score therefore reflects a phenotypic distinctiveness signal that is concentrated within the cluster’s markers rather than broadly shared across the panel. This analysis included all 1,000 perturbations in the screen and all 56 markers, including phase. For each cluster, the 50 knockouts with the highest specificity scores were identified as drivers.

For each marker cluster c and KO gene g, a pooled-variance Cohen’s d analog was computed over the TF-IDF distinctiveness matrix X as:

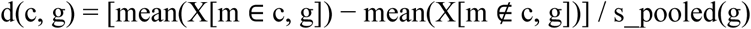

with s_pooled = √[((n_in − 1) σ²_in + (n_out − 1) σ²_out) / (n_in + n_out − 2)], where σ_in and σ_out are sample standard deviations (delta degrees of freedom = 1) across in- and out-of-cluster markers, respectively, and n_in + n_out = 56. The score is scale-free and ranks genes by their cluster-vs-background mean shift in pooled-SD units.

Top-50 per-cluster driver lists (ranked by Cohen’s d) were submitted to over-representation analysis via Enrichr^46^ through gseapy 1.2.1^47^. Five libraries were queried: GO Biological Process 2023, GO Cellular Component 2023, GO Molecular Function 2023, Reactome 2022, and KEGG 2021 Human. The 1,000-gene screen library, instead of the human genome, was used as the background set.

### 12.8 PHATE embeddings

Two-dimensional PHATE^48^ embeddings were calculated using the phate python library (1.2.1) with knn=10, decay=15 (image) or decay=7 (RNA), metric = ‘euclidean’ (image) or ‘cosine’ (RNA) and otherwise standard parameters. The input for image PHATE embeddings were PCA-reduced Cell-DINO embeddings (see above). The input for RNA PHATE embeddings was the perturbation effect matrix from sVAE+.

### 12.9 Unsupervised clustering and ontology enrichment

Leiden clustering was performed on both imaging and RNA perturbation embeddings. First a neighborhood graph was calculated using scanpy’s^49^ pp.neighbors function with n_neighbors=8, metric=’cosine’. Then leiden clustering was performed using scanpy’s tl.leiden function varying resolution parameters and flavor=’igraph’. For subsequent annotations resolution = 29 was chosen.

### 12.10 Ontology enrichment of modality-preferential genes

To identify functional categories preferentially recovered by imaging versus CROP-seq, a per-perturbation preference score was calculated as

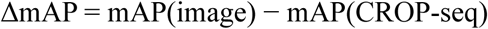

where positive values flag genes whose perturbation phenotype is better resolved by the imaging embedding and negative values are better resolved by the RNA embedding. Gene-set enrichment analysis (GSEA) was performed using gseapy 1.2.1^47^ with 1,000 permutations against the GO Biological Process 2025 and GO Cellular Component 2025 Enrichr libraries^46^. For each library, terms were ranked by FDR and closely related terms (duplicates) were removed. The top five terms for each ontology library and enrichment direction were plotted.

### 12.11 Adjusted Rand Index comparison of Imaging and RNA

To investigate whether phase imaging optical pooled screens allow us to gain comparable unsupervised biological insights to a current gold-standard CROP-seq screen, Leiden clusters were compared between the modalities using the Adjusted Rand Index (ARI).

Because not every perturbation causes a strong biological phenotype we do not expect concordant clustering across all 1,000 gene knockouts. Therefore, for each of the 186 clusters obtained by Leiden clustering of the RNA modality, an over-representation test against five Enrichr^46^ libraries (GO Biological Process 2025, GO Cellular Component 2025, Reactome 2022, COMPARTMENTS Curated 2025, and WikiPathways 2024 Human) was performed using gseapy 1.2.1^47^. For each cluster, significantly enriched terms across the five libraries, along with gene names and potentially confounding covariates such as chromosome arm location of each gene were written to a CSV file. To obtain a final annotation a large language model (Claude 4.7 Opus xHigh) was tasked to combine this information into a consensus functional label and a confidence score (1–5). We subsequently subset to 37 clusters with confidence score 5 containing 267 perturbed genes, representing confident biological groupings that can be identified from the RNA modality.

To obtain comparable Leiden clusters for the imaging modality Leiden resolutions from 25 to 35 were swept, and the resolution maximizing the ARI was selected. The ARI was calculated comparing the subset of clusters that overlapped the 267 genes with confidence score 5 in the RNA modality. The ARI has an expected value of 0 under the null hypothesis of random assignment and 1 with perfect concordance. To obtain an empirical p-value, 1000 random permutations of cluster labels were performed. The resulting p-value was 0.001, the lowest possible value at that number of permutations.

Finally, a cluster-by-cluster confusion matrix of RNA × image cluster labels was built, restricted to cluster pairs containing ≥ 2 shared genes. Rows and columns dropped by this overlap filter were summarized in a “catch-all” column and row whose color encodes per-cluster coverage (count / cluster total) (see Supplementary Figure 9A).

### 12.12 Modeling CROP-seq Data Using sVAE+

Raw count matrices were used as input. Each cell’s perturbation identity served as the condition label for the sVAE+^50^ structured prior, while batch assignment and cell cycle phase (G1, G2M, S) were included as categorical covariates. Non-targeting control cells (n = 45,320) were subsampled without replacement to match the cell count of the largest gene perturbation group (n = 1,196), preventing the control class from dominating the learned prior. Perturbations represented by fewer than 50 cells were excluded, retaining all 1,001 conditions. Genes were restricted to a pre-computed set of 6,526 highly variable genes identified from pseudo-bulk expression profiles. The final filtered dataset comprised 561,951 cells (down-sampling NTC cells to avoid over-representation) across 6,526 genes and 1,001 perturbation conditions.

Hyperparameters were selected via a 120-trial grid search over the following ranges: number of encoder/decoder layers in {2, 3, 4}, hidden layer width in {128, 256}, sparse mask penalty in {1, 2, 5, 20, 100}, KL warmup epochs in {50, 120, 200, 400}, with dropout rate fixed at 0.05. Each trial was trained for 50 epochs with early stopping, a 90/10 train/validation split, and a batch size of 4,096. The best configuration was selected by minimum validation ELBO (reconstruction loss + KL divergence). Post-sweep evaluation additionally assessed cluster compactness and mean Average Precision (mAP) on known protein complex annotations to confirm biological plausibility of the top configurations.

The best hyperparameters (n_layers = 2, n_hidden = 256, sparse_mask_penalty = 1, KL warmup = 200 epochs) were used for two-phase final training. In Phase 1, the model was trained for 300 epochs (batch size 4,096, early stopping, 90/10 split) with Gumbel-Sigmoid mask learning enabled, allowing the model to discover which latent dimensions each perturbation affects. In Phase 2, mask probabilities were binarized at a threshold of 0.5, the mask was frozen, global KL was disabled, and the model was retrained for 100 epochs (batch size 1,024, KL warmup = 1 epoch) to refine the action means and decoder under the fixed sparsity structure. The final trained model produces three outputs: (1) a perturbation effect matrix W (1,001 perturbations x 50 latent dimensions), computed as the element-wise product of the learned action means and the binary mask, (2) cell-level latent representations (561,951 x 50), and (3) guide-level latent representations obtained by averaging cell latent representations per sgRNA (4,206 x 50).

sVAE+ was benchmarked against three alternative dimensionality reduction approaches using the same mAP evaluation framework. PCA was applied to the 6,526-HVG expression matrix at the cell level (50 components, no batch correction), with cell-level PCs averaged per sgRNA for guide-level metrics and further averaged per perturbation for perturbation-level metrics. Consensus non-negative matrix factorization (cNMF^51^) was fit on DESeq2^52^ perturbation-level log2 fold-changes (20 factors), producing a perturbation usage matrix; only 4 perturbation-level mAP metrics were computed, as the DESeq2-derived NMF has no faithful guide-level projection. ContrastiveVI^53^ was trained with 10 salient latent dimensions using the background/target split to separate perturbation-specific variation from shared variation; salient cell-level representations were averaged per sgRNA and per perturbation. For all methods, the same set of perturbations, protein complex annotations, and permutation testing procedure were used to ensure a fair comparison.

## 13. Attention classifier

### 13.1 Single-Cell Embeddings

Single-cell embeddings of dimension 1,024 were extracted from Cell-DINO^41^ for each channel of data (either label-free phase imaging or a fluorescence marker). Embeddings were grouped by experiment and by channel; within each group, each embedding dimension was z-standardized using the mean and standard deviation of the non-targeting control (NTC) cells. Separate classification models were trained on (1) cells obtained from label-free imaging and (2) cells obtained from fluorescence imaging, spanning different fluorescence markers. Label-free imaging had ∼65M single-cell embeddings (median 61,000 per gene knockout). Across 55 markers, fluorescence imaging had ∼86M single-cell embeddings when separately counting individual channels in multiplex-labeled fixed cell images (median 1,300 per gene knockout, per marker). In each case, 80% of the embeddings were randomly assigned to the training set and the other 20% were reserved as a hold-out set for validation.

### 13.2 Set Classifier

A neural network classification model was trained to input a set of cells receiving the same perturbation and predict the perturbation class from one of *C* labels. Within the machine learning literature, such a problem formulation is known as multiple instance learning^54^. Separate models were trained depending on the particular definition of “perturbation class”; this could either be fine-grained (e.g. *C* = 1,001 classes containing 1,000 genes in the knockout library plus NTC), or more coarse-grained (e.g. *C =* 99 protein complex classes that each group together multiple genes) – depending on whether the desired objective is to understand the uniqueness of each gene knockout or each complex. The model architecture was based on a Set Transformer^55^, which enforces permutation invariance (i.e. reordering of cells within the set does not change the final prediction) and enables multi-scale inference (i.e. the model can return a prediction regardless of set size). Within each set, cell embeddings were concatenated with a learnable channel embedding signifying each cell’s channel identity and then passed through a linear projection to yield a vector of *d=*512 dimensions per cell. The set of channel-aware embeddings was then passed through *l=2* layers of inducing-point set attention blocks (ISAB)^55^ with *p*=32 inducing points. Compared to standard attention blocks, ISAB has computational complexity that scales linearly with set size, allowing the model to efficiently process sets containing up to tens of thousands of cells. The model employed *h=4* attention heads per layer and a feedforward dimension of *f = 4 * d* = 2048. The final layers of the model consisted of a pooling by multi-head attention (PMA) layer^55^ that collapses the set of embeddings to a single embedding of dimension *d*, followed by a cosine classifier^56^ that yields a probability distribution over class labels.

During model training, each training example was a set of *n=*100 cells receiving the same perturbation, sampled randomly from the pool of cells for that perturbation. A training epoch consisted of iterating through *s=*32 sets for each perturbation. Each model was trained for 200 epochs with 64 sets per batch, 0.0001 peak learning rate, and 20 epochs of linear warmup followed by cosine annealing. Even though the model was consistently trained with *n=*100 cells per set, it is possible for the set classifier to provide predictions for any value of *n*. As shown in Figure 3B, after the model was trained, it was evaluated on different values of *n* (ranging from 10 to 5,000) on the validation set.

### 13.3 Classification-Based Ranking of Single Cells

After training the classifier, a removal-based interpretability technique^57^ was used to identify the cells that have the greatest influence on the classifier’s predictions. For each knockout/complex, these “top-predictive cells” enable the classifier to accurately identify that particular knockout/complex from the pool of all classes, and therefore contain a phenotypic signature of that class. To find the top-predictive cells, the following scoring mechanism was employed: given a cell x with perturbation class c, we constructed a bag *B_n_* of n cells containing x and n-1 randomly sampled cells from the same class. The score of x in the context of *B_n_* was defined as the difference between the classifier’s output probability p of the correct label when x is present in the bag and when x is removed, i.e.

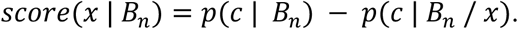

This quantity captures the marginal value of x to the specific bag *B_n_*. To construct a global score for x across different contexts, we take an average over many different bags, varying bag size and the partners within the bag, i.e.

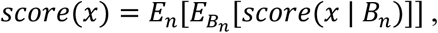

where the outer expectation *E_n_* denotes an averaging over various bag sizes n=1, 2, 5, 10, 20, 50, 100, 200, 500 (chosen to be roughly evenly-spaced on a logarithmic scale) and the inner expectation *E_Bn_* denotes an averaging over bags that contain x for each size n. Since the inner expectation is expensive to compute exactly over all possible bags containing x, we employed a Monte Carlo approximation with 100 random samples. For each perturbation class and marker combination, scores were calculated for all cells. The top-scoring ones, termed “top-predictive cells”, were used for downstream visualization and further analysis.

### 13.4 Top-Predictive Cells Distill Phenotypic Distinctiveness

The Cell-DINO mAP scores from phase cells (n=65M cells) (Fig. 3D) were recomputed using only the top-predictive cells per perturbation, using the top 20,000 cells per gene for phase, and top 500 for each of the 55 fluorescence channels — with weights normalized to mean=1 per (sgRNA, experiment) group. Rankings for gene-level mAP use the Set Transformer trained to classify 1,000 gene KOs; rankings for protein-complex mAP use the Set Transformer trained to classify 99 protein complex classes. The rank-derived score is 1 − rank / N, where N is the number of cells for that perturbation, so higher-ranked cells contribute more. Post-weighting, the same PCA post-processing method was applied (see Supplementary Figure 6).

### 13.5 Generative single-cell counterfactual traversals

#### Generative model

A diffusion autoencoder^58^ was trained on single-cell images (label-free phase, or individual fluorescent markers). Each cell is represented by two disentangled latents: a 1,024-dimensional semantic embedding obtained from Cell-DINO, and a stochastic (noise) latent that captures the residual identity of the individual cell — its local context. The decoder is a conditional diffusion model that reconstructs the cell image from the semantic embedding using classifier-free guidance (guidance scale w = 1.5, with a learned null embedding). Because phenotype and identity are encoded separately, a given cell can be re-synthesized while its phenotype is smoothly and independently varied.

#### Perturbation direction and counterfactual traversal

For each gene knockout, a perturbation direction was defined in the 1,024-dimensional Cell-DINO embedding space using the top-predictive 1,000 cells of the knockout and NTC populations, as described above. A linear (logistic) classifier separating the two populations defines the unit direction *d* pointing from the control toward the knockout state (i.e., the phenotypic trajectory). We scale this direction by the centroid gap, the distance between the average NTC and average knockout embeddings.

A counterfactual traversal was generated for each base cell (seed) by displacing its semantic embedding z₀ along *d* while holding its stochastic latent fixed, and decoding each step with the diffusion model. Rather than drawing that latent at random, we recovered it by DDIM inversion (Denoising Diffusion Implicit Models^59^) of the base-cell image — integrating the deterministic diffusion ODE in reverse (data → noise) under the cell’s own semantic embedding and the same classifier-free guidance^60^ used for forward sampling (“guided inversion”), yielding the noise latent x_T. Because the conditional generator takes the semantic embedding and this residual context x_T separately, holding x_T fixed while z₀ is displaced regenerates the exact cell at zero displacement (pixel Pearson ≈ 0.99) and lets the traversal alter only the phenotype, preserving the cell’s context.

The amount of displacement needed to reach a knockout’s full phenotype varies from gene to gene, so we report displacement on a per-gene normalized axis φ. φ = 0 is the unperturbed control and φ = 1 is the point of maximal recovery of the real knockout phenotype; φ > 1 extrapolates beyond it. Recovery uses a nearest-centroid classification in Cell-DINO embedding space. The centroid for each knockout class is the mean of its top-predictive 1,000 real cells, z-scored per dimension and L2-normalized. At each displacement value, 200 cells for a gene knockout of interest were generated, and embedded using Cell-DINO. These displaced embeddings were standardized with a single per-dimension mean and standard deviation, computed once from generating zero-displacement images for the same 200 seed cells. At each displacement value, each generated cell was assigned to one of 1,000 knockout classes based on its cosine similarity to the nearest class centroid. The displacement at which this classification accuracy peaks defines the per-gene scale *f*. The normalized axis is φ = displacement / *f*.

Classical morphometric features were measured on both real and generated images to confirm the traversal produces the expected phenotypic effect in interpretable feature space. For three example knockouts spanning distinct organelles — TAF1B (total nucleolar area per cell), SAMM50 (mitochondrial fragment count per cell), and MTOR (total lysosome area per cell), each feature was measured on the top-1,000 predictive real NTC/KO cells and on 100 independently-traversed generated cells per φ, with a bar at the median and percent change reported relative to a single shared real-NTC baseline. Violin axes are limited to the 1st–98th percentile of each panel’s pooled data; densities and medians are computed from the full per-cell values.

---

**Supplementary Figure 1:**
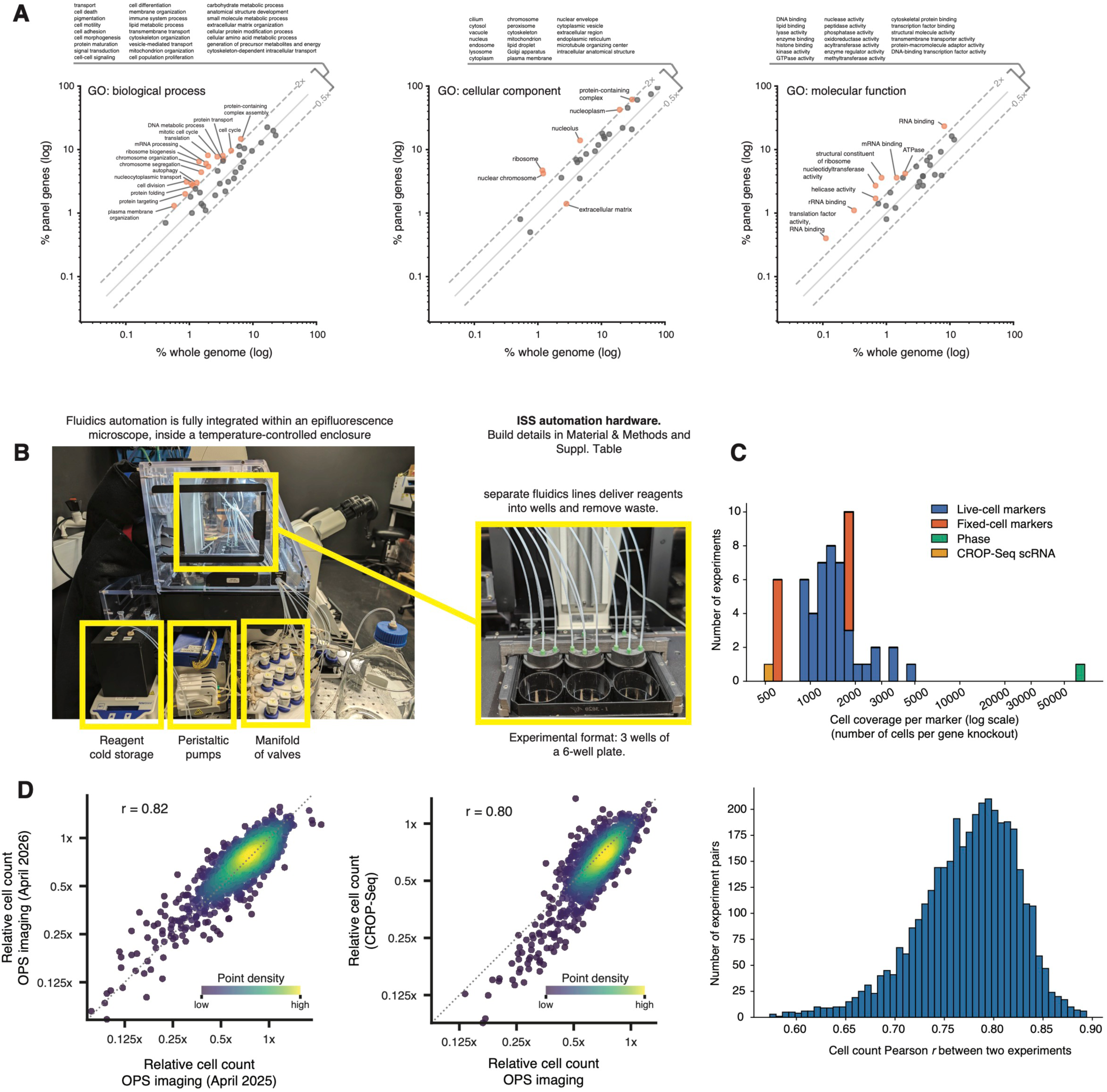
Screening library and deployment. **(A)** Gene Ontology annotation distribution of the 1,000-gene knockout library relative to the whole genome. Over- and under-represented terms are highlighted. While we ensured a balanced representation of different cellular pathways, core biological processes, such as gene expression, translation, and protein transport, were intentionally enriched in the library because they have been shown to generate rich phenotypic signatures in other screens. **(B)** Fluidics automation hardware for *in situ* sequencing. See Suppl. Table 6 for hardware bill of materials. **(C)** Cell coverage (number of cells per gene knockout) across the whole marker library. **(D)** The reproducibility of phenotypic penetrance between different screens was measured by comparing the cell count for individual gene knockouts (relative to NTC controls) between two experiments; that is, the similarity of essential gene depletion between these two screens. High reproducibility was observed between two OPS screens performed one year apart (left panel, Pearson r = 0.82), or between OPS and CROP-seq experiments (middle panel, Pearson r = 0.8). The full distribution of pairwise Pearson r values between all 88 screens in our dataset is shown on the right.

**Supplementary Figure 2:**
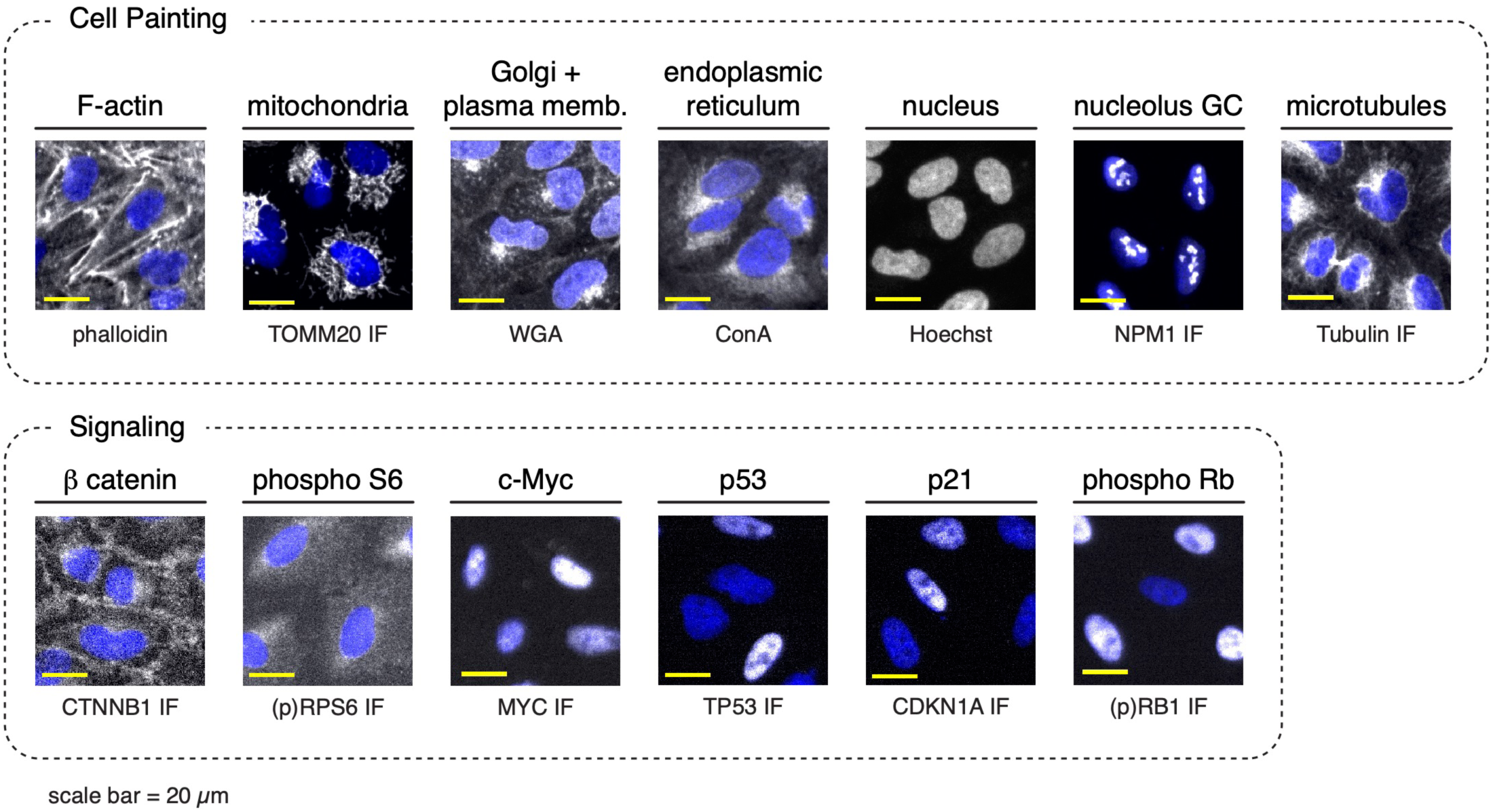
Example images of fixed cell markers. Marker signal is shown in grayscale, while nuclei are shown in blue.

**Supplementary Figure 3:**
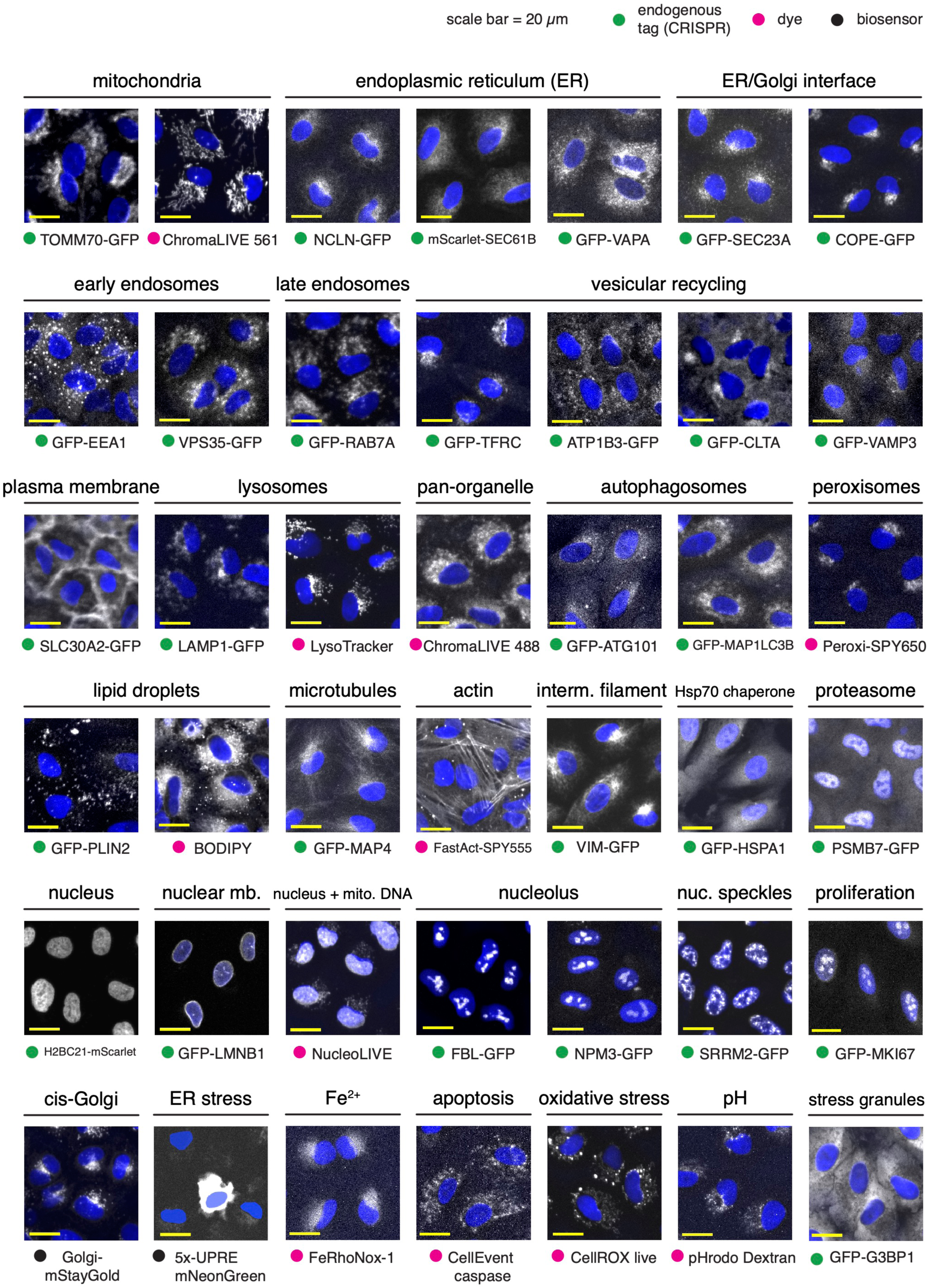
Example images of live cell markers. Marker signal is shown in grayscale, while nuclei are shown in blue.

**Supplementary Figure 4:**
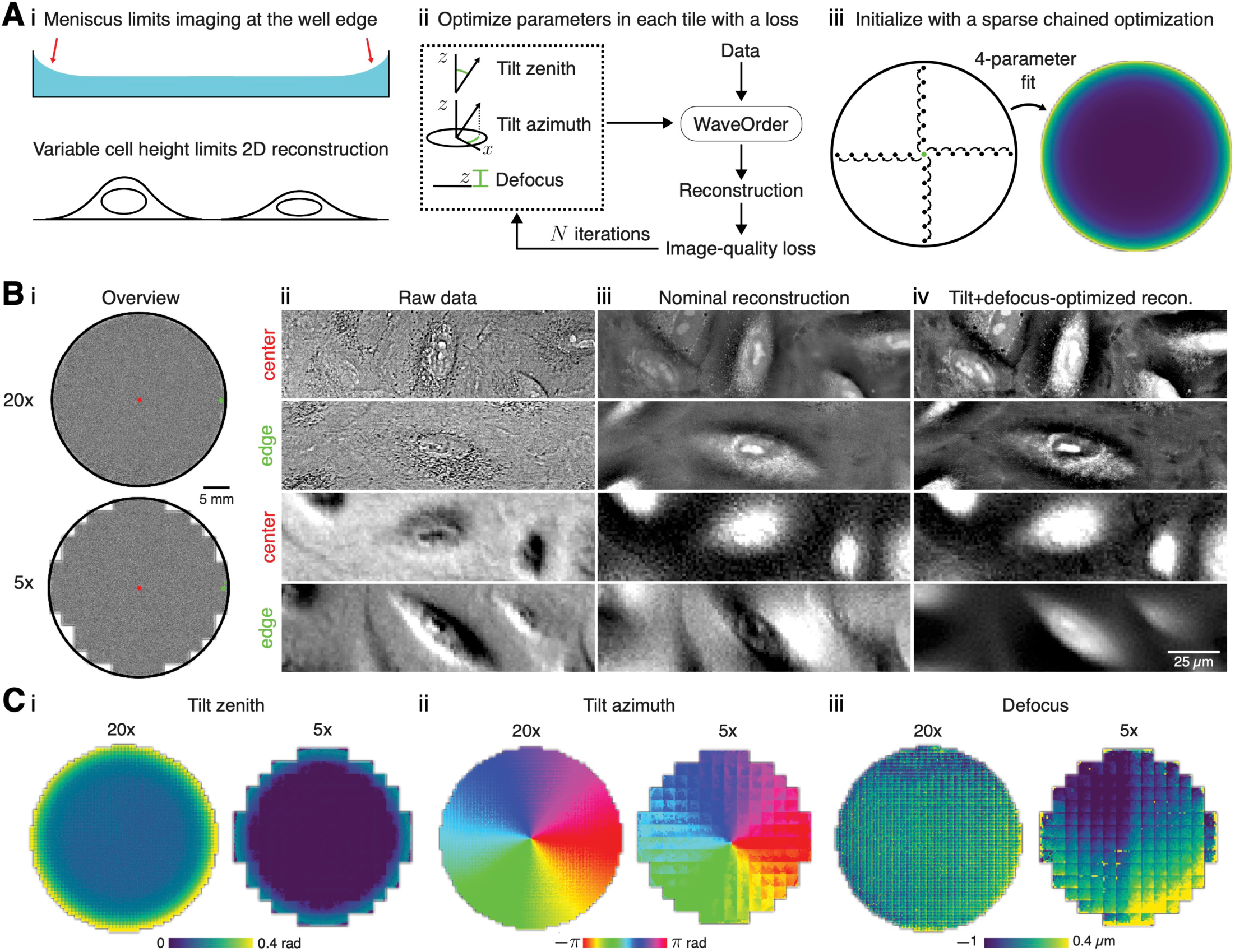
Quantitative phase reconstruction with WaveOrder. Phase reconstruction with tilt and defocus optimization enables label-free tracking and phenotyping across entire wells. **(A)** Label-free transmission imaging is impacted by **(i)** the air-media meniscus near well edges that distorts illumination light (top), and variable cell height and defocus (bottom), which limit how much information can be encoded in a single slice. We developed **(ii)** an iterated reconstruction approach where three scalar parameters, tilt zenith, tilt azimuth, and defocus, were estimated by optimizing an image-quality loss (mid-band spatial frequency). We found that our iterated scheme was sensitive to initial conditions at the well edge, so **(iii)** we developed a sparse chained optimization to find and fit good initial conditions. **(B)** Label-free transmission imaging with defocus across **(i)** the entire well at 20× (phenotyping, top) and 5× (tracking, bottom) generates **(ii)** raw data with well-position-dependent contrast; ROIs from the well center (first and third rows, red dot in **i**) show visibly different contrast from well edges (second and fourth rows, green dot in **i**). **(iii)** Phase-from-defocus reconstructions improve contrast, but perform poorly at the well edge, particularly in 5×. Our iterated approach resulted in **(iv)** consistent high-contrast reconstructions across the entire well together with (C) parameter maps that can help us understand the source of contrast changes across each well.

**Supplementary Figure 5:**
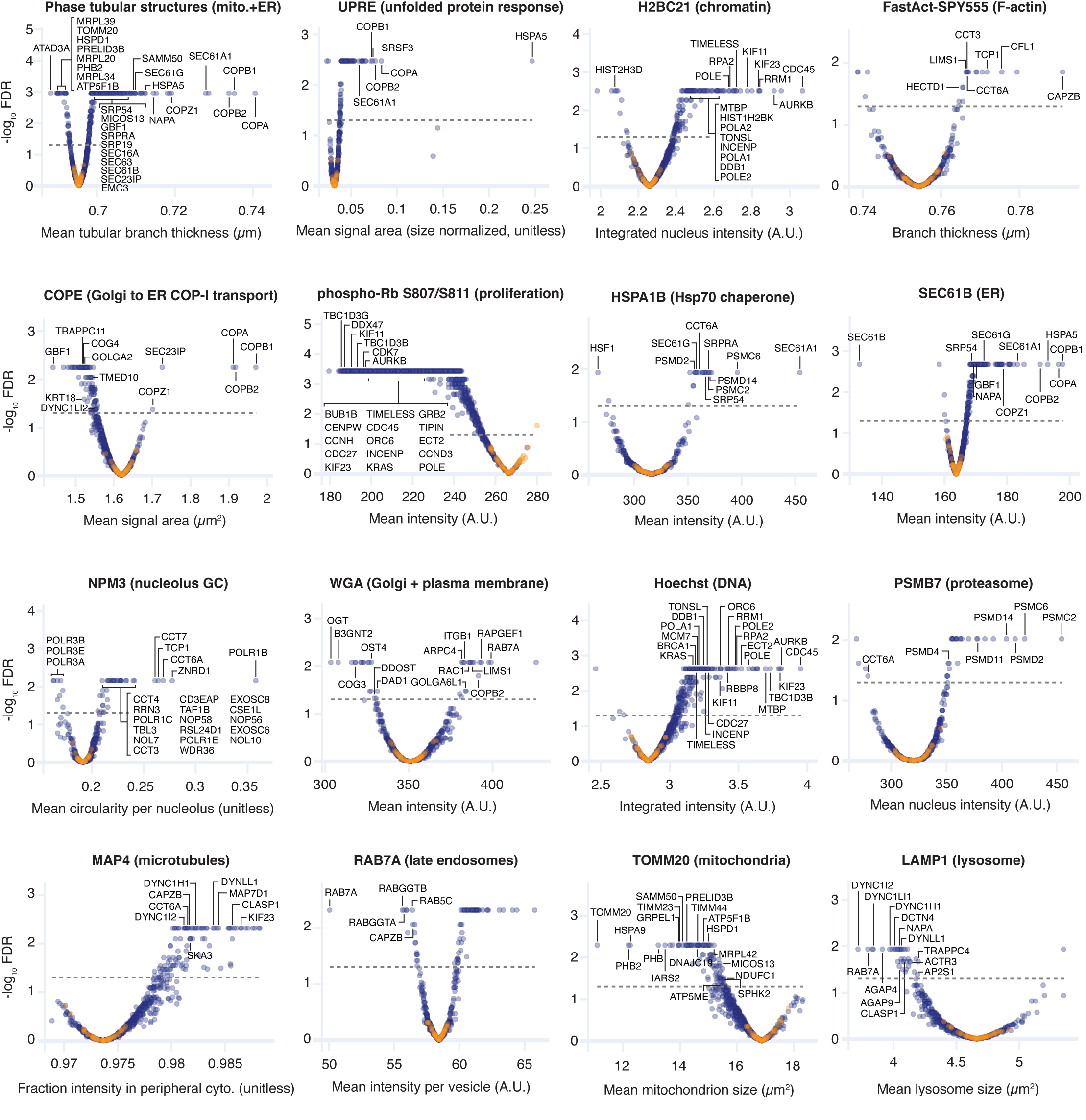
Volcano plots validate expected biology in our screens. Each symbol: median measurement for a KO, calculated from up to 5,000 random cells sampled from different biological replicate experiments. Orange symbols: non-targeting controls, blue symbols: KO perturbations. Genes known to be related to the biology represented by each marker are highlighted. Definition of the features being quantified and comprehensive plot values can be found in Suppl. Tables 12 and 13, respectively. See also Figures 1F-I.

**Supplementary Figure 6:**
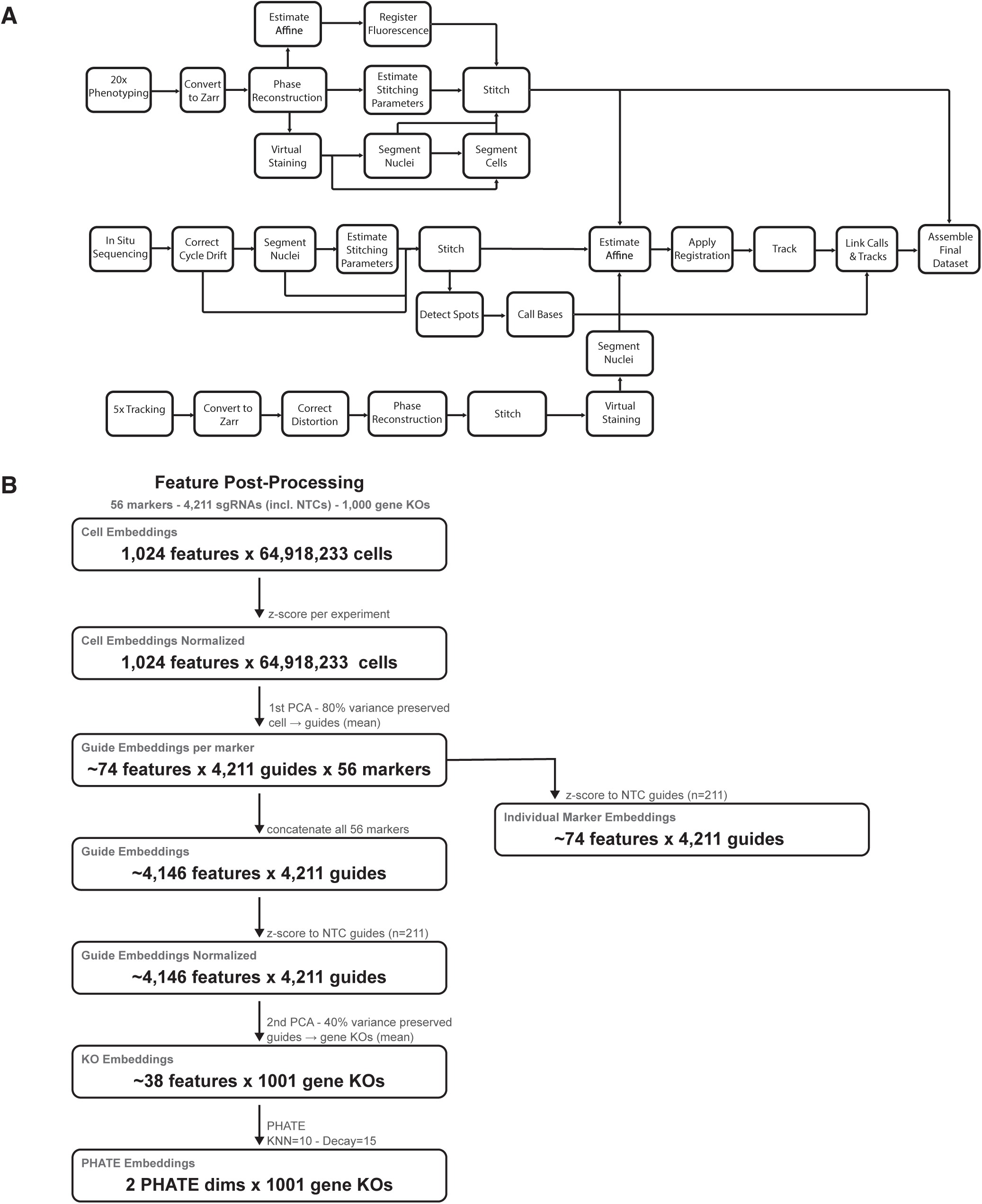
Image processing & embedding strategy. **(A)** Process flow diagram of the OPS image processing pipeline, highlighting major steps and their dependencies. Three co-acquired imaging modalities are processed for each experiment: 20× confocal phenotyping (volumetric brightfield and fluorescence, 7 z-slices), 5× live-cell tracking (single-plane brightfield), and 5× fluorescence multi-cycle *in situ* sequencing (ISS). Preprocessing is run independently for each modality, before they are co-registered and assembled into the final dataset. **(B)** Embeddings from Cell-DINO (and all other benchmarked models) are post-processed to remove batch effects or other sources of noise, and combine information from many different markers into a compact final embedding. PCA dimensionality reduction is applied twice, once at the single-cell level, per marker to reduce batch effects, and again at the combined marker embedding level, to minimize the influence of noisy or low information markers.

**Supplementary Figure 7:**
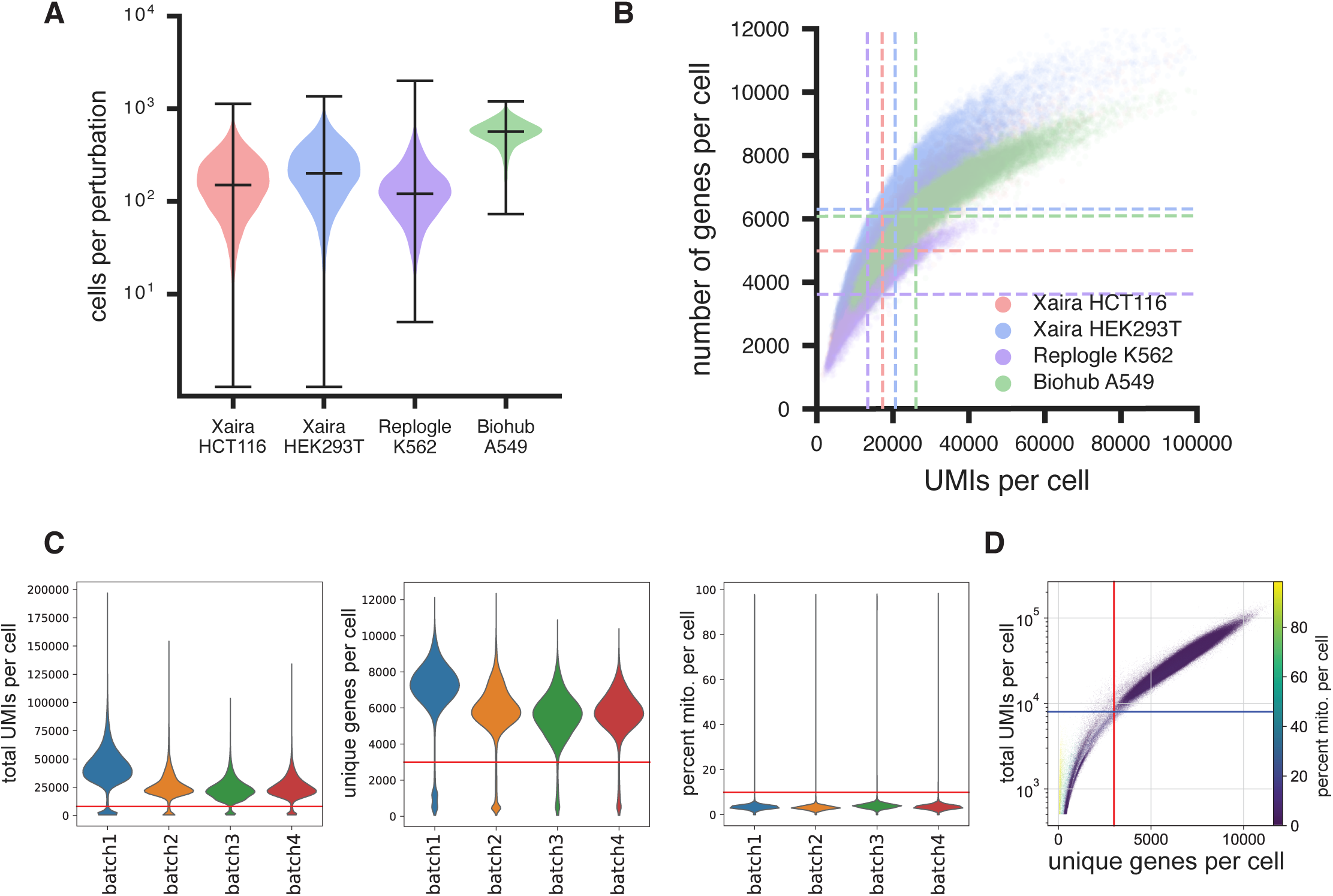
CROP-seq metrics. Comparison of sequencing depth to recently published CROP-seq datasets. **(A)** Violin plots of the number of cells per perturbation for datasets from Huang et al.^75^ (Xaira HCT116, Xaira HEK293T), Replogle et al.^11^ (Replogle K562) and the dataset from this publication (Biohub A549). Cell numbers for different guides targeting the same gene were summed. The Biohub dataset sequences more cells per perturbed gene on average. Note however that it covers 1,000 gene knockouts compared to the genome-wide coverage of the comparison datasets. **(B)** Scatterplot of unique molecular identifiers (UMIs) per cell versus number of genes with at least one UMI. The Biohub dataset has the deepest average sequencing depth per cell. Biohub A549 point cloud overlaps with Xaira HCT116; dotted lines mark average values for each dataset. **(C)** Violin plots showing QC metrics by batch for all 10X droplet sequencing experiments (our 1,000 knockout library was processed across four separate experimental batches). Note that batch 1 was re-sequenced for a total of ∼twice the read depth of all other batches. Left: Total UMIs per cell. Middle: Unique genes per cell. Right: Percent mitochondrial UMIs per cell. Red lines show quality thresholds; all droplets falling below these thresholds are excluded from downstream analysis (except for mitochondrial UMIs per cell, for which droplets exceeding the threshold are excluded). **(D)** Scatter plot showing relationship between QC metrics in (C). Only cells in the upper right quadrant were kept for all analyses shown in this paper.

**Supplementary Figure 8:**
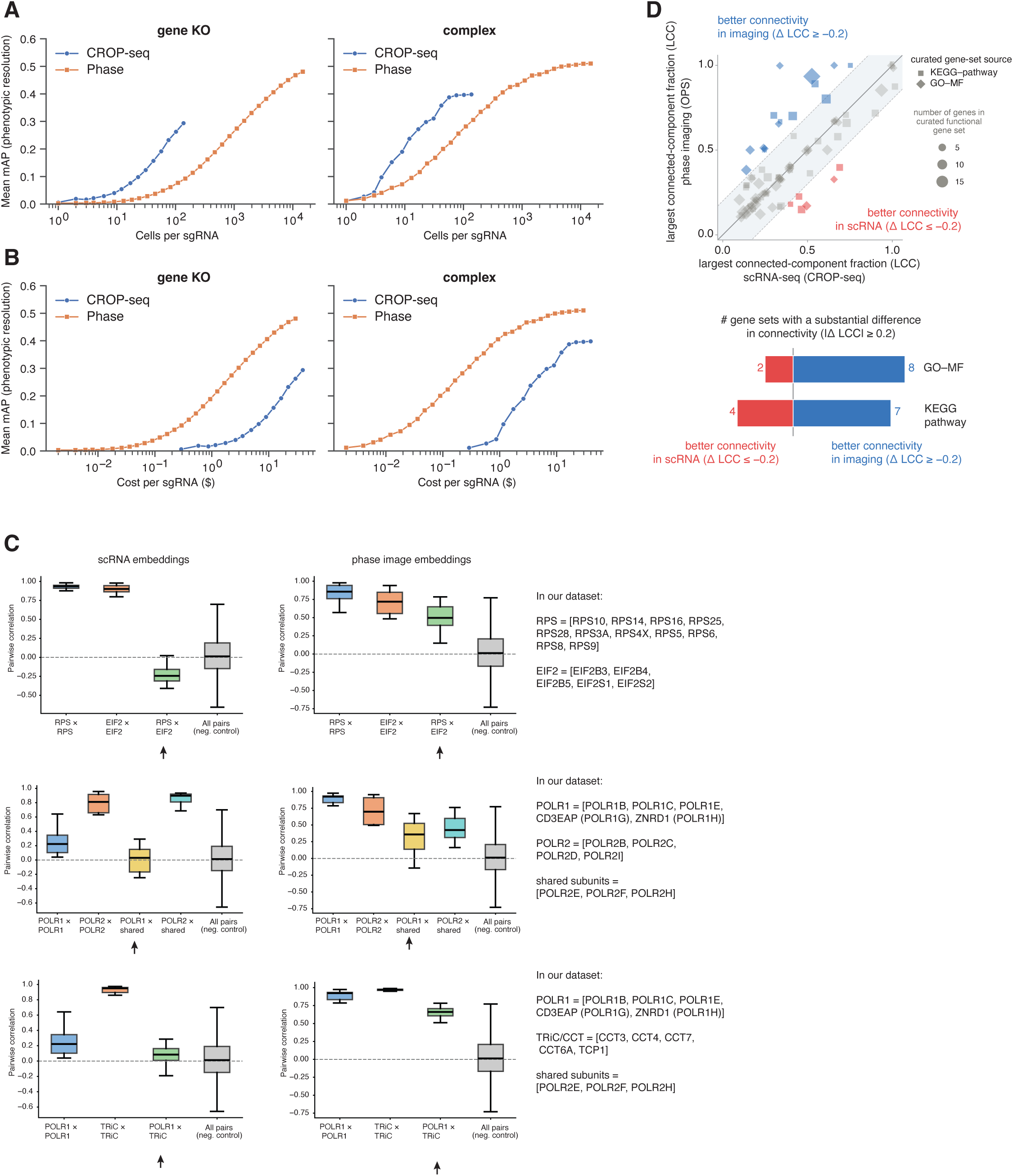
Quantitative comparison of scRNA vs. phase imaging phenotypic resolution. **(A)** mAP scores for CROP-seq and Phase imaging OPS data at varying cells-per-sgRNA coverage. Left: gene-KO mAP; right: protein-complex mAP. **(B)** mAP scores for CROP-seq and Phase imaging OPS data at varying $ cost-per-sgRNA coverage. Left: gene-KO mAP; right: protein-complex mAP. **(C)** Distribution of pairwise embedding correlations within and across subunits of (i) the eIF2 and RPS protein complexes (top); (ii) the Pol I, Pol II protein complexes, and their shared subunits (middle); (iii) the TRiC/CCT and Pol I protein complexes. **(D)** For each gene set in an ontology, we computed the Largest Connected Component fraction (LCC). LCC measures the largest number of gene set members that are connected to at least one other member in the k-NN graph, relative to total set size. Values are shown for phase imaging OPS versus scRNA-seq k-NN graphs. A difference in connectivity was considered substantial when |ΔLCC| > 0.2 between the two modalities. Phase imaging showed higher LCC overall (paired Wilcoxon test, n = 107 gene sets, p = 0.051).

**Supplementary Figure 9:**
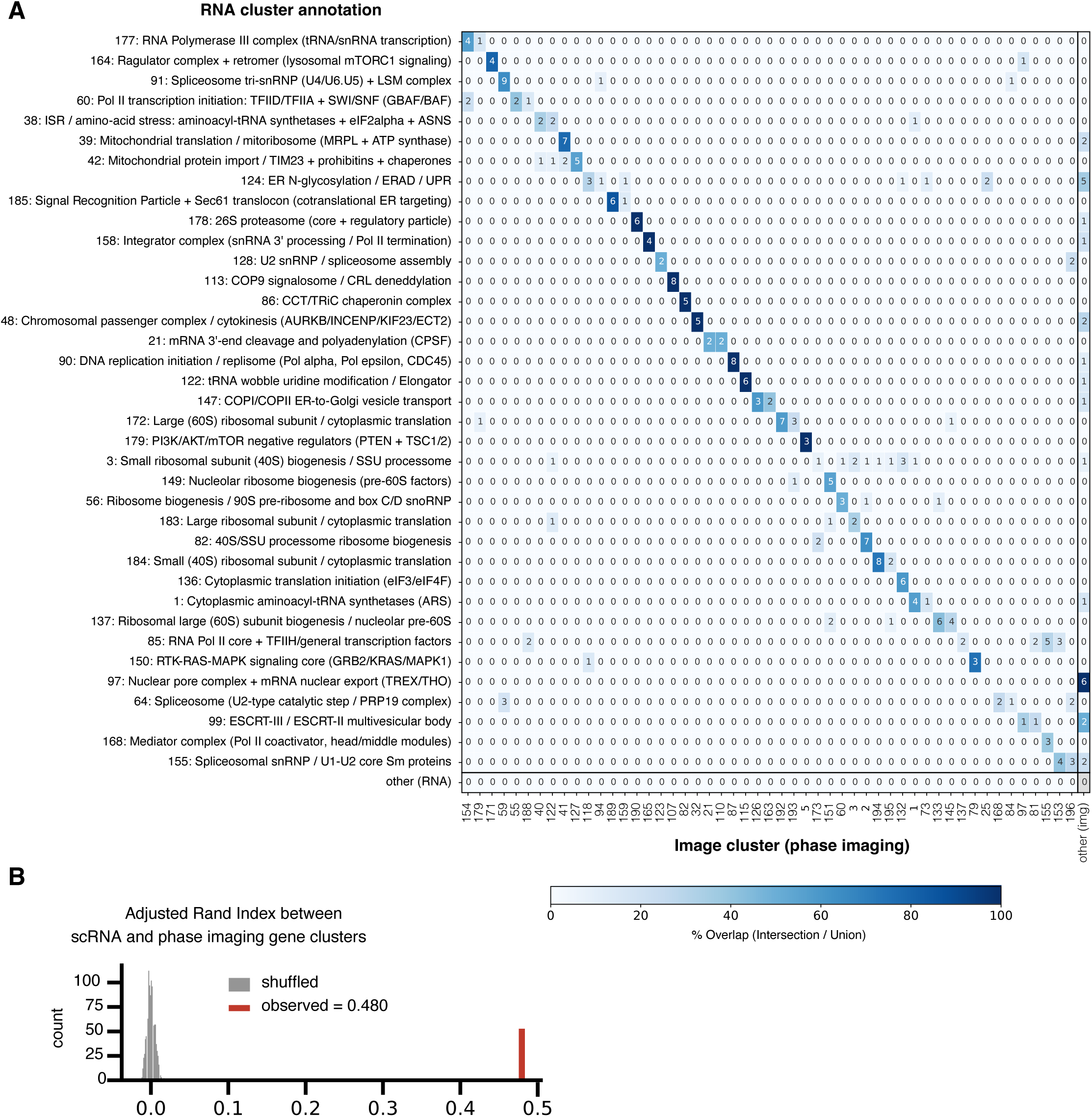
Cluster comparison in CROP-seq vs. phase image embeddings. Unsupervised biological insights from CROP-seq screens can largely be replicated with optical pooled screens. **(A)** Confusion matrix comparing unsupervised Leiden clustering of the RNA modality to the Image modality. Numbers indicate the overlapping perturbed genes and colors indicate the percent of common genes (intersection) over the total number of genes in both clusters (union). RNA sVAE+ embeddings were clustered with Leiden (resolution = 29) partitioning perturbations into 186 clusters of similar transcriptional response. These were enriched against gene ontology libraries, and for each cluster a final annotation and confidence score was obtained by summarization with a large language model. Only 37 clusters with a confidence score of 5 were considered here and are shown on the y-axis. The x-axis shows unsupervised Leiden clusters (a separate Leiden clustering at resolution 28 was used here for the best comparison) of PCA-reduced image embeddings with an overlap of at least 2 genes with an RNA cluster (additional clusters with smaller overlap are collected in the bottom and right margin columns). Rows and columns were paired on the diagonal by maximum intersection over union, then the paired block was reordered jointly by hierarchical clustering. **(B)** The subset of scRNA high-resolution Leiden clusters (resolution = 29) that displayed significant enrichment for gene ontology terms was identified (i.e. the clusters with clearest biological coherence, n = 37 clusters). The Adjusted Rand Index between these scRNA clusters and corresponding Leiden clusters in phase image space was calculated (red horizontal line). The null distribution for 1,000 permuted cluster labels is shown as a grey histogram, indicating that both modalities largely agree in grouping functionally similar perturbations.

