## Supplementary material for "A multimodal perturbation atlas defines the phenotypic resolution of cellular morphology": Data S1

### Fully annotated version of Figure 2H

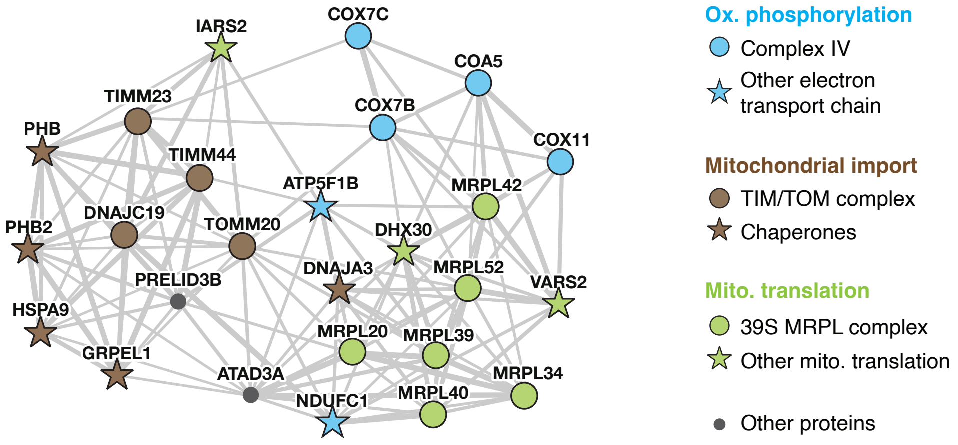


### Fully annotated version of Figure 2I

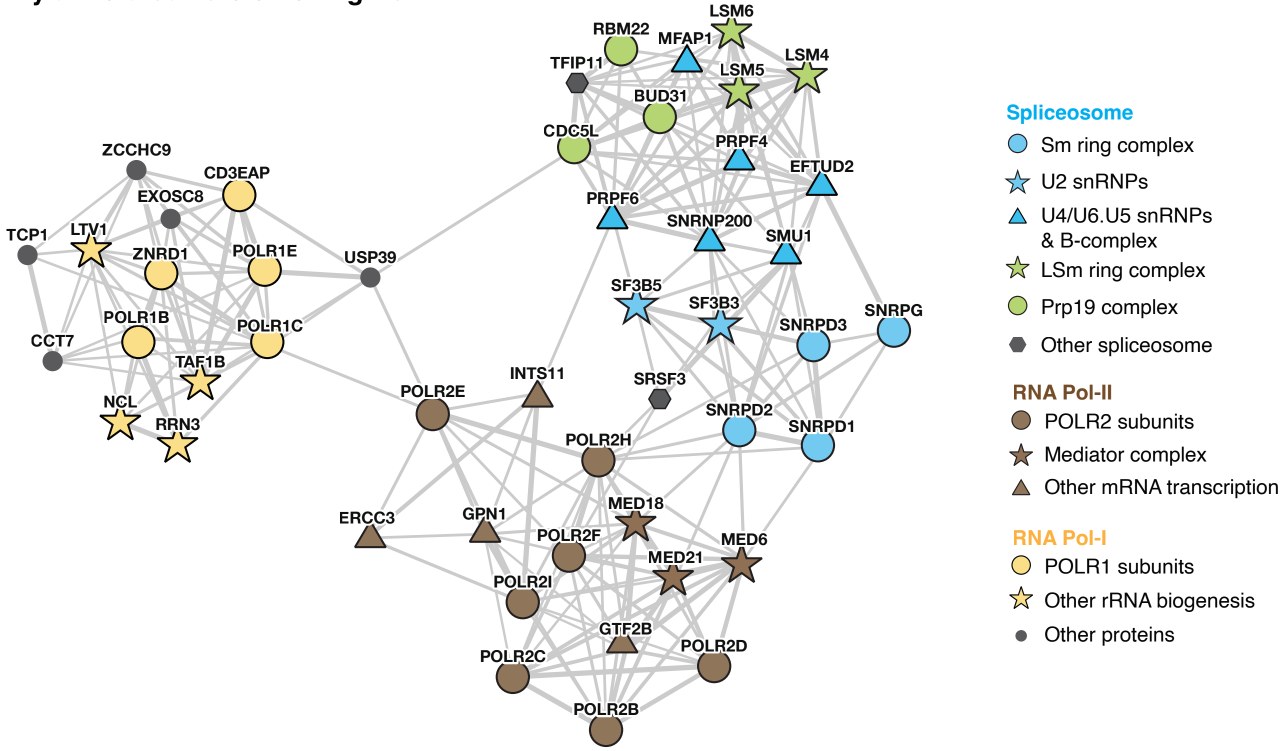

### Fully annotated version of Figure 4C

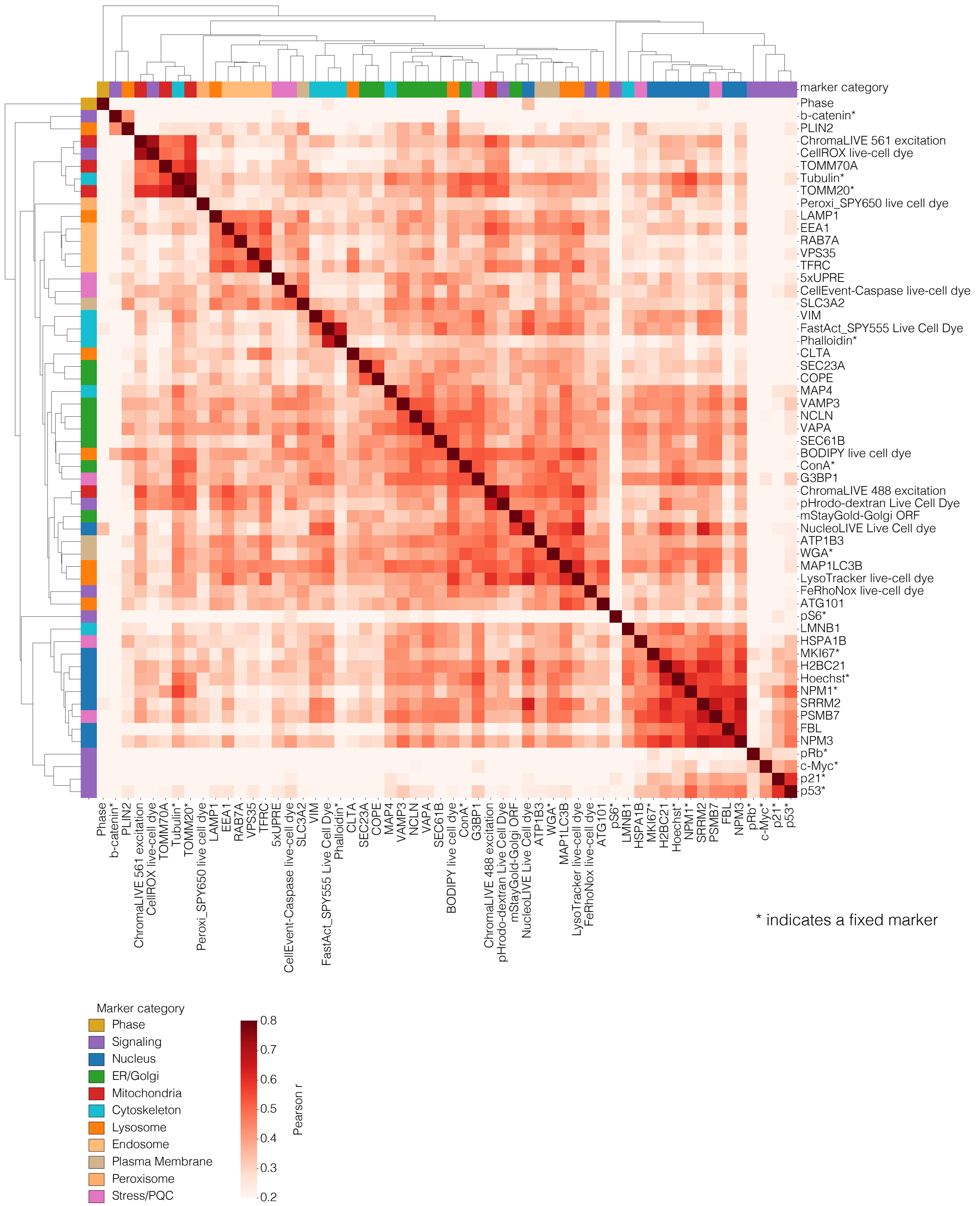

**Fully annotated version of Figure 5 panels**

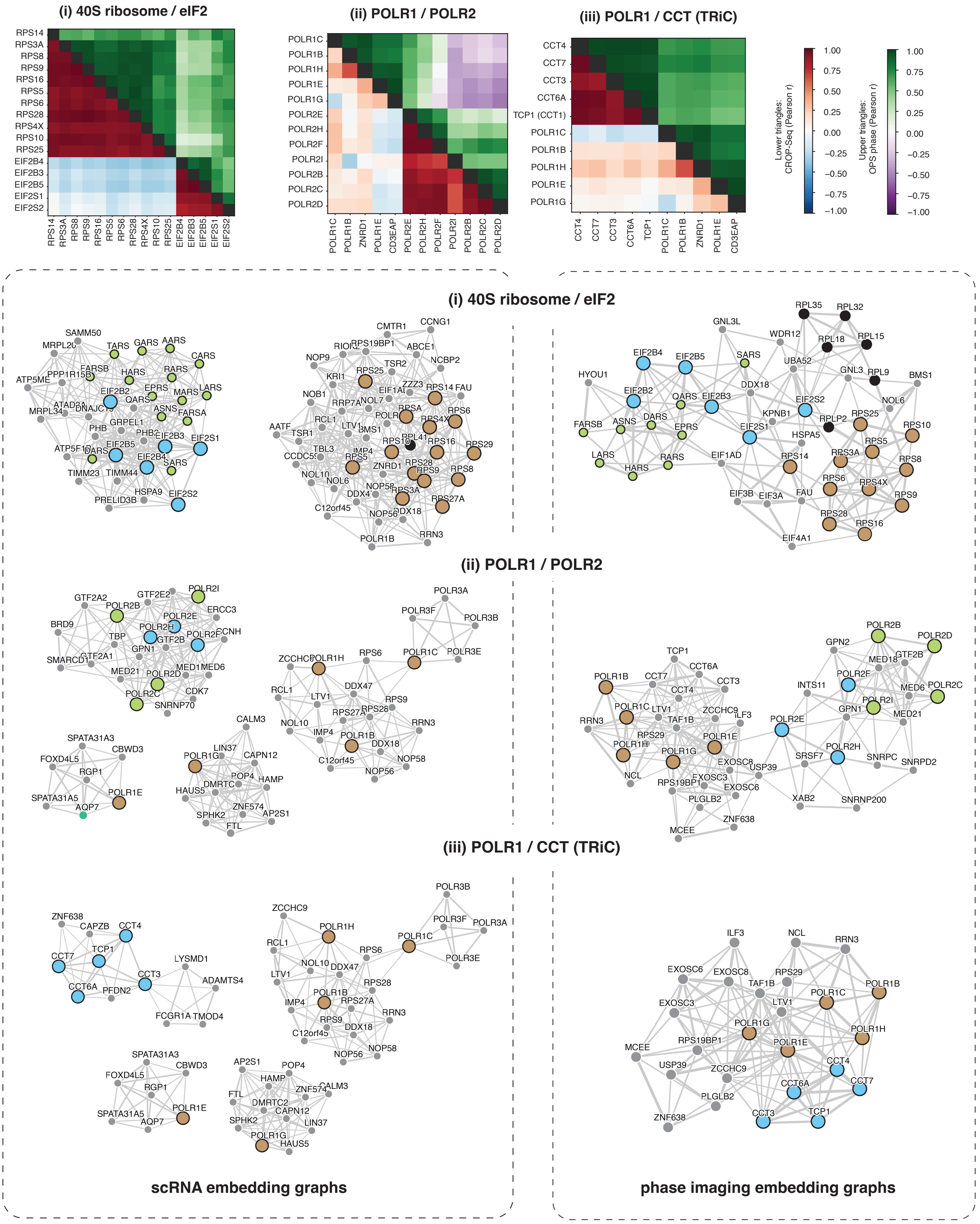
